## Supplementary Figures for "Accounting for differences between Infinium MethylationEPIC v2 and v1 in DNA methylation-based tools"

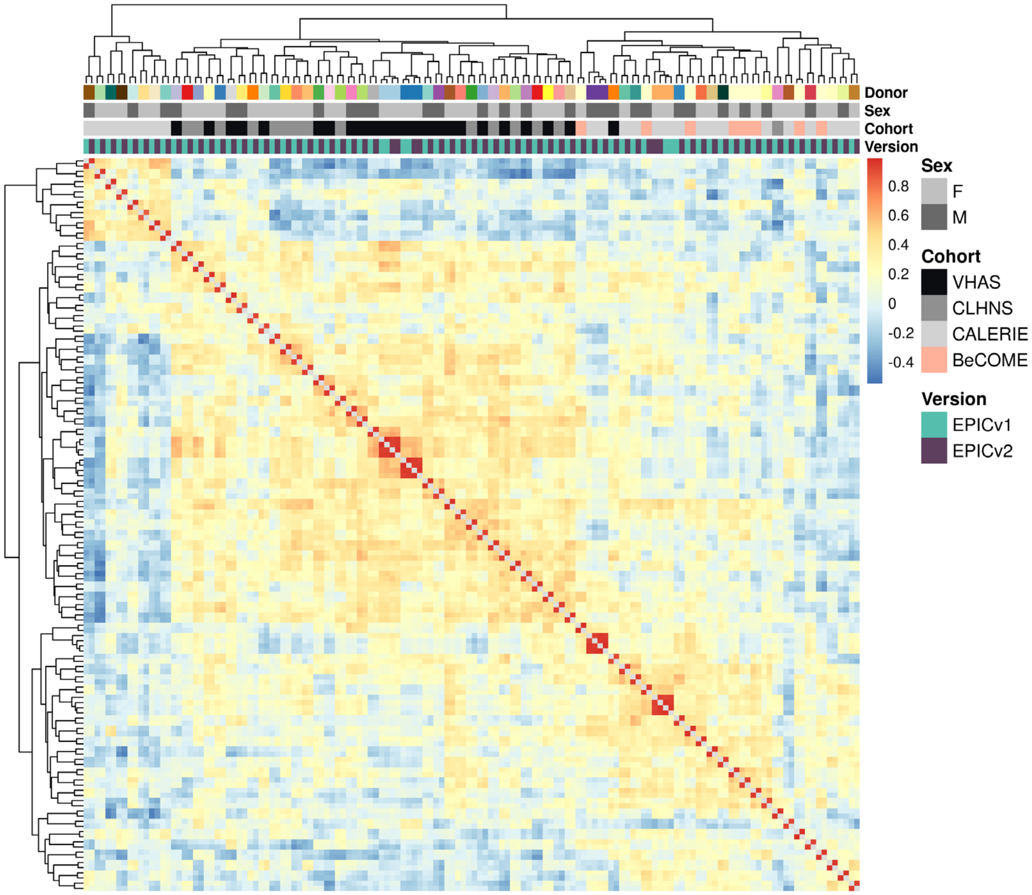


**Supplementary Figure 1.** Unsupervised hierarchical clustering using complete linkage with Euclidean distance on sample-to-sample Spearman correlations, calculated using the 57 SNP probes shared between EPICv1 and EPICv2 with raw data; blue to red color range denotes Spearman *rho* correlation from low to high.

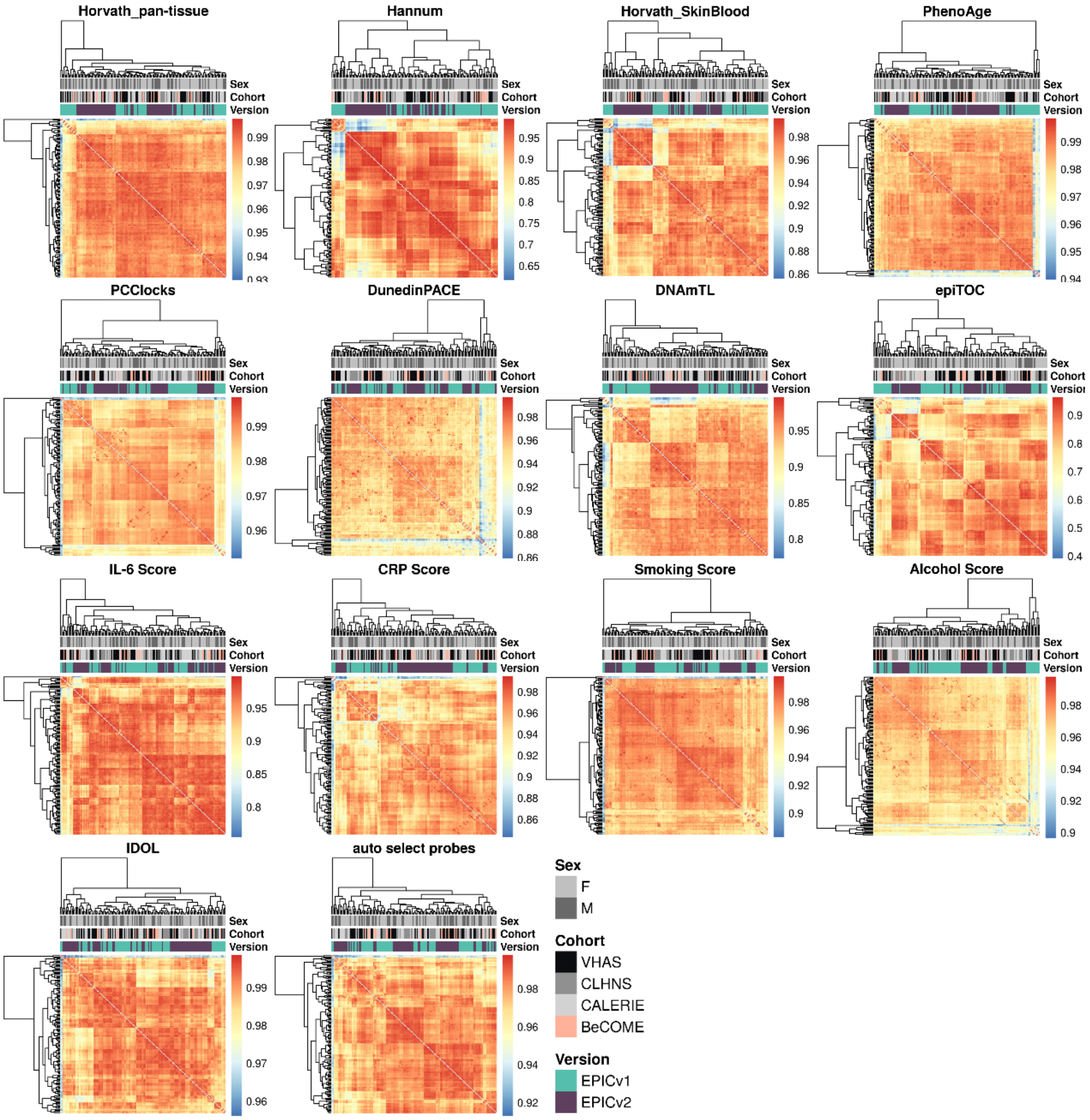


**Supplementary Figure 2.** Unsupervised hierarchical clustering using complete linkage with Euclidean distance on sample-to-sample Pearson correlations, calculated using predictive CpGs employed by clocks, biomarker predictors, and cell type deconvolution algorithms shared between EPICv1 and EPICv2 with functional normalized data; blue to red color range denotes Pearson correlation from low to high.

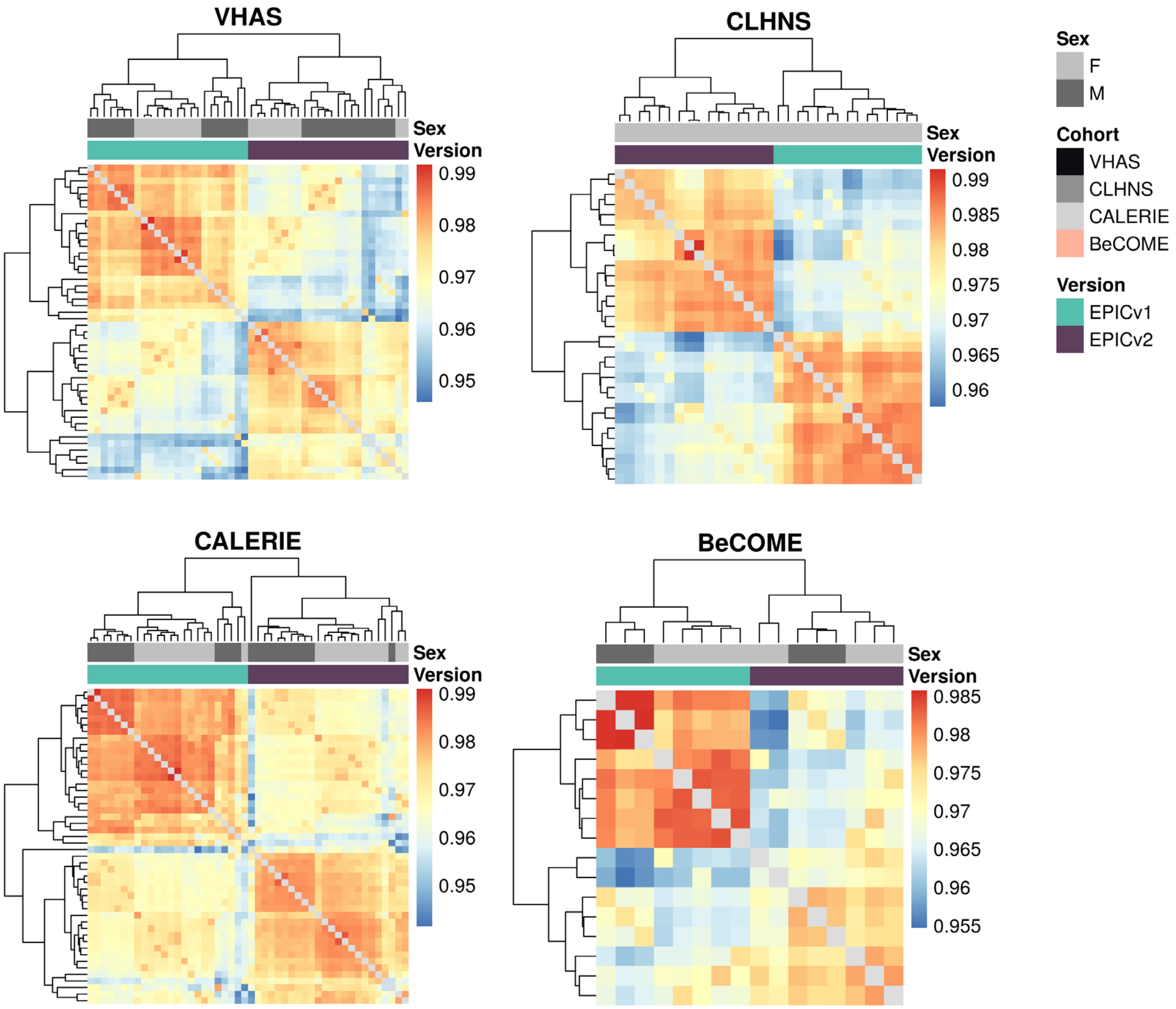


**Supplementary Figure 3.** Unsupervised hierarchical clustering using complete linkage with Euclidean distance on sample-to-sample Spearman correlations, calculated using the 721,378 probes shared between EPICv1 and EPICv2 with functional normalized data for each cohort; blue to red color range denotes Spearman *rho* correlation from low to high.

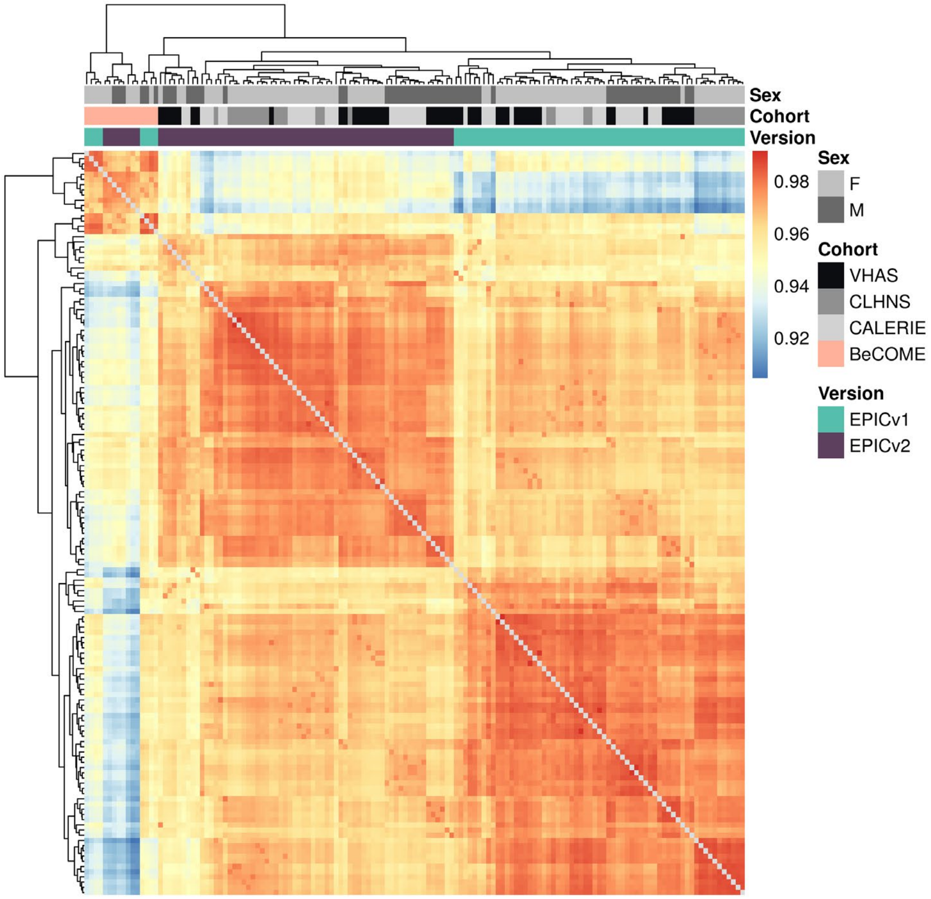


**Supplementary Figure 4.** Unsupervised hierarchical clustering using complete linkage with Euclidean distance on sample-to-sample Spearman correlations, calculated using the 721,378 probes shared between EPICv1 and EPICv2 with functional normalized data with four cohorts combined; blue to red color range denotes Spearman *rho* correlation from low to high.

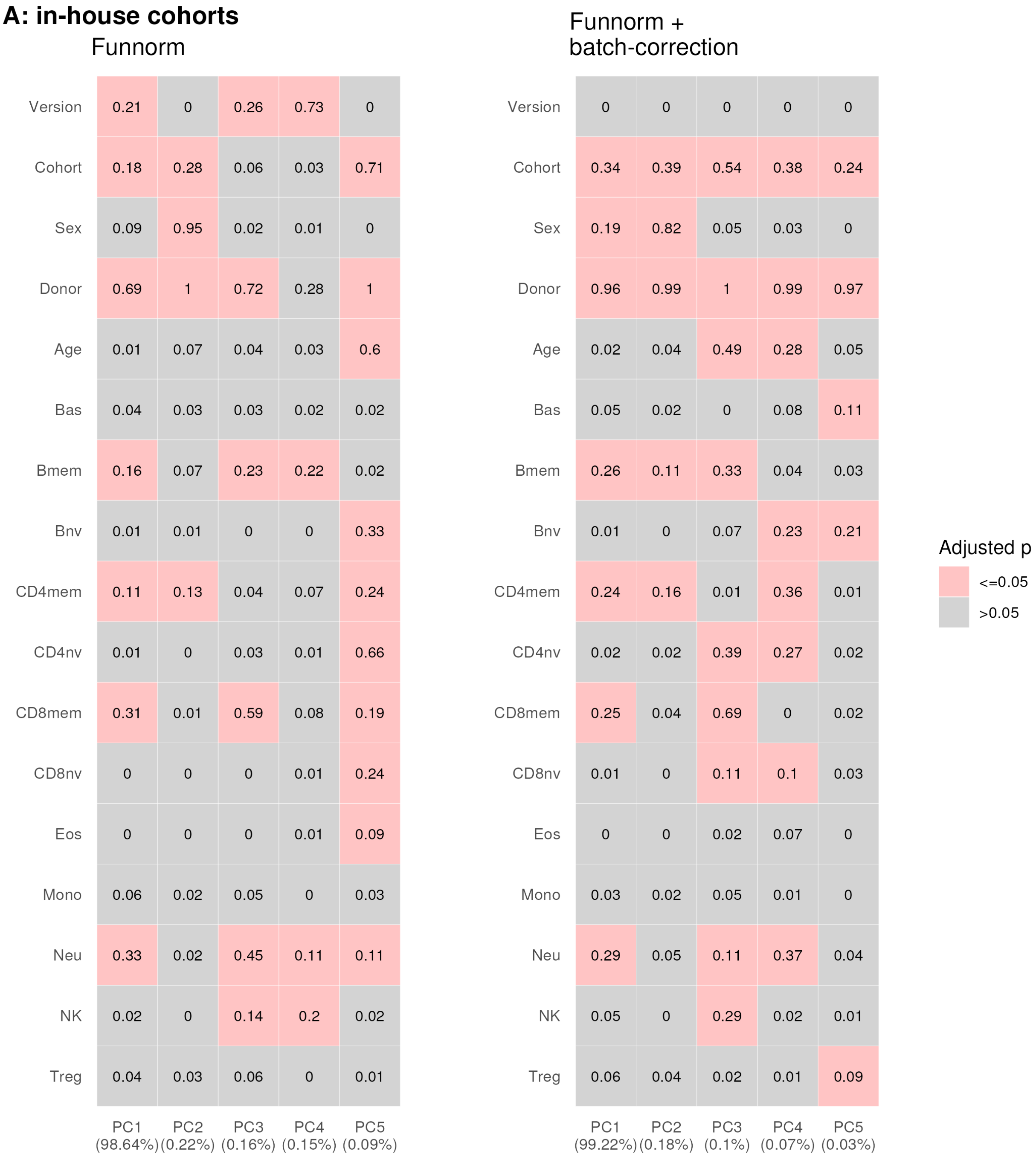
A: in-house cohorts
Funnorm
Version 0.21 0 0.26 0.73 0
Cohort 0.18 0.28 0.06 0.03 0.71
Sex 0.09 0.95 0.02 0.01 0
Donor 0.69 1 0.72 0.28 1
Age 0.01 0.07 0.04 0.03 0.6
Bas 0.04 0.03 0.03 0.02 0.02
Bmem 0.16 0.07 0.23 0.22 0.02
Bnv 0.01 0.01 0 0 0.33
CD4mem 0.11 0.13 0.04 0.07 0.24
CD4nv 0.01 0 0.03 0.01 0.66
CD8mem 0.31 0.01 0.59 0.08 0.19
CD8nv 0 0 0 0.01 0.24
Eos 0 0 0 0.01 0.09
Mono 0.06 0.02 0.05 0 0.03
Neu 0.33 0.02 0.45 0.11 0.11
NK 0.02 0 0.14 0.2 0.02
Treg 0.04 0.03 0.06 0 0.01
PC1 (98.64%) PC2 (0.22%) PC3 (0.16%) PC4 (0.15%) PC5 (0.09%)
Funnorm + batch-correction
Version 0 0 0 0 0
Cohort 0.34 0.39 0.54 0.38 0.24
Sex 0.19 0.82 0.05 0.03 0
Donor 0.96 0.99 1 0.99 0.97
Age 0.02 0.04 0.49 0.28 0.05
Bas 0.05 0.02 0 0.08 0.11
Bmem 0.26 0.11 0.33 0.04 0.03
Bnv 0.01 0 0.07 0.23 0.21
CD4mem 0.24 0.16 0.01 0.36 0.01
CD4nv 0.02 0.02 0.39 0.27 0.02
CD8mem 0.25 0.04 0.69 0 0.02
CD8nv 0.01 0 0.11 0.1 0.03
Eos 0 0 0.02 0.07 0
Mono 0.03 0.02 0.05 0.01 0
Neu 0.29 0.05 0.11 0.37 0.04
NK 0.05 0 0.29 0.02 0.01
Treg 0.06 0.04 0.02 0.01 0.09
PC1 (99.22%) PC2 (0.18%) PC3 (0.1%) PC4 (0.07%) PC5 (0.03%)
Adjusted p
<=0.05
>0.05

**B: i.VHAS**

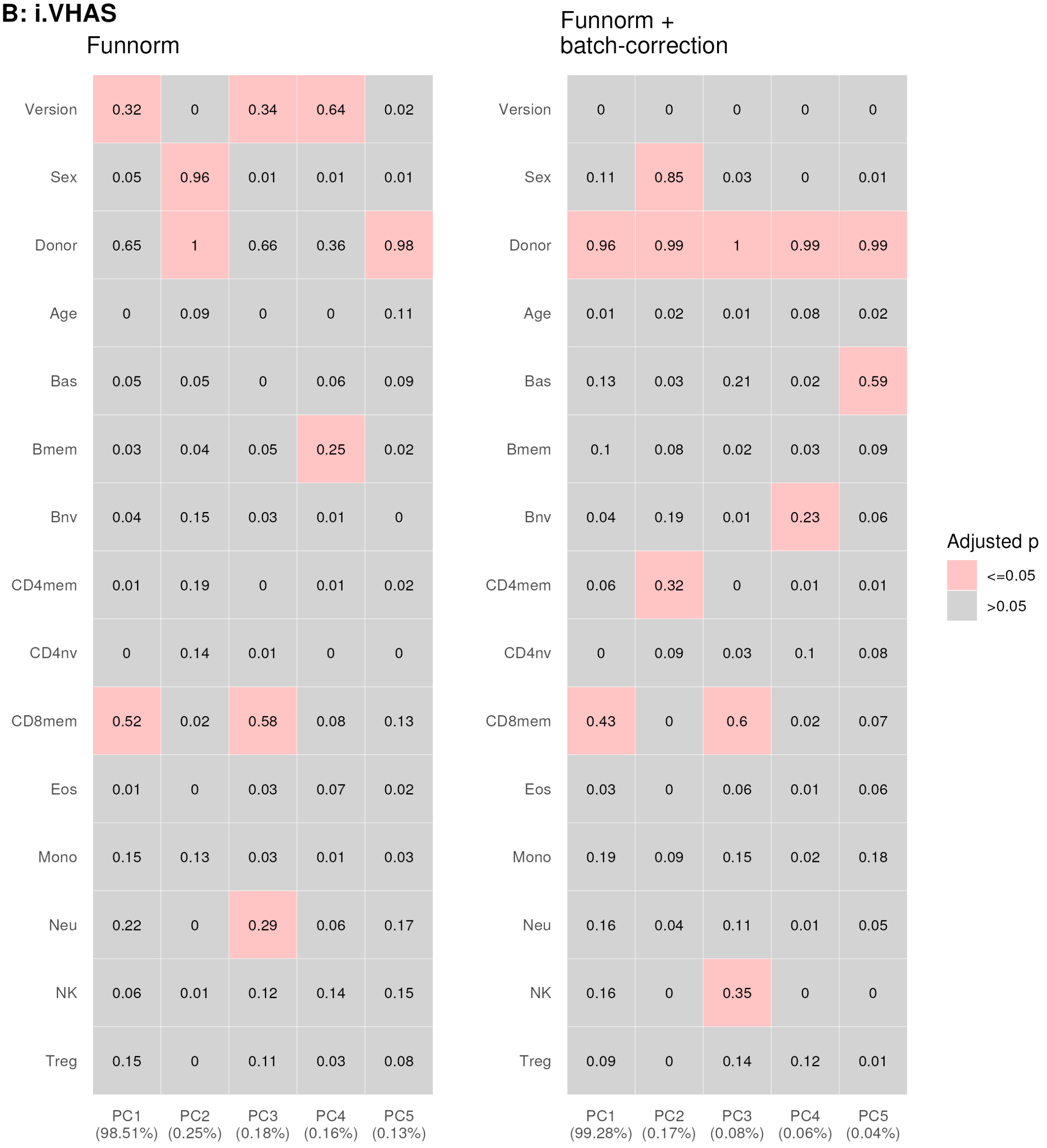
Funnorm
Funnorm + batch-correction
Version 0.32 0 0.34 0.64 0.02
Sex 0.05 0.96 0.01 0.01 0.01
Donor 0.65 1 0.66 0.36 0.98
Age 0 0.09 0 0 0.11
Bas 0.05 0.05 0 0.06 0.09
Bmem 0.03 0.04 0.05 0.25 0.02
Bnv 0.04 0.15 0.03 0.01 0
CD4mem 0.01 0.19 0 0.01 0.02
CD4nv 0 0.14 0.01 0 0
CD8mem 0.52 0.02 0.58 0.08 0.13
Eos 0.01 0 0.03 0.07 0.02
Mono 0.15 0.13 0.03 0.01 0.03
Neu 0.22 0 0.29 0.06 0.17
NK 0.06 0.01 0.12 0.14 0.15
Treg 0.15 0 0.11 0.03 0.08
PC1 (98.51%) PC2 (0.25%) PC3 (0.18%) PC4 (0.16%) PC5 (0.13%)
Version 0 0 0 0 0
Sex 0.11 0.85 0.03 0 0.01
Donor 0.96 0.99 1 0.99 0.99
Age 0.01 0.02 0.01 0.08 0.02
Bas 0.13 0.03 0.21 0.02 0.59
Bmem 0.1 0.08 0.02 0.03 0.09
Bnv 0.04 0.19 0.01 0.23 0.06
CD4mem 0.06 0.32 0 0.01 0.01
CD4nv 0 0.09 0.03 0.1 0.08
CD8mem 0.43 0 0.6 0.02 0.07
Eos 0.03 0 0.06 0.01 0.06
Mono 0.19 0.09 0.15 0.02 0.18
Neu 0.16 0.04 0.11 0.01 0.05
NK 0.16 0 0.35 0 0
Treg 0.09 0 0.14 0.12 0.01
PC1 (99.28%) PC2 (0.17%) PC3 (0.08%) PC4 (0.06%) PC5 (0.04%)
Adjusted p
<=0.05
>0.05

### B: ii.CLHNS

#### Funnorm

| | PC1 (99.08%) | PC2 (0.22%) | PC3 (0.19%) | PC4 (0.04%) | PC5 (0.04%) |
|---|---|---|---|---|---|
| Version | **0.52** | 0 | **1** | 0 | 0 |
| Donor | 0.42 | **1** | 0.02 | **0.99** | **0.99** |
| Age | 0.04 | 0.04 | 0 | 0.06 | 0.04 |
| Bas | 0.15 | 0.12 | 0.07 | 0.07 | **0.35** |
| Bmem | 0.23 | **0.74** | 0.01 | 0 | 0 |
| Bnv | 0.01 | 0.18 | 0.03 | 0.11 | 0.01 |
| CD4mem | 0.09 | **0.63** | 0.07 | 0.01 | 0 |
| CD4nv | 0.11 | 0.18 | 0 | **0.62** | 0.01 |
| CD8mem | **0.56** | **0.72** | 0.05 | 0.07 | 0 |
| Eos | 0.05 | 0 | 0.14 | 0 | 0.15 |
| Mono | 0.07 | 0 | 0.1 | 0.2 | 0.06 |
| Neu | **0.4** | **0.83** | 0 | 0 | 0.04 |
| NK | 0.01 | 0.17 | 0.04 | 0.14 | 0 |
| Treg | 0 | 0.21 | 0.12 | 0.1 | 0 |

#### Funnorm + batch-correction

| | PC1 (99.32%) | PC2 (0.2%) | PC3 (0.04%) | PC4 (0.04%) | PC5 (0.03%) |
|---|---|---|---|---|---|
| Version | 0.01 | 0 | 0 | 0 | 0 |
| Donor | **0.94** | **1** | **0.99** | **0.99** | **1** |
| Age | 0.1 | 0.07 | 0.06 | 0.08 | **0.39** |
| Bas | 0.09 | 0.13 | 0.07 | **0.37** | 0.1 |
| Bmem | **0.57** | **0.7** | 0 | 0 | 0.01 |
| Bnv | 0.09 | 0.15 | 0.12 | 0.02 | 0.07 |
| CD4mem | **0.44** | **0.58** | 0.01 | 0 | 0.19 |
| CD4nv | 0.21 | 0.16 | **0.63** | 0.02 | 0.02 |
| CD8mem | **0.75** | **0.74** | 0.07 | 0 | 0.01 |
| Eos | 0 | 0 | 0 | 0.14 | 0.28 |
| Mono | 0.01 | 0.01 | 0.2 | 0.07 | 0 |
| Neu | **0.72** | **0.82** | 0 | 0.03 | 0.05 |
| NK | 0.16 | 0.21 | 0.14 | 0.01 | 0.32 |
| Treg | 0.08 | 0.18 | 0.1 | 0 | 0.02 |

Adjusted p
- <=0.05
- >0.05

**B: iii.CALERIE**

Funnorm

| | PC1 (98.91%) | PC2 (0.24%) | PC3 (0.14%) | PC4 (0.12%) | PC5 (0.06%) |
|---|---|---|---|---|---|
| Version | 0.07 | 0 | 0.99 | 0 | 0 |
| Sex | 0.03 | 0.98 | 0 | 0.02 | 0 |
| Donor | 0.85 | 1 | 0 | 1 | 0.99 |
| Age | 0.01 | 0.05 | 0 | 0 | 0.01 |
| Bas | 0 | 0 | 0.11 | 0 | 0.44 |
| Bmem | 0.13 | 0.02 | 0.02 | 0.35 | 0.07 |
| Bnv | 0.02 | 0.02 | 0 | 0.06 | 0.15 |
| CD4mem | 0.19 | 0 | 0.02 | 0.34 | 0 |
| CD4nv | 0.01 | 0.09 | 0 | 0.13 | 0 |
| CD8mem | 0.23 | 0.07 | 0.07 | 0.26 | 0.18 |
| CD8nv | 0 | 0 | 0 | 0.09 | 0.06 |
| Eos | 0.25 | 0.09 | 0.04 | 0.08 | 0.42 |
| Mono | 0.08 | 0 | 0.02 | 0.05 | 0 |
| Neu | 0.33 | 0.15 | 0 | 0.74 | 0 |
| NK | 0.11 | 0.03 | 0.07 | 0.29 | 0.01 |
| Treg | 0 | 0.02 | 0.05 | 0.02 | 0.15 |

Funnorm + batch-correction

| | PC1 (99.52%) | PC2 (0.17%) | PC3 (0.05%) | PC4 (0.03%) | PC5 (0.02%) |
|---|---|---|---|---|---|
| Version | 0 | 0 | 0 | 0 | 0 |
| Sex | 0.11 | 0.85 | 0.01 | 0 | 0.01 |
| Donor | 0.89 | 1 | 0.99 | 0.99 | 0.99 |
| Age | 0.02 | 0.03 | 0.03 | 0.06 | 0.01 |
| Bas | 0 | 0 | 0.01 | 0.28 | 0 |
| Bmem | 0.12 | 0 | 0.29 | 0 | 0.01 |
| Bnv | 0.02 | 0.02 | 0.17 | 0.11 | 0.04 |
| CD4mem | 0.23 | 0.03 | 0.3 | 0.03 | 0.11 |
| CD4nv | 0 | 0.04 | 0.06 | 0.02 | 0.01 |
| CD8mem | 0.11 | 0 | 0.19 | 0.07 | 0 |
| CD8nv | 0 | 0 | 0.06 | 0.01 | 0.12 |
| Eos | 0.12 | 0.02 | 0.08 | 0.18 | 0.09 |
| Mono | 0.05 | 0.01 | 0 | 0.03 | 0 |
| Neu | 0.2 | 0.02 | 0.64 | 0 | 0.01 |
| NK | 0.1 | 0 | 0.18 | 0.02 | 0.02 |
| Treg | 0 | 0.02 | 0.02 | 0.13 | 0.09 |

Adjusted p

<=0.05

>0.05

### B: iv.BeCOME

#### Funnorm

| | PC1 (98.79%) | PC2 (0.27%) | PC3 (0.23%) | PC4 (0.15%) | PC5 (0.1%) |
|---|---|---|---|---|---|
| Version | 0.53 | 0.69 | 0.28 | 0.01 | 0 |
| Sex | 0 | 0.28 | 0.7 | 0 | 0 |
| Donor | 0.19 | 0.3 | 0.71 | 0.99 | 0.99 |
| Age | 0 | 0.05 | 0.07 | 0.48 | 0.3 |
| Bas | 0.01 | 0.18 | 0.15 | 0.01 | 0.04 |
| Bmem | 0.28 | 0.01 | 0.23 | 0 | 0.11 |
| Bnv | 0 | 0.02 | 0.07 | 0.26 | 0.27 |
| CD4mem | 0.03 | 0.01 | 0.04 | 0.41 | 0.46 |
| CD4nv | 0 | 0.02 | 0.13 | 0.79 | 0.02 |
| CD8mem | 0.02 | 0.5 | 0.45 | 0 | 0 |
| CD8nv | 0 | 0.06 | 0.14 | 0.55 | 0.11 |
| Eos | 0.17 | 0.12 | 0.21 | 0.02 | 0.08 |
| Mono | 0.02 | 0.04 | 0.24 | 0.2 | 0.02 |
| Neu | 0.02 | 0.05 | 0.04 | 0.81 | 0.01 |
| NK | 0.06 | 0.02 | 0.39 | 0.1 | 0.02 |
| Treg | 0.31 | 0.15 | 0.06 | 0.38 | 0.18 |

#### Funnorm + batch-correction

| | PC1 (99.92%) | PC2 (0.01%) | PC3 (0.01%) | PC4 (0.01%) | PC5 (0.01%) |
|---|---|---|---|---|---|
| Version | 0.02 | 0 | 0.01 | 0.01 | 0 |
| Sex | 0 | 0.07 | 0 | 0 | 0 |
| Donor | 0.03 | 0.07 | 0 | 0.01 | 0 |
| Age | 0 | 0.01 | 0 | 0 | 0 |
| Bas | 0 | 0 | 0.01 | 0 | 0.02 |
| Bmem | 0.11 | 0.01 | 0.08 | 0 | 0 |
| Bnv | 0 | 0.01 | 0 | 0.01 | 0 |
| CD4mem | 0.01 | 0 | 0.01 | 0 | 0 |
| CD4nv | 0.01 | 0.01 | 0.01 | 0 | 0 |
| CD8mem | 0.01 | 0.07 | 0 | 0 | 0 |
| CD8nv | 0.01 | 0.03 | 0.01 | 0 | 0 |
| Eos | 0.01 | 0 | 0.02 | 0.06 | 0.02 |
| Mono | 0 | 0.02 | 0.01 | 0 | 0 |
| Neu | 0.01 | 0.01 | 0.01 | 0 | 0 |
| NK | 0 | 0.04 | 0.01 | 0 | 0 |
| Treg | 0.06 | 0 | 0 | 0.01 | 0.05 |

Adjusted p
- <=0.05
- >0.05

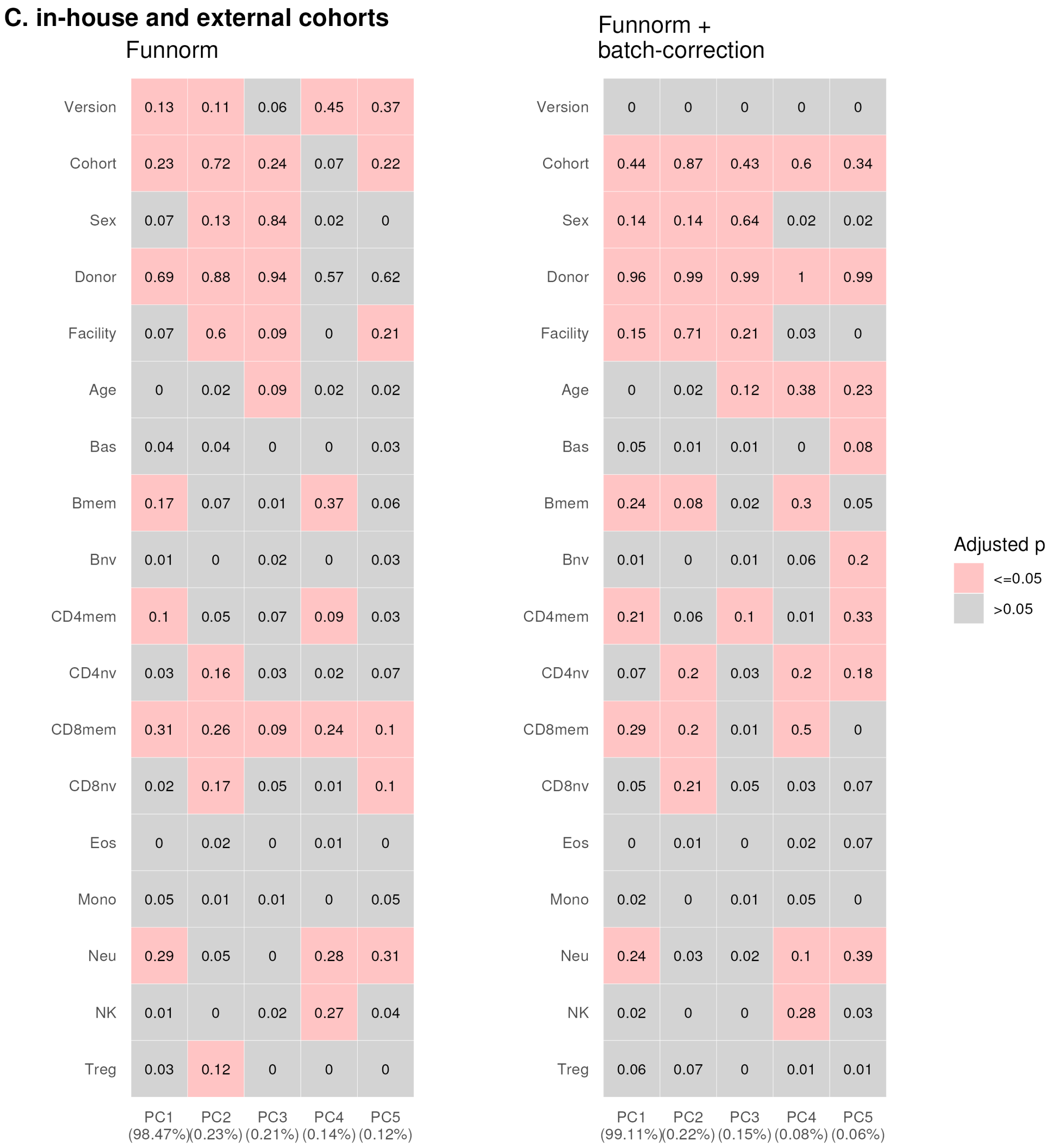


**Supplementary Figure 5.** Association of variables and loadings of the top five principal components using functional normalized, functional normalized and batch-corrected data **(A)** the three cohorts (VHAS, CLHNS and CALERIE) processed by in-house facility. **(B)** each cohort separately. **(C)** the three in-house cohorts and the external validation cohort (BeCOME). An overall F test was carried out for 5*18 = 90 separate simple linear regression models for each combination of 5 PCs as dependent variable and 18 predictor variables: Facility, version, sex, cohort, donor, age, and 12 cell type proportions. Data processing method is indicated for each panel and $R^2$ is indicated in each cell. Accounted variance of PCs are shown in brackets in the x-axis label. *p*-values were Bonferroni adjusted.

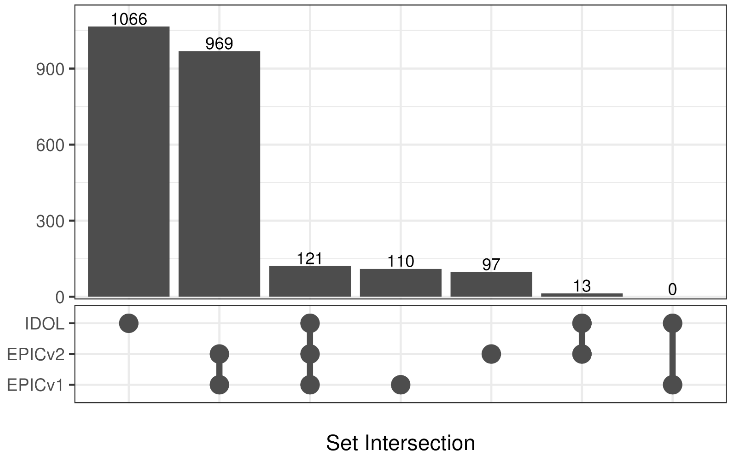

1066
969
121
110
97
13
0
900
600
300
0
IDOL
EPICv2
EPICv1
Set Intersection

**Supplementary Figure 6.** Upset plot denoting overlap of probes selected when using the IDOL pre-selected probes and auto-selected probes in EPICv1 and EPICv2.

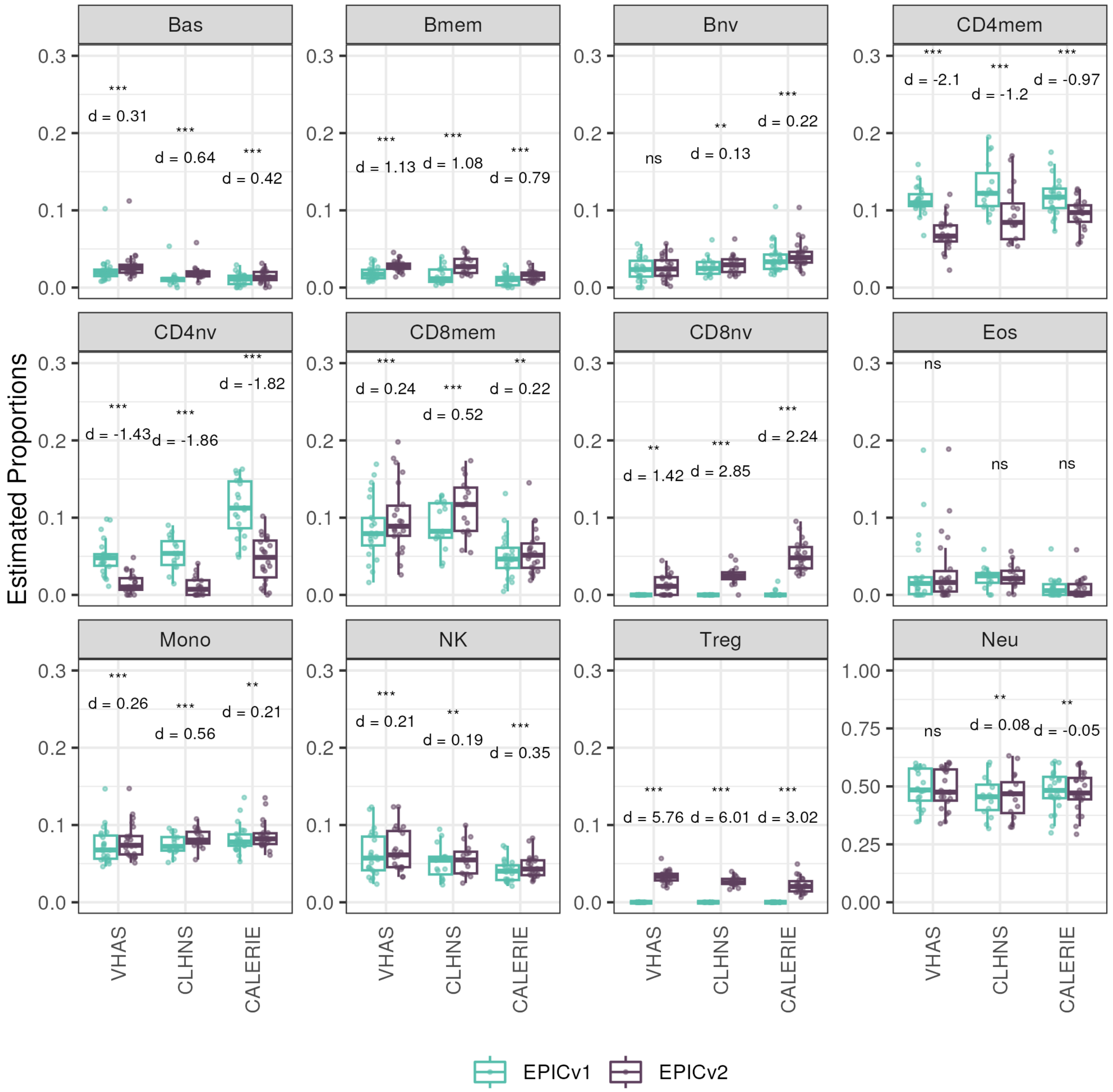


**Supplementary Figure 7.** Differences in DNA methylation-based immune cell type proportions estimated using the "auto" method on matched samples assessed on EPICv1 and EPICv2 in VHAS, CLHNS, and CALERIE. Paired t-tests were performed to compare estimates between EPICv1 and EPICv2, and p-values were derived. Statistical significance was defined as Bonferroni adjusted p-value <0.05. ** denotes Bonferroni p <0.05, *** denotes Bonferroni p<0.001, "ns" denotes "not significant", and "d" denotes effect size measured using Cohen's d. A positive Cohen's d indicates higher average estimated cell proportions in EPICv2 compared to EPICv1.

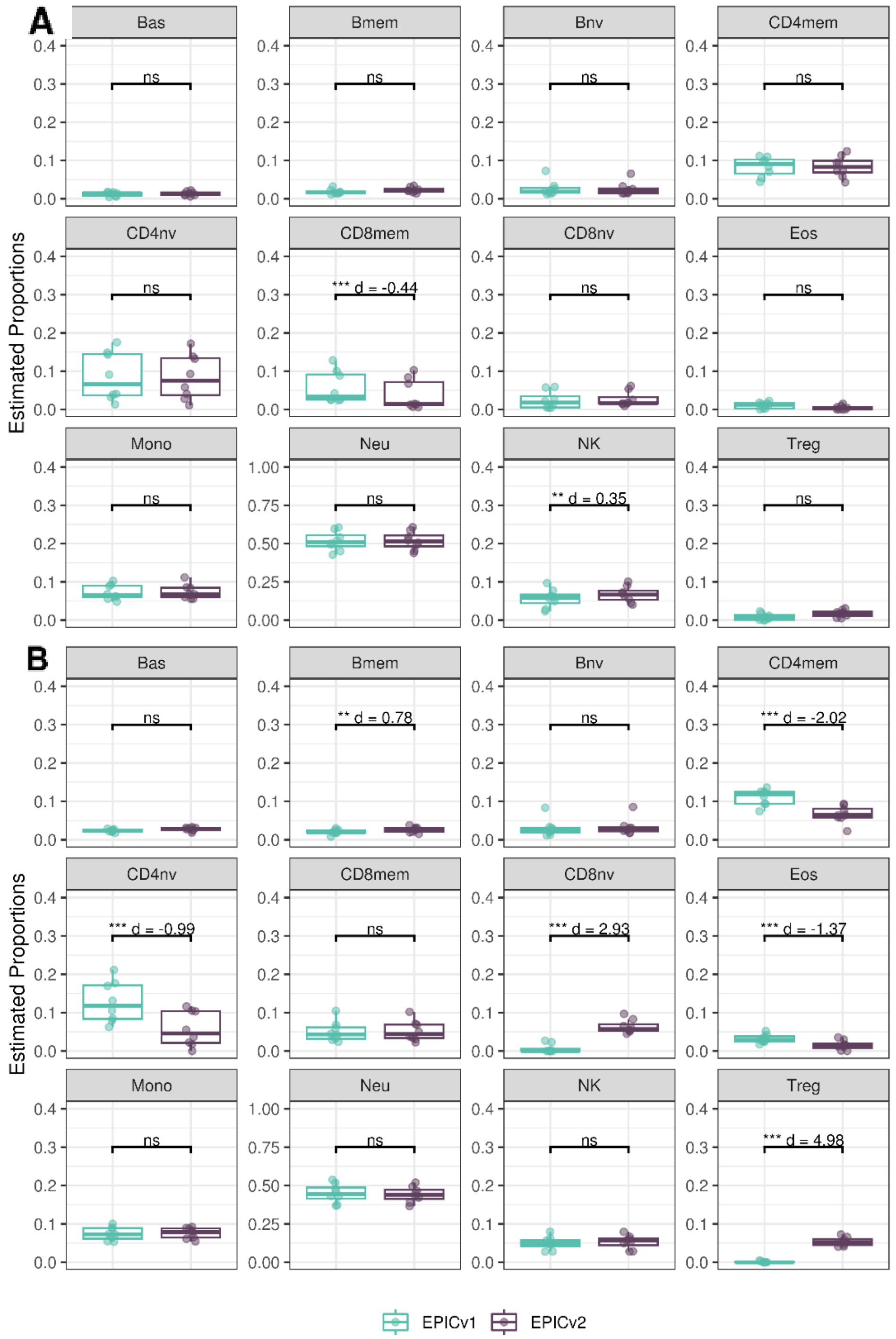


**Supplementary Figure 8.** Differences in DNA methylation-based immune cell type proportions estimated on matched samples assessed on EPICv1 and EPICv2 in BeCOME. (A). using the “IDOL” method. **(B).** using the “auto” method.

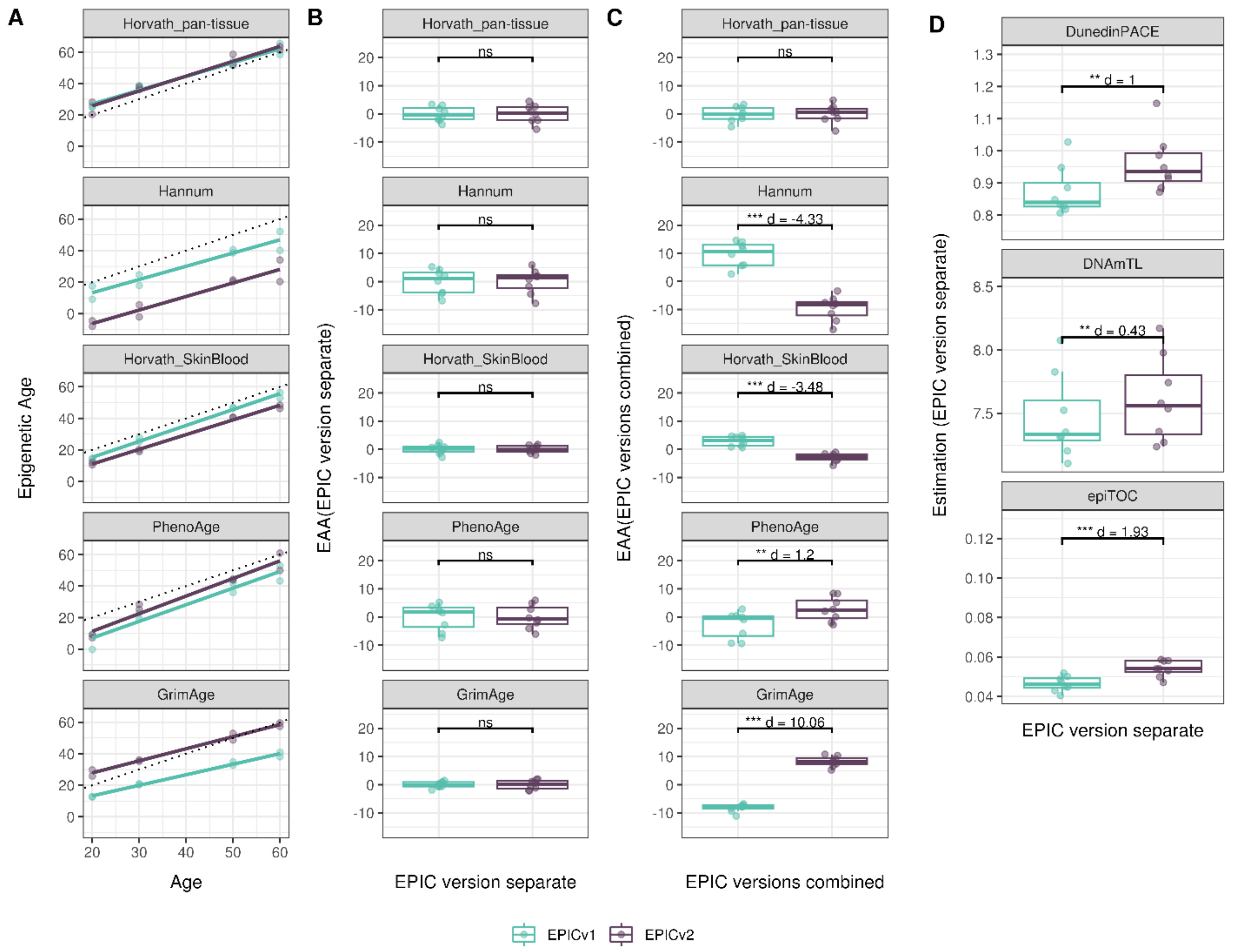


**Supplementary Figure 9.** Epigenetic age estimated on matched samples assessed on EPICv1 and EPICv2 in BeCOME using functional normalization. **(A)** Scatter plot of Horvath pan-tissue, Hannum, Horvath skin and blood, PhenoAge, and GrimAge clock estimates (Y axis) and chronological age (X axis) with dotted line indicating x=y, coloured by EPIC version. **(B-C)** Boxplots comparing EPICv1 and EPICv2 EAAs calculated by considering EPIC versions separately and combined, respectively. **(D)** Boxplots comparing DunedinPACE, DNAmTL and epiTOC estimates calculated by considering EPIC versions separately between EPICv1 and EPICv2. Paired t-tests were performed to compare estimates between EPICv1 and EPICv2, and p-values were derived. Statistical significance was defined as Bonferroni adjusted p-value <0.05. ** denotes Bonferroni p <0.05, *** denotes Bonferroni p<0.001, “ns” denotes “not significant”, and “d” denotes effect size measured using Cohen’s d. A positive Cohen’s d indicates estimates in EPICv2 compared to EPICv1.

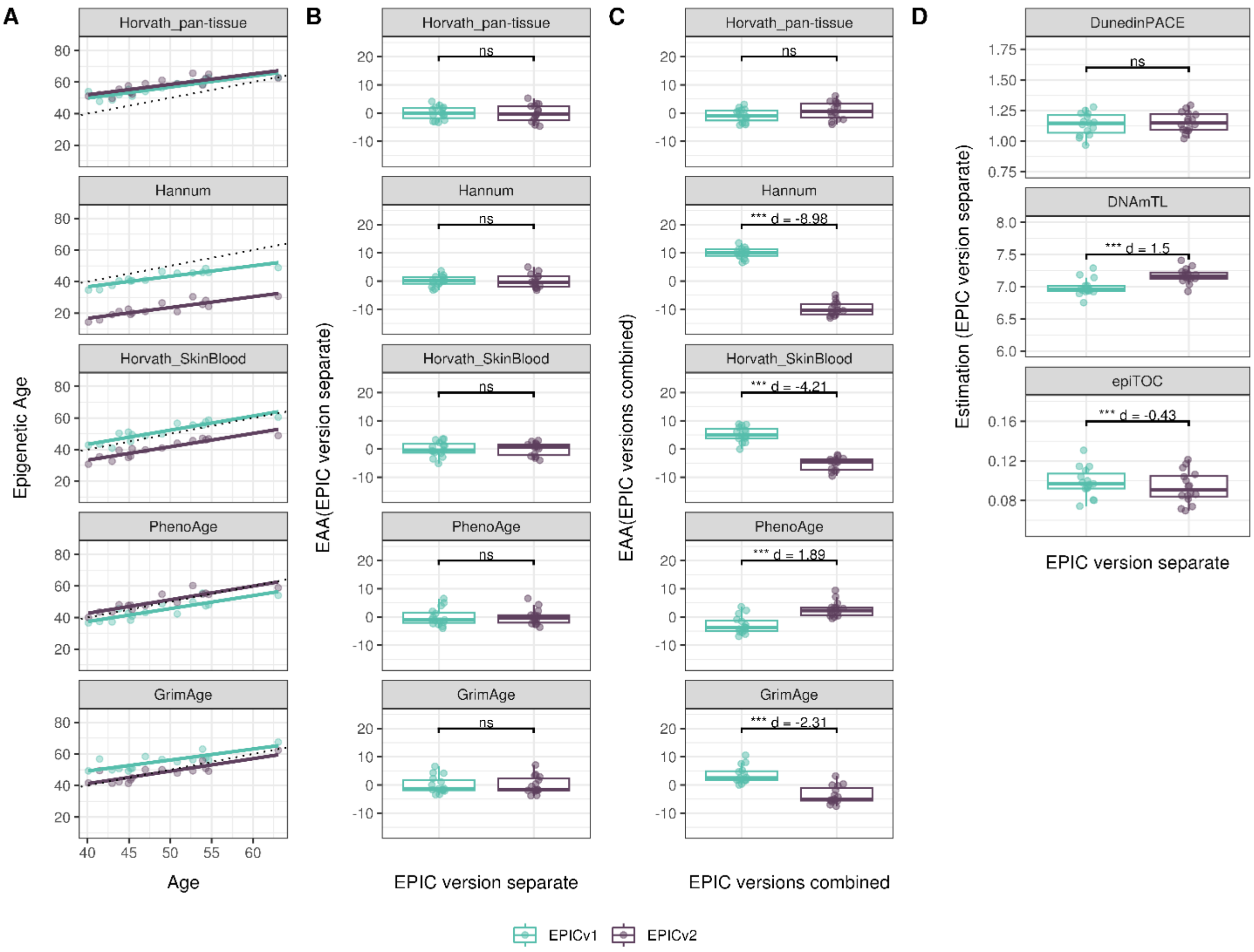


**Supplementary Figure 10**. Epigenetic age estimated on matched samples assessed on EPICv1 and EPICv2 in CLHNS using functional normalization. **(A)** Scatter plot of Horvath pan-tissue, Hannum, Horvath skin and blood, PhenoAge, and GrimAge clock estimates (Y axis) and chronological age (X axis) with dotted line indicating x=y, coloured by EPIC version. **(B-C)** Boxplots comparing EPICv1 and EPICv2 EAAs calculated by considering EPIC versions separately and combined, respectively. **(D)** Boxplots comparing DunedinPACE, DNAmTL and epiTOC estimates calculated by considering EPIC versions separately between EPICv1 and EPICv2. Paired t-tests were performed to compare estimates between EPICv1 and EPICv2, and p-values were derived. Statistical significance was defined as Bonferroni adjusted p-value <0.05. ** denotes Bonferroni p <0.05, *** denotes Bonferroni p<0.001, "ns" denotes "not significant", and "d" denotes effect size measured using Cohen's d. A positive Cohen's d indicates estimates in EPICv2 compared to EPICv1.

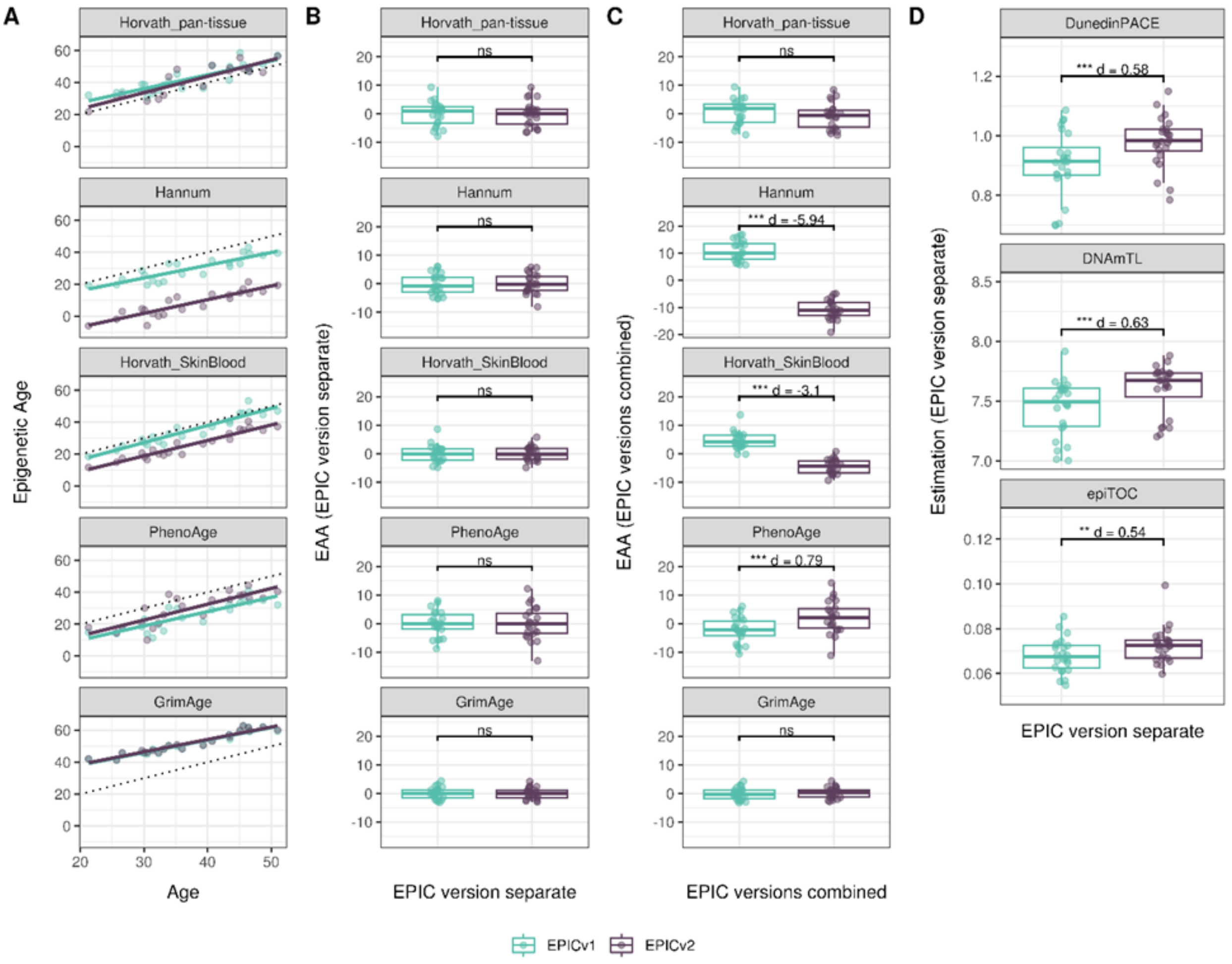


**Supplementary Figure 11**. Epigenetic age estimated on matched samples assessed on EPICv1 and EPICv2 in CALERIE using functional normalization. **(A)** Scatter plot of Horvath pan-tissue, Hannum, Horvath skin and blood, PhenoAge, and GrimAge clock estimates (Y axis) and chronological age (X axis) with dotted line indicating x=y, coloured by EPIC version. **(B-C)** Boxplots comparing EPICv1 and EPICv2 EAAs calculated by considering EPIC versions separately and combined, respectively. **(D)** Boxplots comparing DunedinPACE, DNAmTL and epiTOC estimates calculated by considering EPIC versions separately between EPICv1 and EPICv2. Paired t-tests were performed to compare estimates between EPICv1 and EPICv2, and p-values were derived. Statistical significance was defined as Bonferroni adjusted p-value <0.05. ** denotes Bonferroni p <0.05, *** denotes Bonferroni p<0.001, "ns" denotes "not significant", and "d" denotes effect size measured using Cohen's d. A positive Cohen's d indicates estimates in EPICv2 compared to EPICv1.

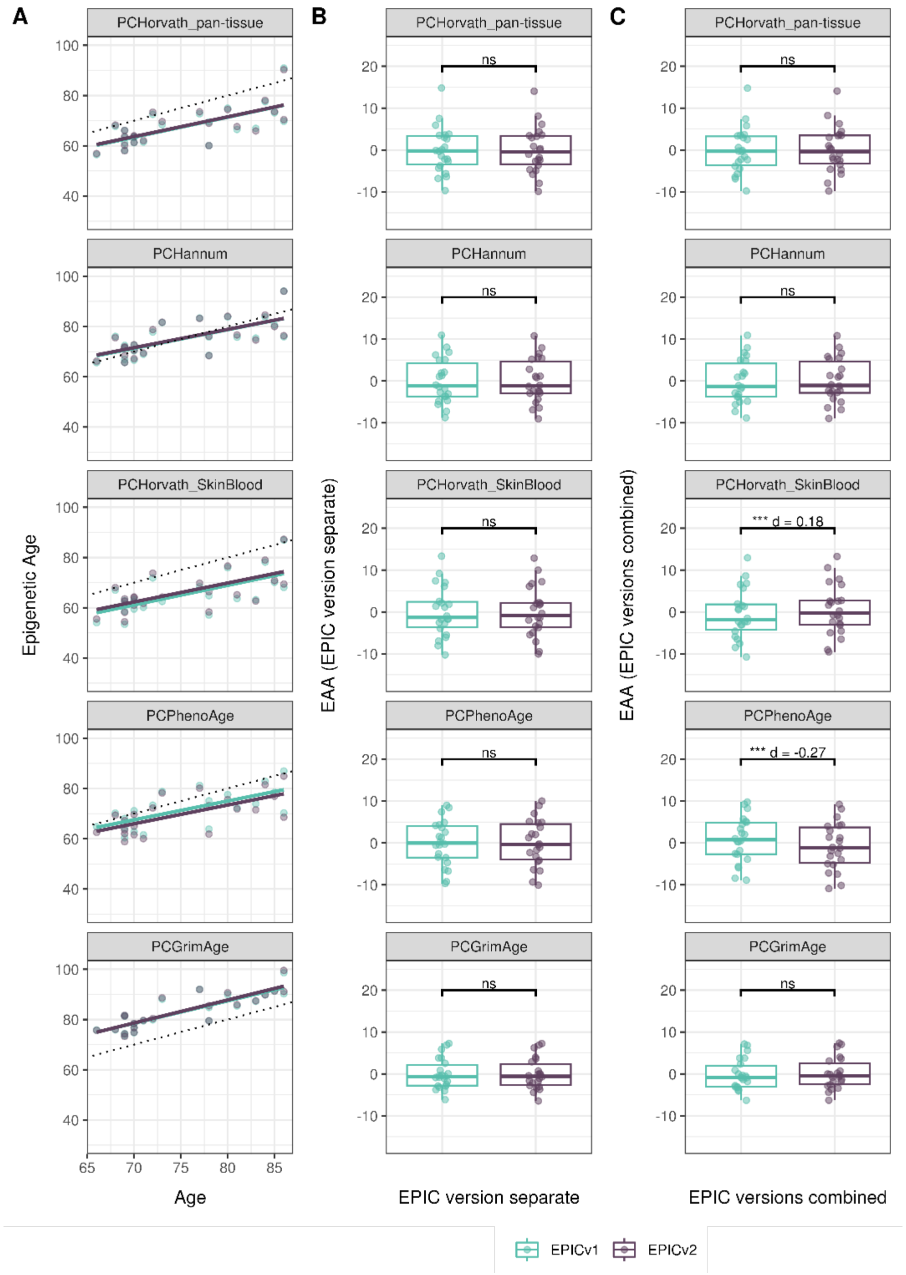


**Supplementary Figure 12.** Epigenetic age estimated on matched samples assessed on EPICv1 and EPICv2 in VHAS using functional normalization. **(A)** Scatter plot of PC clocks of Horvath pan-tissue, Hannum, Horvath skin and blood, PhenoAge, and GrimAge clock ages (Y axis) and chronological age (X axis) with dotted line indicating x=y, coloured by EPIC version. **(B-C)** Boxplots comparing EPICv1 and EPICv2 EAAs calculated by considering EPIC versions separately, combined, respectively. Paired t-tests were performed to compare estimates between EPICv1 and EPICv2, and p-values were derived. Statistical significance was defined as Bonferroni adjusted p-value <0.05. ** denotes Bonferroni p <0.05, *** denotes Bonferroni p<0.001, "ns" denotes "not significant", and "d" denotes effect size measured using Cohen's d. A positive Cohen's d indicates higher estimates in EPICv2 compared to EPICv1.

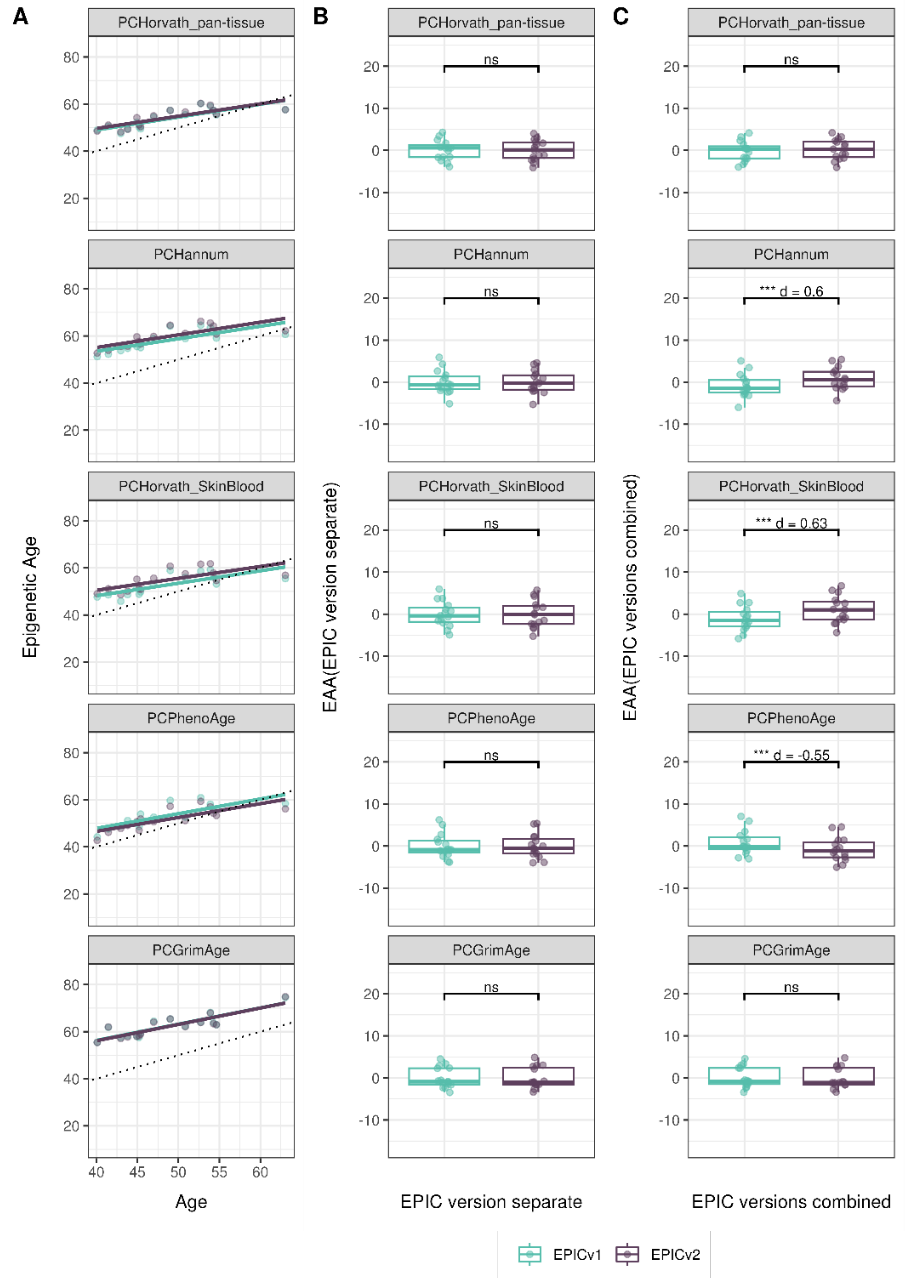


**Supplementary Figure 13.** Epigenetic age estimated on matched samples assessed on EPICv1 and EPICv2 in CLHNS using functional normalization. **(A)** Scatter plot of PC clocks of Horvath pan-tissue, Hannum, Horvath skin and blood, PhenoAge, and GrimAge clock ages (Y axis) and chronological age (X axis) with dotted line indicating x=y, coloured by EPIC version. **(B-C)** Boxplots comparing EPICv1 and EPICv2 EAAs calculated by considering EPIC versions separately, combined, respectively. Paired t-tests were performed to compare estimates between EPICv1 and EPICv2, and p-values were derived. Statistical significance was defined as Bonferroni adjusted p-value <0.05. ** denotes Bonferroni p <0.05, *** denotes Bonferroni p<0.001, "ns" denotes "not significant", and "d" denotes effect size measured using Cohen's d. A positive Cohen's d indicates higher estimates in EPICv2 compared to EPICv1.

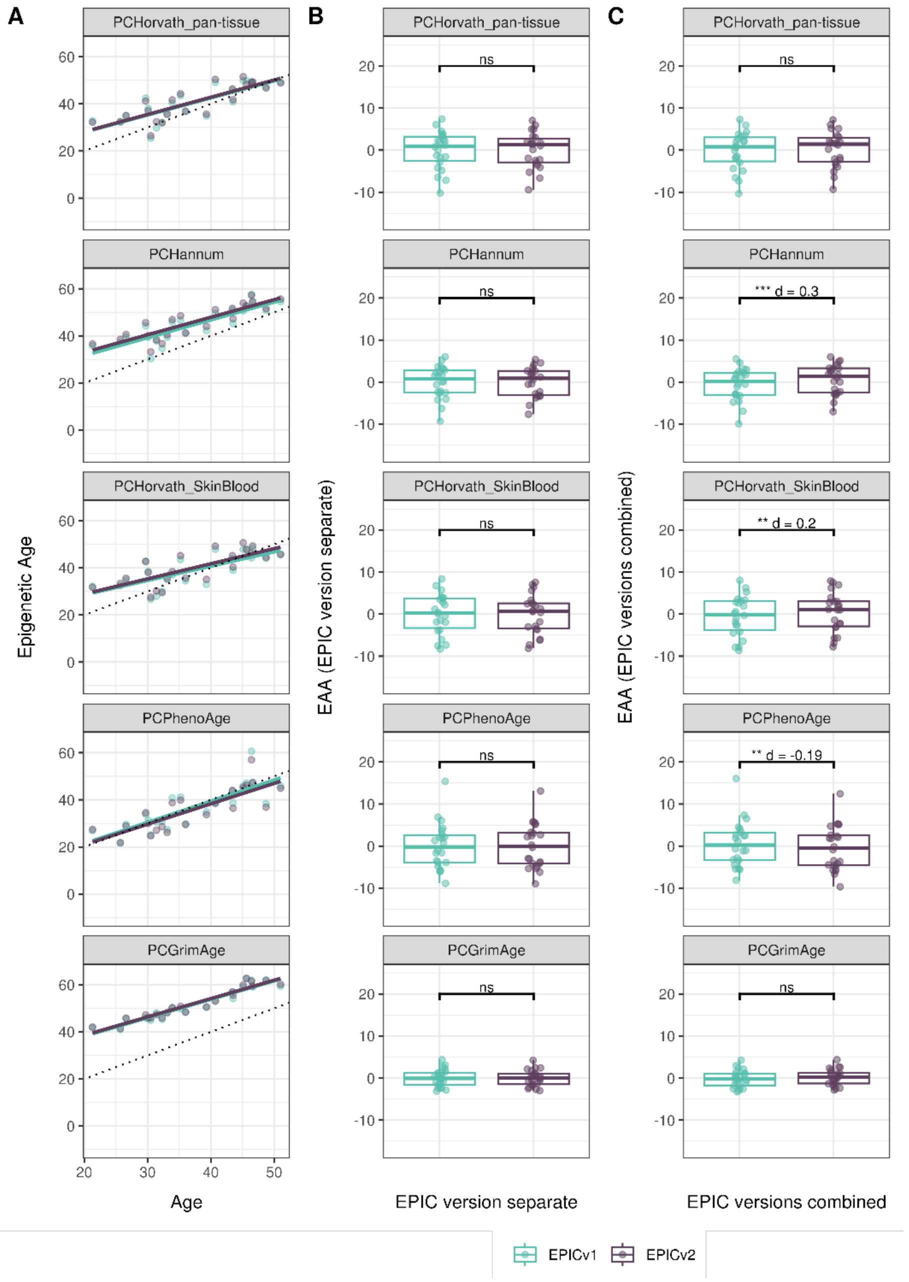


**Supplementary Figure 14.** Epigenetic age estimated on matched samples assessed on EPICv1 and EPICv2 in CALERIE using functional normalization. **(A)** Scatter plot of PC clocks of Horvath pan-tissue, Hannum, Horvath skin and blood, PhenoAge, and GrimAge clock ages (Y axis) and chronological age (X axis) with dotted line indicating x=y, coloured by EPIC version. **(B-C)** Boxplots comparing EPICv1 and EPICv2 EAAs calculated by considering EPIC versions separately, combined, respectively. Paired t-tests were performed to compare estimates between EPICv1 and EPICv2, and p-values were derived. Statistical significance was defined as Bonferroni adjusted p-value <0.05. ** denotes Bonferroni p <0.05, *** denotes Bonferroni p<0.001, "ns" denotes "not significant", and "d" denotes effect size measured using Cohen's d. A positive Cohen's d indicates higher estimates in EPICv2 compared to EPICv1.

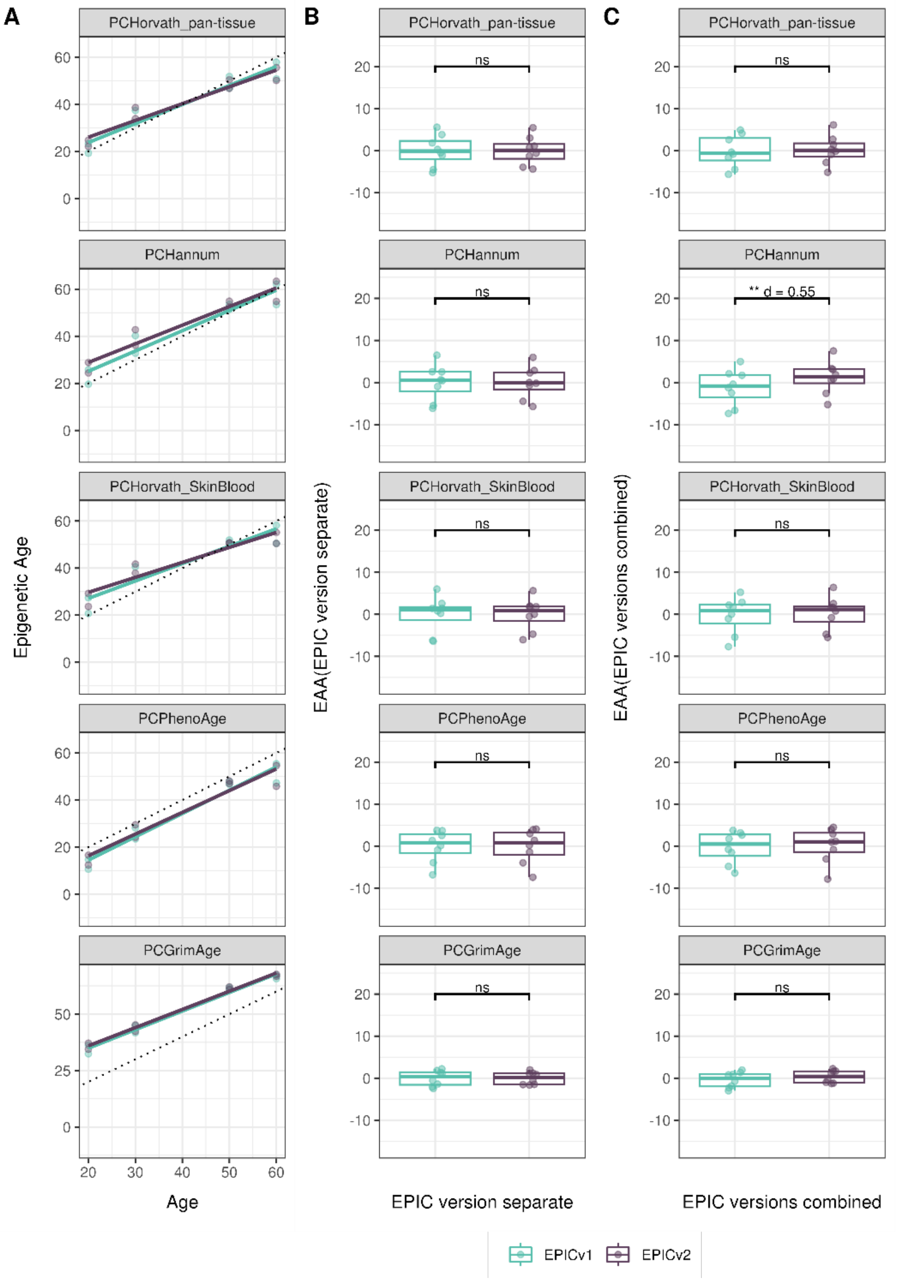


**Supplementary Figure 15.** Epigenetic age estimated on matched samples assessed on EPICv1 and EPICv2 in BeCOME using functional normalization. **(A)** Scatter plot of PC clocks of Horvath pan-tissue, Hannum, Horvath skin and blood, PhenoAge, and GrimAge clock ages (Y axis) and chronological age (X axis) with dotted line indicating x=y, coloured by EPIC version. **(B-C)** Boxplots comparing EPICv1 and EPICv2 EAAs calculated by considering EPIC versions separately, combined, respectively. Paired t-tests were performed to compare estimates between EPICv1 and EPICv2, and p-values were derived. Statistical significance was defined as Bonferroni adjusted p-value <0.05. ** denotes Bonferroni p <0.05, *** denotes Bonferroni p<0.001, "ns" denotes "not significant", and "d" denotes effect size measured using Cohen's d. A positive Cohen's d indicates higher estimates in EPICv2 compared to EPICv1.

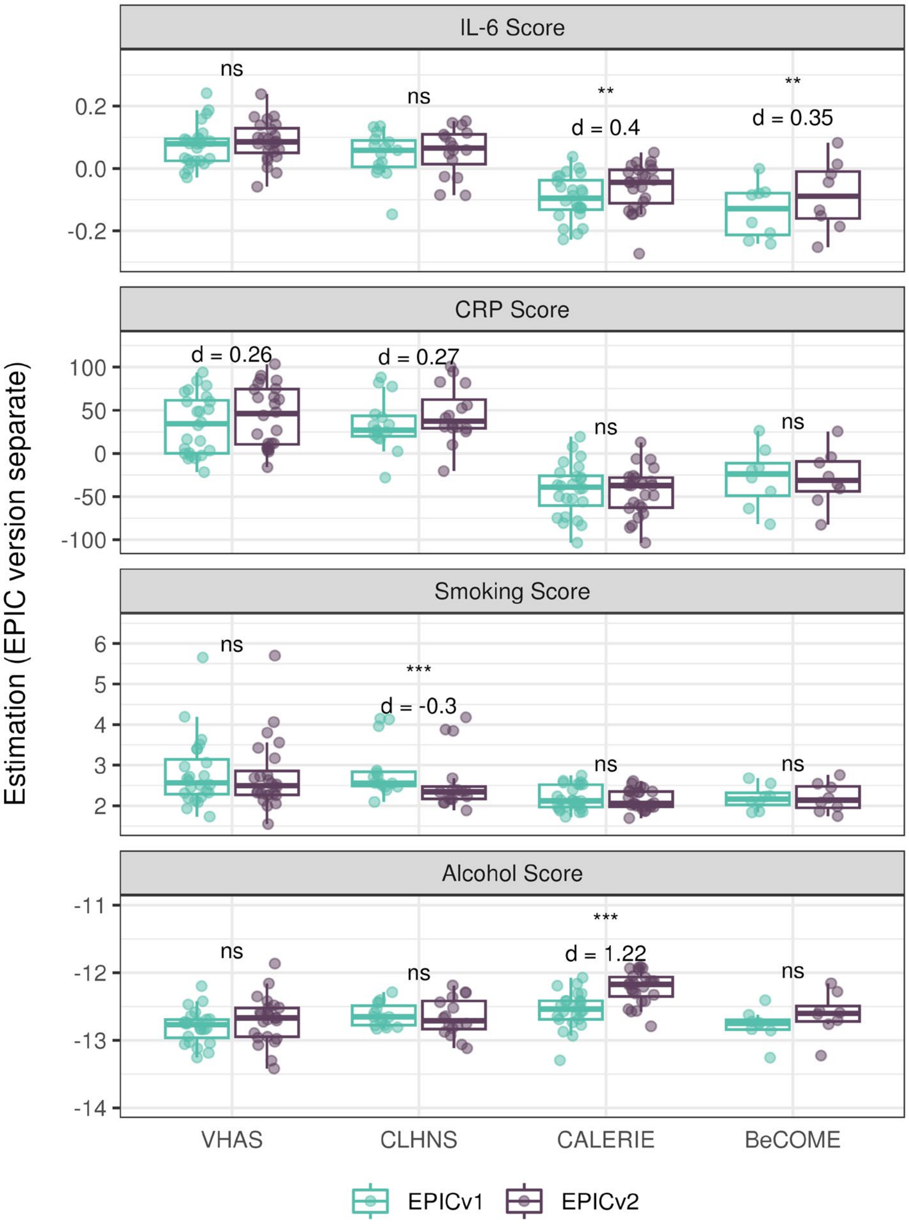


**Supplementary Figure 16**. DNA methylation-based inflammation and lifestyle biomarkers estimated on matched samples assessed on EPICv1 and EPICv2 in VHAS**,** CLHNS**,** CALERIE and BeCOME using functional normalization. Boxplots comparing EPICv1 and EPICv2 proxy IL-6, CRP, smoking, and alcohol scores calculated by considering EPIC versions separately. Paired t-tests were performed to compare estimates between EPICv1 and EPICv2, and p-values were derived. Statistical significance was defined as Bonferroni adjusted p-value <0.05. ** denotes Bonferroni p <0.05, *** denotes Bonferroni p<0.001, "ns" denotes "not significant", and "d" denotes effect size measured using Cohen's d. A positive Cohen's d indicates higher average estimated cell proportions in EPICv2 compared to EPICv1.

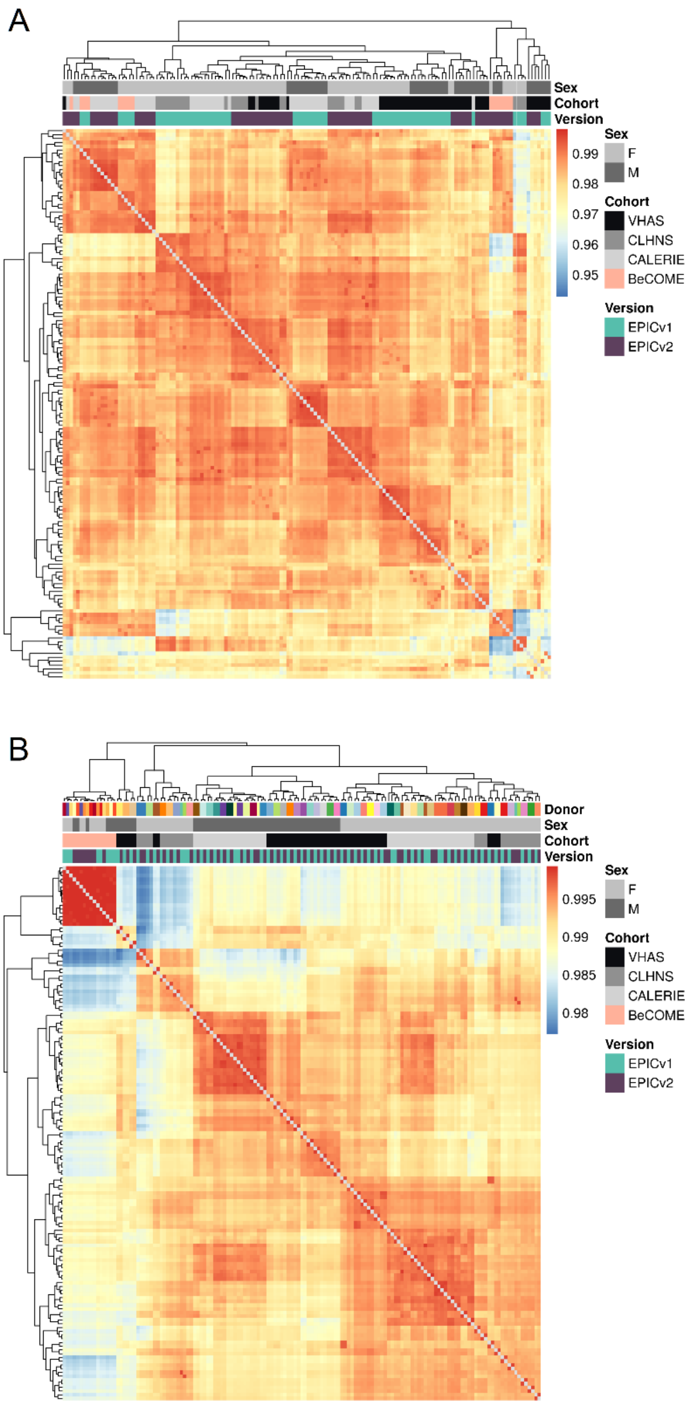


**Supplementary Figure 17.** Unsupervised hierarchical clustering using complete linkage with Euclidean distance on sample-to-sample Pearson correlations, calculated using the 721,378 probes shared between EPICv1 and EPICv2 using **A).** raw data; **B).**functional normalization with batch-correction for EPIC version, chip and row; blue to red color range denotes Pearson correlation from low to high.

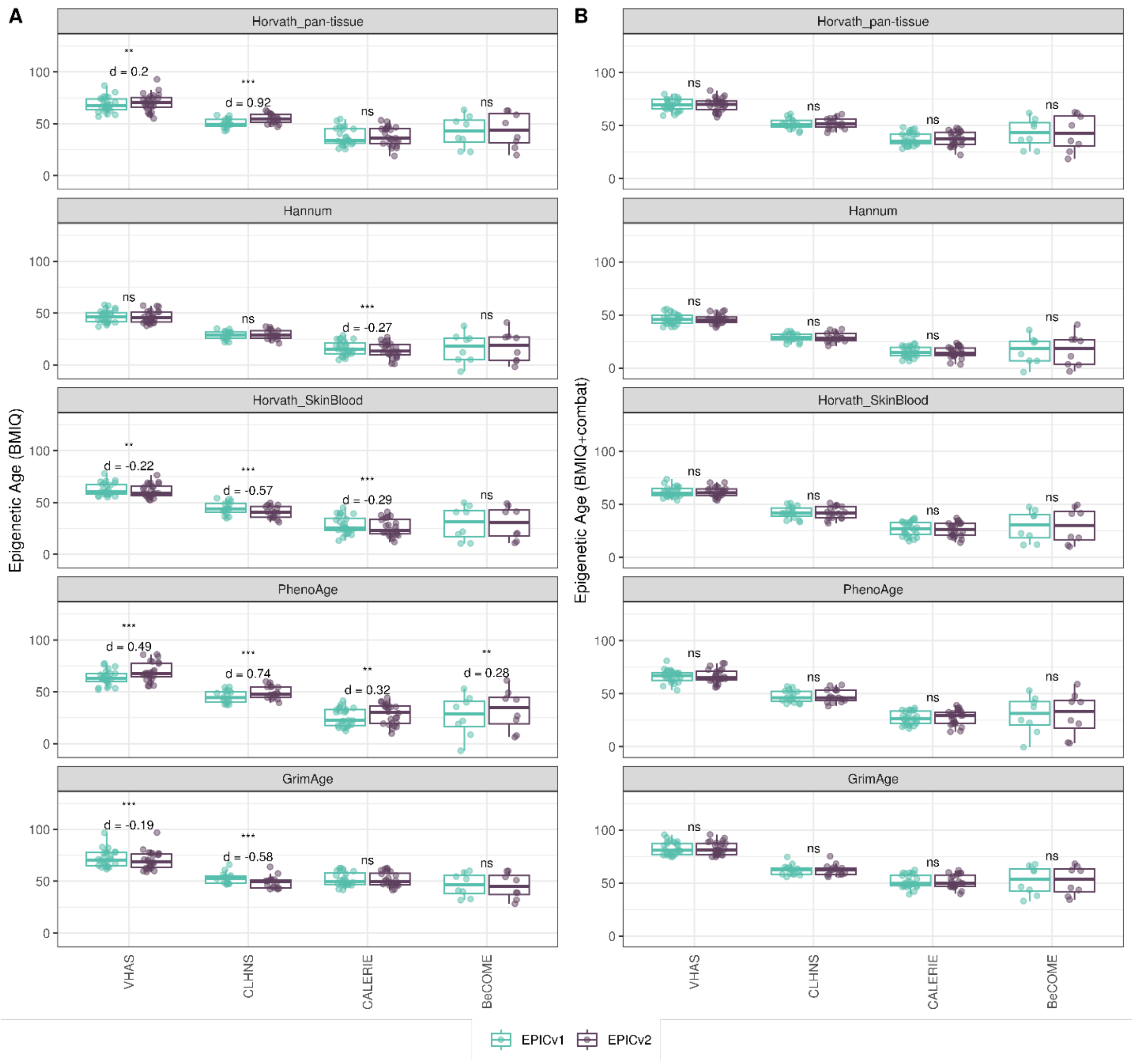


**Supplementary Figure 18.** Differences in epigenetic ages between EPICv1 and EPICv2 using the Horvath pan-tissue, Hannum, Horvath skin and blood, PhenoAge, and GrimAge clocks in VHAS, CLHNS, and CALERIE when using **(A)** normalizing EPICv1 and EPICv2 together using BMIQ normalization; **(B)** normalizing EPICv1 and EPICv2 together using BMIQ normalization with batch-correction for EPIC version, chip and row. Paired t-tests were performed to compare estimates between EPICv1 and EPICv2, and p-values were derived. Statistical significance was defined as Bonferroni adjusted p-value <0.05. ** denotes Bonferroni p <0.05, *** denotes Bonferroni p<0.001, "ns" denotes "not significant", and "d" denotes effect size measured using Cohen's d. A positive Cohen's d indicates estimates in EPICv2 compared to EPICv1.

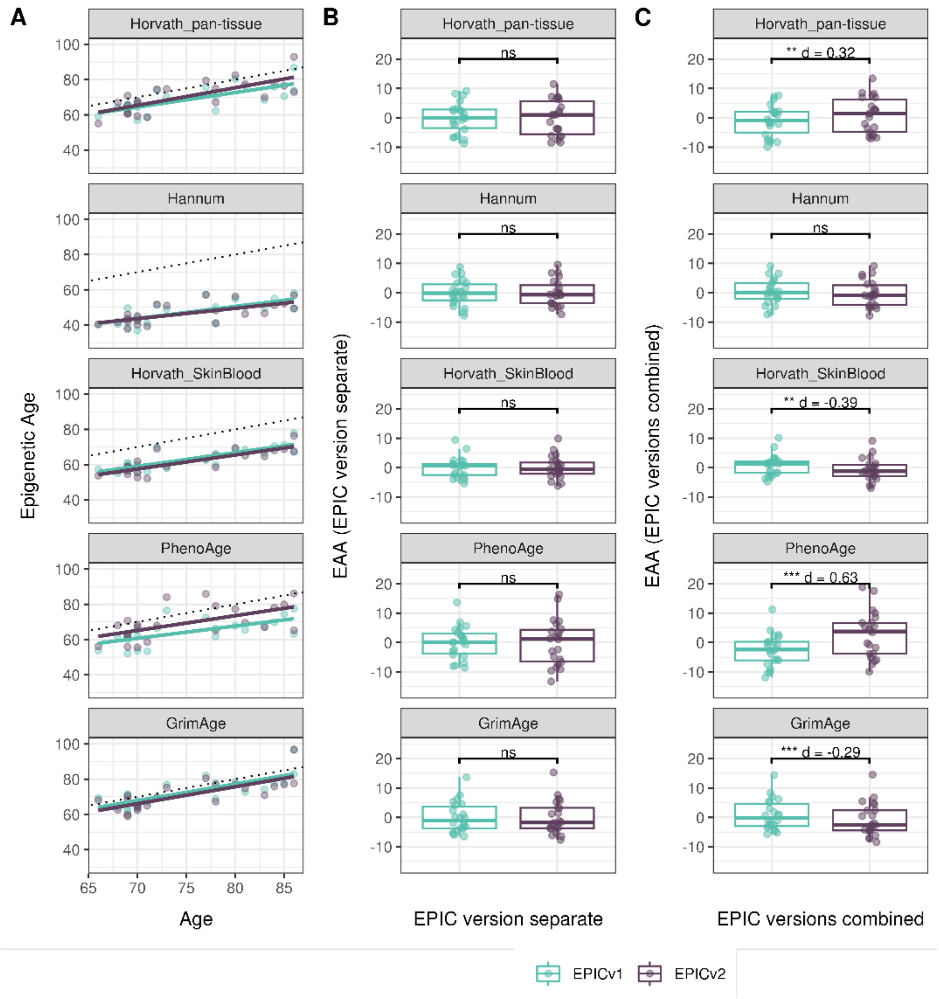


**Supplementary Figure 19.** Epigenetic age estimated on matched samples after normalizing EPICv1 and EPICv2 together using BMIQ in VHAS. **(A)** Scatter plot of Horvath pan-tissue, Hannum, Horvath skin and blood, PhenoAge, and GrimAge clock estimates (Y axis) and chronological age (X axis) with dotted line indicating x=y, coloured by EPIC version. **(B-C)** Boxplots comparing EPICv1 and EPICv2 EAAs calculated by considering EPIC versions separately and combined, respectively. Paired t-tests were performed to compare estimates between EPICv1 and EPICv2, and p-values were derived. Statistical significance was defined as Bonferroni adjusted p-value <0.05. ** denotes Bonferroni p <0.05, *** denotes Bonferroni p<0.001, "ns" denotes "not significant", and "d" denotes effect size measured using Cohen's d. A positive Cohen's d indicates estimates in EPICv2 compared to EPICv1.

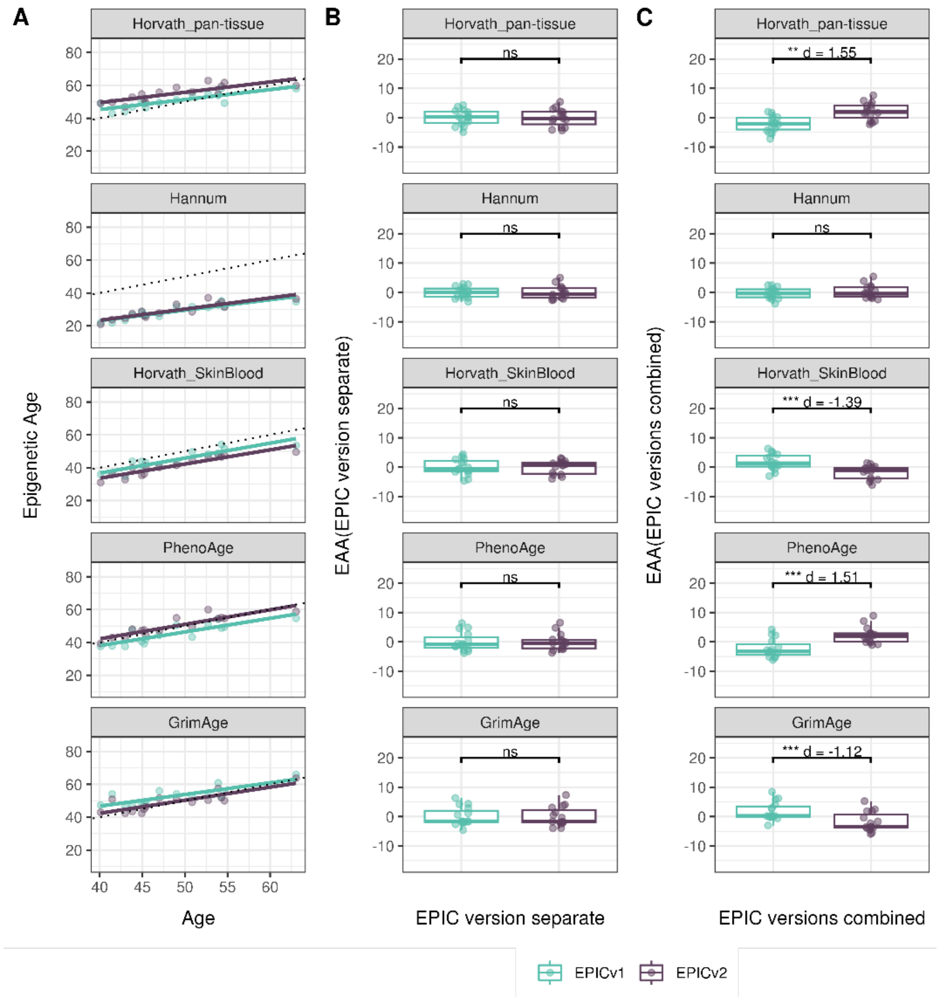


**Supplementary Figure 20.** Epigenetic age estimated on matched samples after normalizing EPICv1 and EPICv2 together using BMIQ in CLHNS. **(A)** Scatter plot of Horvath pan-tissue, Hannum, Horvath skin and blood, PhenoAge, and GrimAge clock estimates (Y axis) and chronological age (X axis) with dotted line indicating x=y, coloured by EPIC version. **(B-C)** Boxplots comparing EPICv1 and EPICv2 EAAs calculated by considering EPIC versions separately and combined, respectively. Paired t-tests were performed to compare estimates between EPICv1 and EPICv2, and p-values were derived. Statistical significance was defined as Bonferroni adjusted p-value <0.05. ** denotes Bonferroni p <0.05, *** denotes Bonferroni p<0.001, "ns" denotes "not significant", and "d" denotes effect size measured using Cohen's d. A positive Cohen's d indicates estimates in EPICv2 compared to EPICv1.

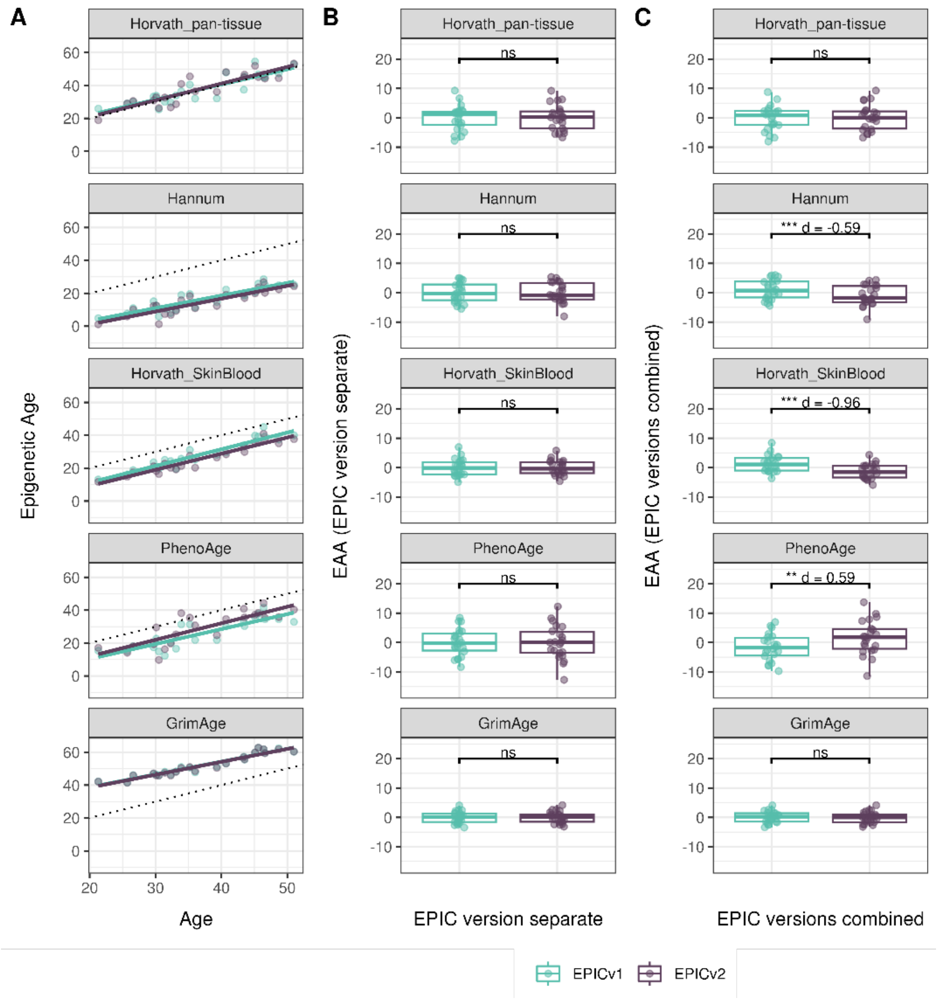


**Supplementary Figure 21.** Epigenetic age estimated on matched samples after normalizing EPICv1 and EPICv2 together using BMIQ in CALERIE. **(A)** Scatter plot of Horvath pan-tissue, Hannum, Horvath skin and blood, PhenoAge, and GrimAge clock estimates (Y axis) and chronological age (X axis) with dotted line indicating x=y, coloured by EPIC version. **(B-C)** Boxplots comparing EPICv1 and EPICv2 EAAs calculated by considering EPIC versions separately and combined, respectively. Paired t-tests were performed to compare estimates between EPICv1 and EPICv2, and p-values were derived. Statistical significance was defined as Bonferroni adjusted p-value <0.05. ** denotes Bonferroni p <0.05, *** denotes Bonferroni p<0.001, "ns" denotes "not significant", and "d" denotes effect size measured using Cohen's d. A positive Cohen's d indicates estimates in EPICv2 compared to EPICv1.

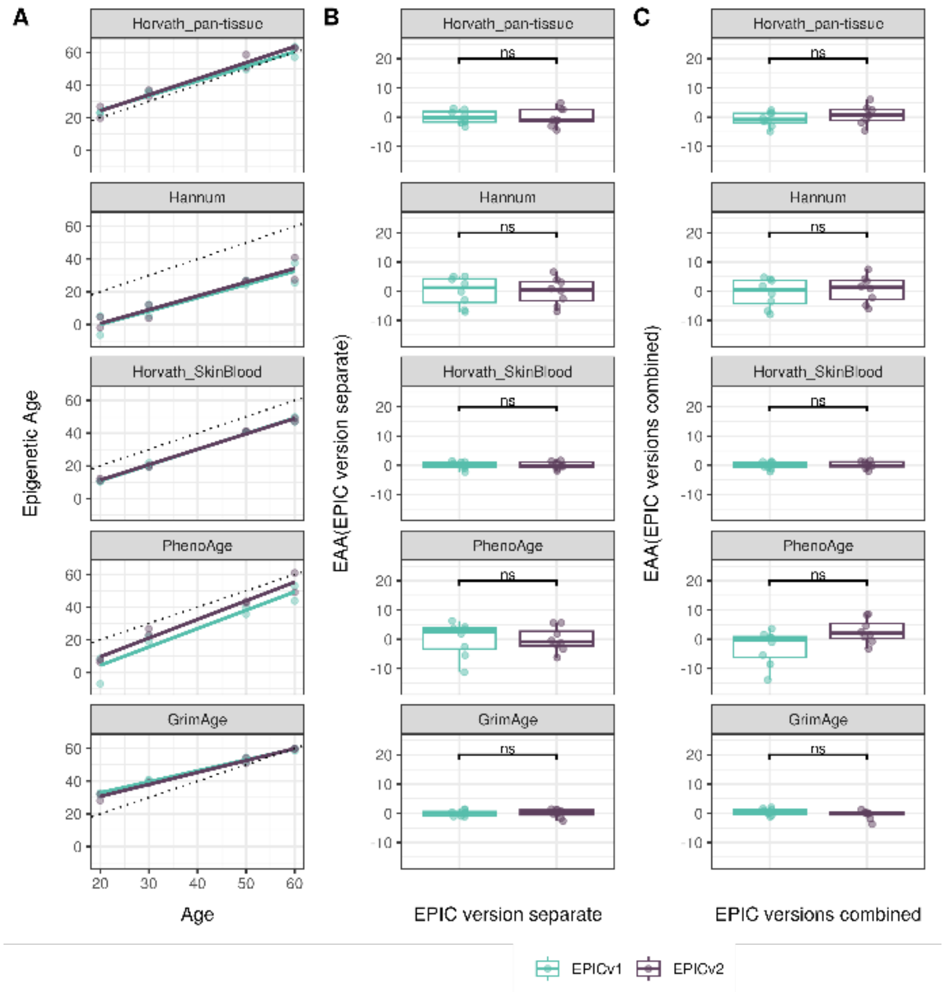


**Supplementary Figure 22.** Epigenetic age estimated on matched samples after normalizing EPICv1 and EPICv2 together using BMIQ in BeCOME. **(A)** Scatter plot of Horvath pan-tissue, Hannum, Horvath skin and blood, PhenoAge, and GrimAge clock estimates (Y axis) and chronological age (X axis) with dotted line indicating x=y, coloured by EPIC version. **(B-C)** Boxplots comparing EPICv1 and EPICv2 EAAs calculated by considering EPIC versions separately and combined, respectively. Paired t-tests were performed to compare estimates between EPICv1 and EPICv2, and p-values were derived. Statistical significance was defined as Bonferroni adjusted p-value <0.05. ** denotes Bonferroni p <0.05, *** denotes Bonferroni p<0.001, “ns” denotes “not significant”, and “d” denotes effect size measured using Cohen’s d. A positive Cohen’s d indicates estimates in EPICv2 compared to EPICv1.

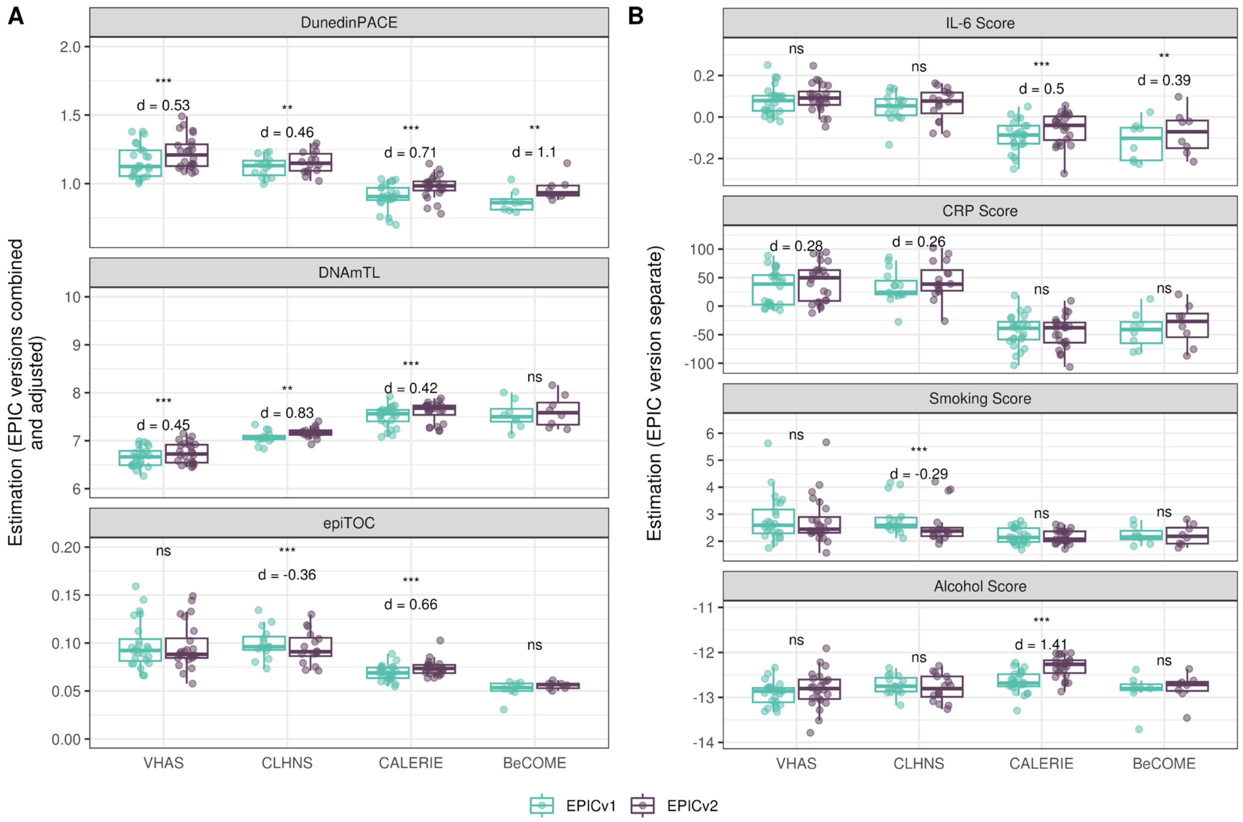


**Supplementary Figure 23.** Rate-based and other clock estimates **(A)**, DNA methylation-based inflammation and lifestyle biomarkers **(B)** obtained after normalizing EPICv1 and EPICv2 together using BMIQ in VHAS, CLHNS, CALERIE, and BeCOME. Paired t-tests were performed to compare estimates between EPICv1 and EPICv2, and p-values were derived. Statistical significance was defined as Bonferroni adjusted *p*-value <0.05. ** denotes Bonferroni *p* <0.05, *** denotes Bonferroni *p* <0.001, "ns" denotes "not significant", and "d" denotes effect size measured using Cohen's d. A positive Cohen's d indicates estimates in EPICv2 compared to EPICv1.

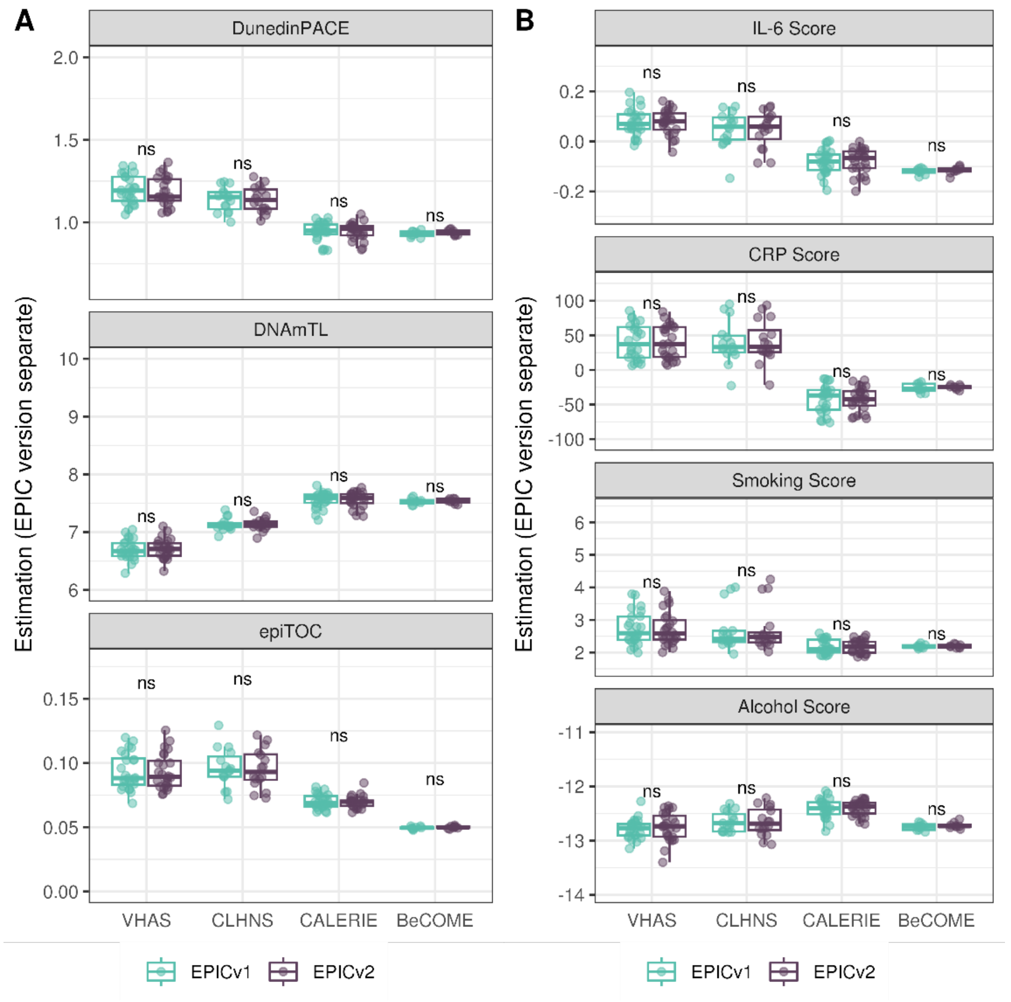


**Supplementary Figure 24.** DunedinPACE, DNAmTL, epiTOC, and DNA methylation-based inflammation and lifestyle biomarkers calculated by considering EPIC versions separately after functional normalization with batch-correction for EPIC version, chip and row in VHAS, CLHNS, CALERIE, and BeCOME. **(A)** rate-based and other clocks **(B)**. DNA methylation-based inflammation and lifestyle biomarkers. Paired t-tests were performed to compare estimates between EPICv1 and EPICv2, and p-values were derived. Statistical significance was defined as Bonferroni adjusted *p*-value <0.05. ** denotes Bonferroni *p* <0.05, *** denotes Bonferroni *p* <0.001, “ns” denotes “not significant”, and “d” denotes effect size measured using Cohen’s d. A positive Cohen’s d indicates estimates in EPICv2 compared to EPICv1.

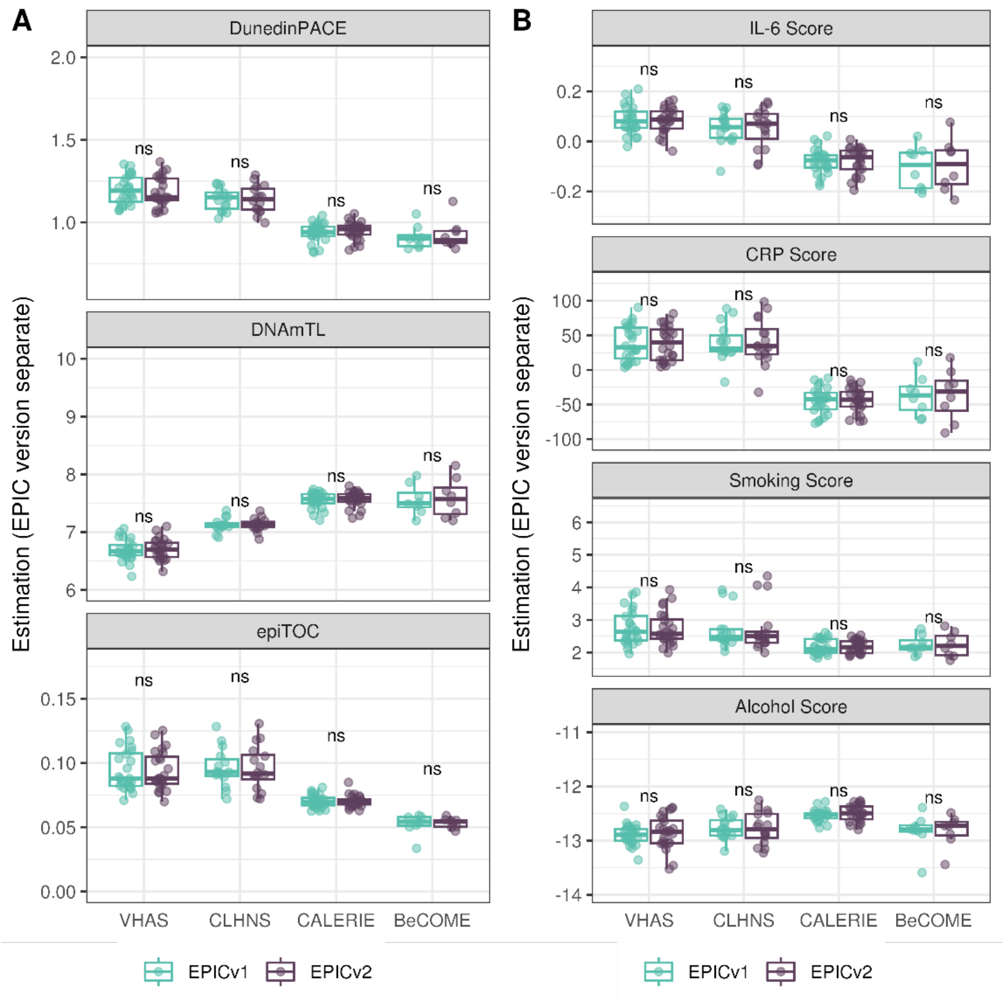


**Supplementary Figure 25.** DunedinPACE, DNAmTL, epiTOC, and DNA methylation-based inflammation and lifestyle biomarkers calculated by considering EPIC versions separately after BMIQ normalization with batch-correction for EPIC version, chip and row in VHAS, CLHNS, CALERIE, and BeCOME. **(A)** rate-based and other clocks **(B)**. DNA methylation-based inflammation and lifestyle biomarkers. Paired t-tests were performed to compare estimates between EPICv1 and EPICv2, and p-values were derived. Statistical significance was defined as Bonferroni adjusted *p*-value <0.05. ** denotes Bonferroni *p* <0.05, *** denotes Bonferroni *p* <0.001, “ns” denotes “not significant”, and “d” denotes effect size measured using Cohen’s d. A positive Cohen’s d indicates estimates in EPICv2 compared to EPICv1.

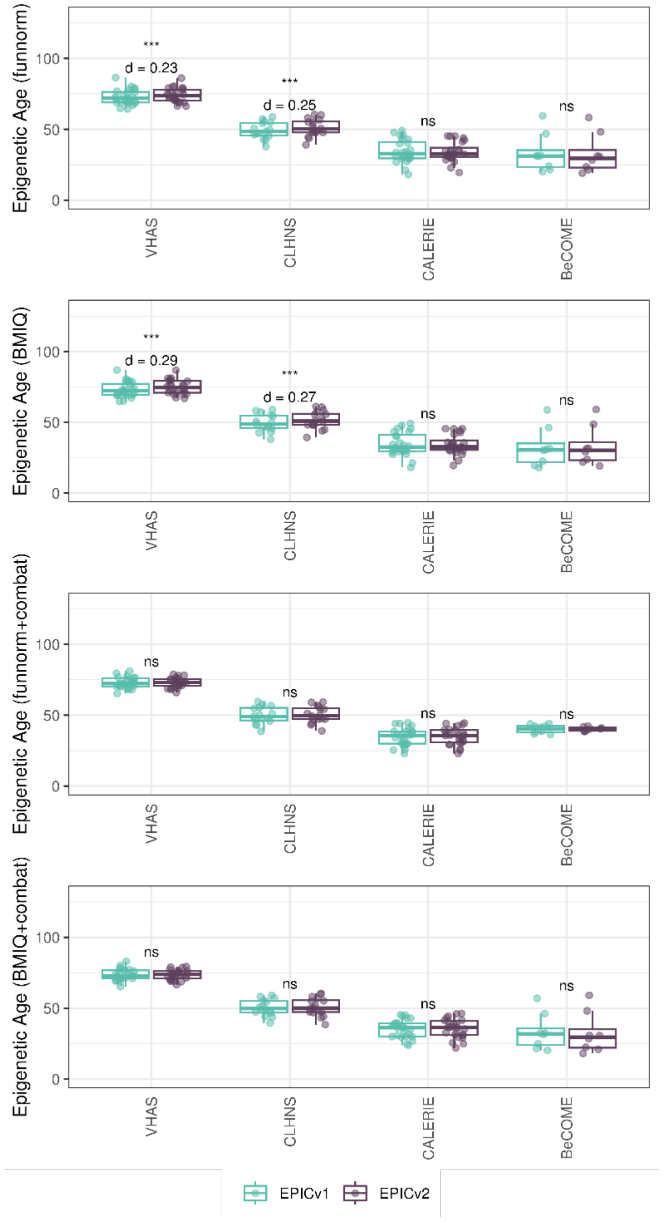


**Supplementary Figure 26.** Differences in epigenetic ages between EPICv1 and EPICv2 using the Garman and Quintela-Fandino clock in VHAS, CLHNS, and CALERIE when using functional normalization; normalizing EPICv1 and EPICv2 together using BMIQ normalization; functional normalization with batch-correction for EPIC version, chip and row and normalizing EPICv1 and EPICv2 together using BMIQ normalization with batch-correction for EPIC version, chip and row. Paired t-tests were performed to compare estimates between EPICv1 and EPICv2, and p-values were derived. Statistical significance was defined as Bonferroni adjusted p-value <0.05. ** denotes Bonferroni p <0.05, *** denotes Bonferroni p<0.001, "ns" denotes "not significant", and "d" denotes effect size measured using Cohen's d. A positive Cohen's d indicates estimates in EPICv2 compared to EPICv1.

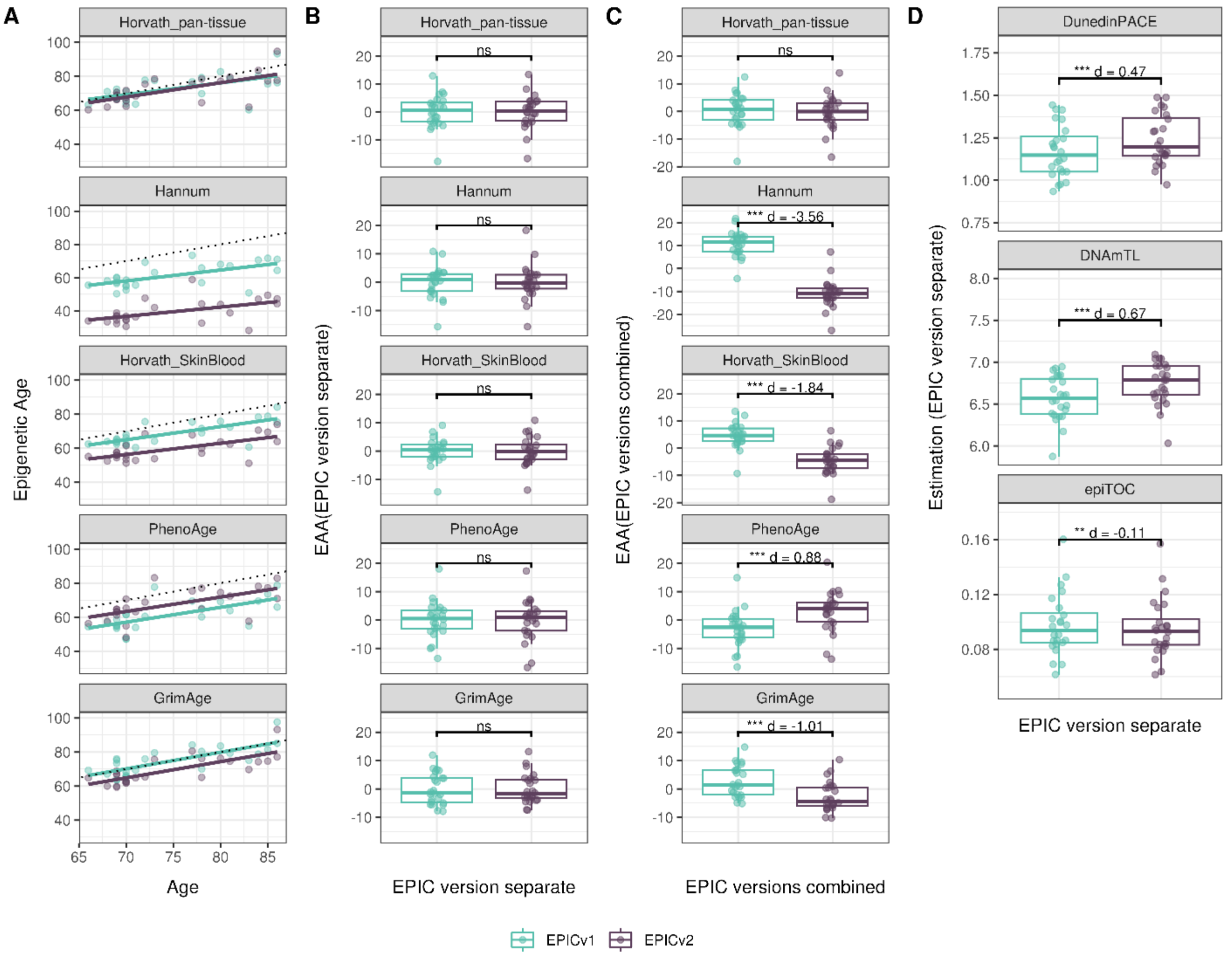


**Supplementary Figure 27.** Epigenetic age estimated on matched samples assessed on EPICv1 and EPICv2 in VHAS capillary blood samples using functional normalization. **(A)** Scatter plot of Horvath pan-tissue, Hannum, Horvath skin and blood, PhenoAge, and GrimAge clock ages (Y axis) and chronological age (X axis) with dotted line indicating x=y, coloured by EPIC version. **(B - C)** Boxplots comparing EPICv1 and EPICv2 EAAs calculated by considering EPIC versions separately and combined, respectively. **(D)** Boxplots comparing DunedinPACE, DNAmTL and epiTOC estimates calculated by considering EPIC versions separately. Paired t-tests were performed to compare estimates between EPICv1 and EPICv2, and p-values were derived. Statistical significance was defined as Bonferroni adjusted p-value <0.05. ** denotes Bonferroni p <0.05, *** denotes Bonferroni p<0.001, “ns” denotes “not significant”, and “d” denotes effect size measured using Cohen’s d. A positive Cohen’s d indicates higher estimates in EPICv2 compared to EPICv1.

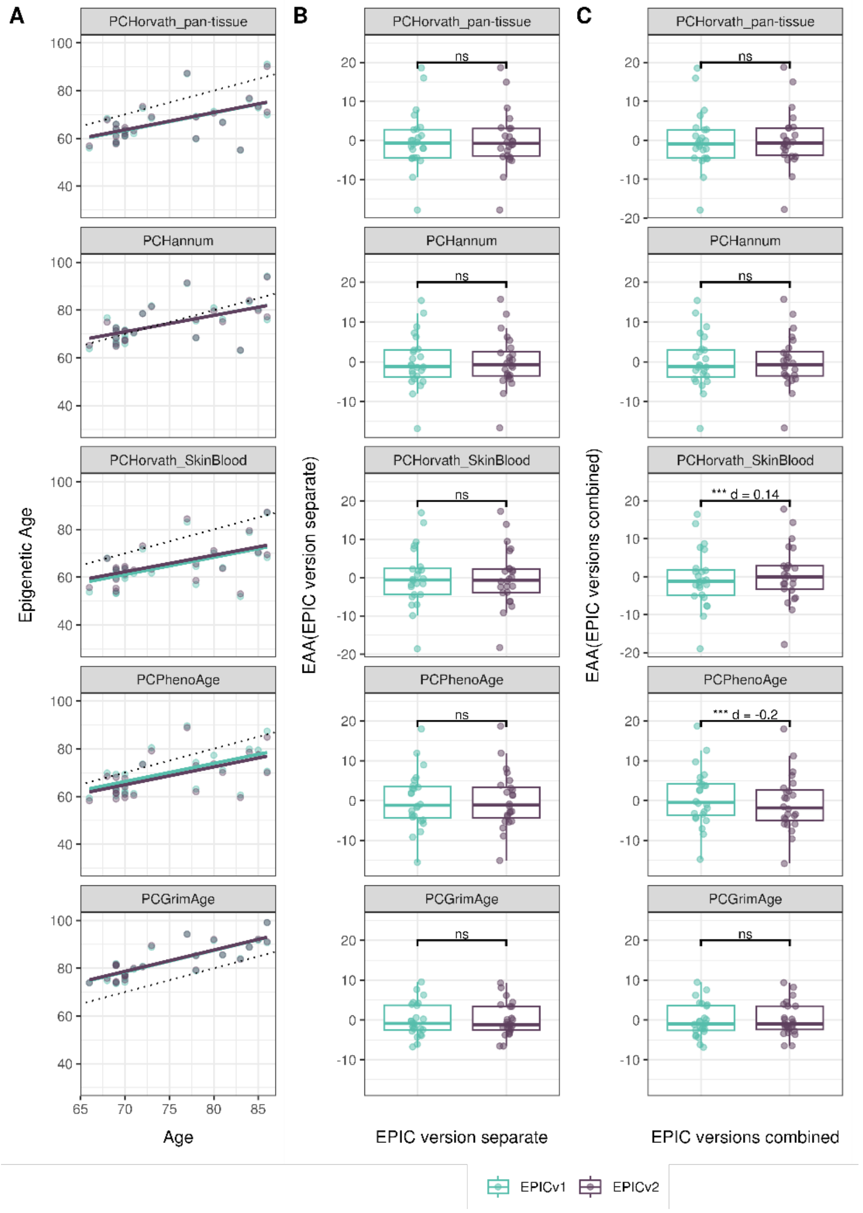


**Supplementary Figure 28.** Epigenetic age estimated on matched samples assessed on EPICv1 and EPICv2 in VHAS capillary blood samples using functional normalization. **(A)** Scatter plot of PC clocks of Horvath pan-tissue, Hannum, Horvath skin and blood, PhenoAge, and GrimAge clock ages (Y axis) and chronological age (X axis) with dotted line indicating x=y, coloured by EPIC version. **(B - C)** Boxplots comparing EPICv1 and EPICv2 EAAs calculated by considering EPIC versions separately and combined, respectively. Paired t-tests were performed to compare estimates between EPICv1 and EPICv2, and p-values were derived. Statistical significance was defined as Bonferroni adjusted p-value <0.05. ** denotes Bonferroni p <0.05, *** denotes Bonferroni p<0.001, "ns" denotes "not significant", and "d" denotes effect size measured using Cohen's d. A positive Cohen's d indicates higher estimates in EPICv2 compared to EPICv1.

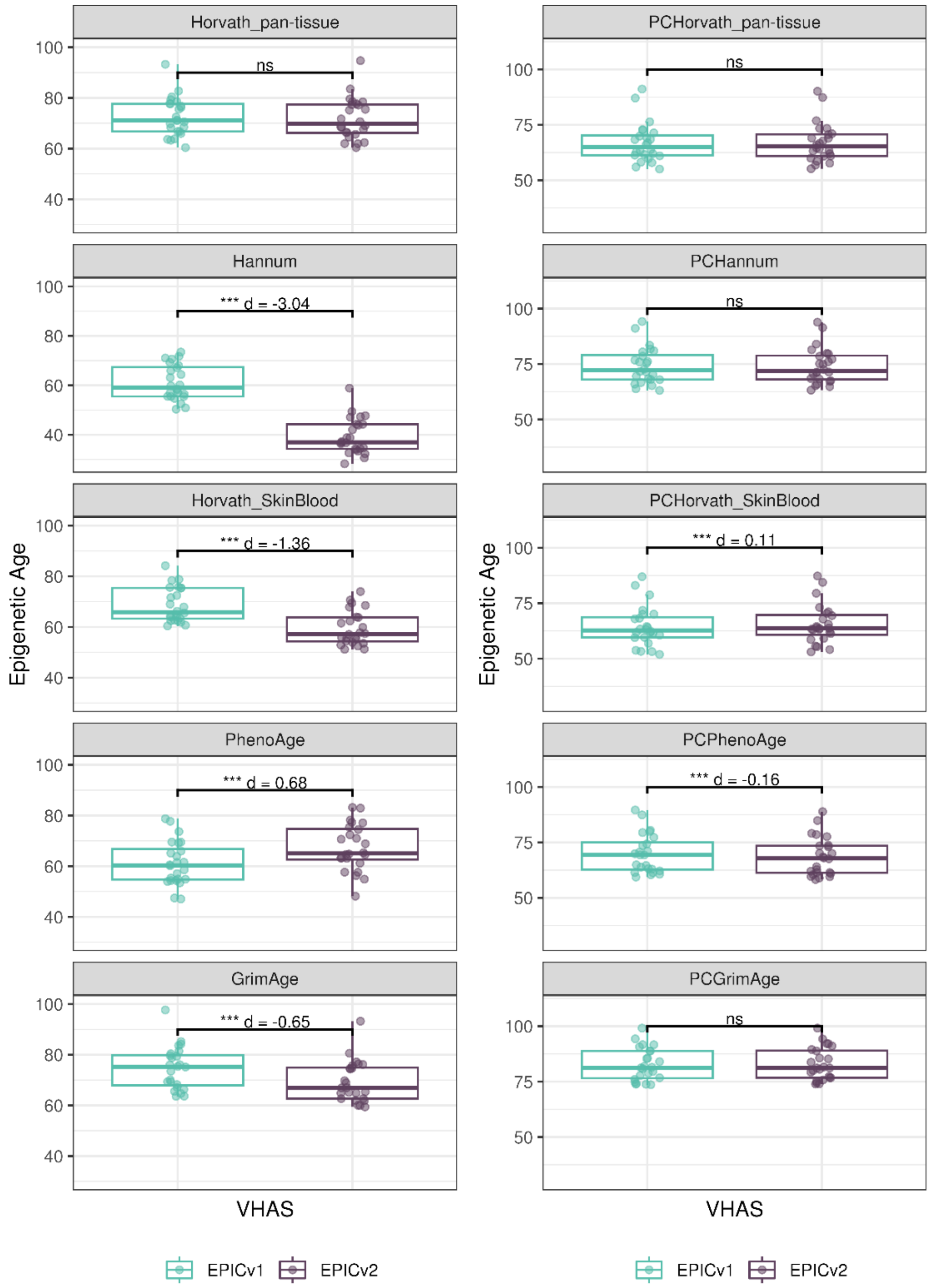


**Supplementary Figure 29.** Differences in epigenetic ages between EPICv1 and EPICv2 using the Horvath pan-tissue, Hannum, Horvath skin and blood, PhenoAge, GrimAge clocks, and PC clocks in VHAS capillary blood samples using functional normalization.

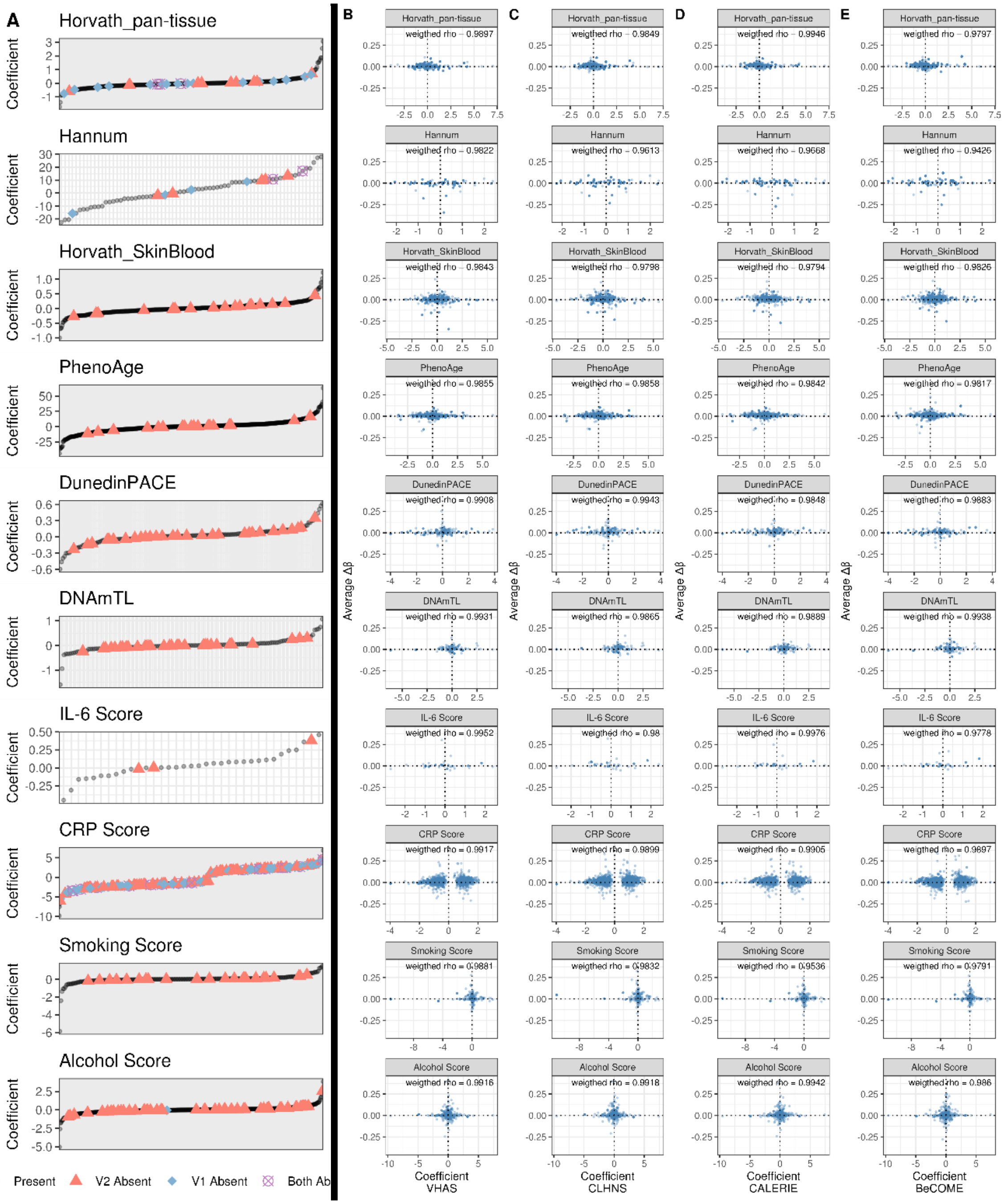


**Supplementary Figure 30. (A)** Clock and biomarker predictor CpGs absent in EPICv1 and EPICv2 and their corresponding coefficients. Both Ab: probe are absent in both EPICv1 and EPICv2. **(B-C)** Average difference in beta values (EPICv2 β - EPICv1 β) of clock and biomarker predictor CpGs coefficients in **(B)** VHAS, **(C)** CLHNS, **(D)** CALERIE, **(E)** BeCOME. Spearman correlation between EPICv1 and EPICv2 beta values of clock and predictor CpGs, weighted by corresponding coefficients are provided. IDOL CpGs, which do not have coefficients, and epiTOC CpGs, which have equal coefficients are not shown here.

113

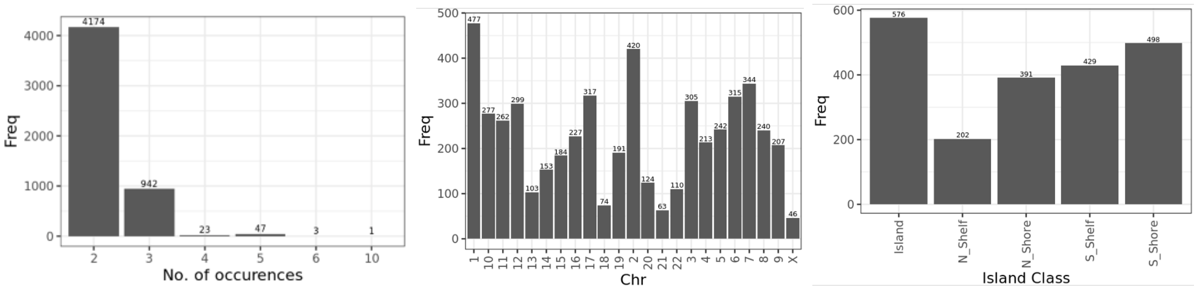


A B C

**Supplementary Figure 31:** Frequency of occurrence of replicate probes on EPIC v2 **(A)**, their distribution across chromosomes **(B)**, and CpG island classes **(C)**.

**(A) VHAS** **(B) CLHNS** **(C) CALERIE**

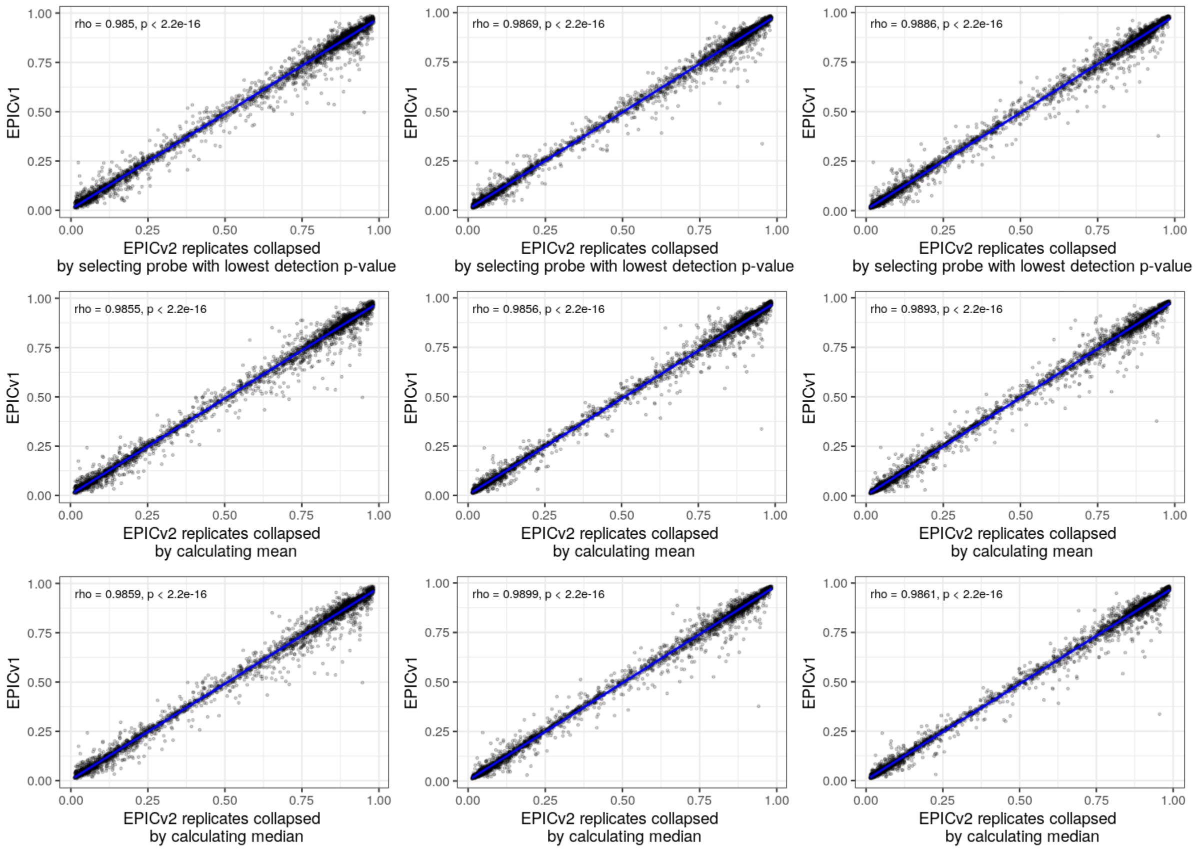


**Supplementary Figure 32:** Spearman correlation between β values of EPICv2 replicate probes and corresponding EPICv1 probes (n=3602). EPICv2 replicate probes were collapsed to obtain a single β value using detection *p*-value (row 1), mean (row 2), and median (row 3) based strategies for (A) VHAS, (B) CLHNS, and (C) CALERIE.
