## Supplementary material for "Accounting for differences between Infinium MethylationEPIC v2 and v1 in DNA methylation-based tools"

### High coverage of epigenetic clock, biomarker predictor, and cell type deconvolution CpGs on EPICv2

To get a first sense of the applicability of widely used DNA methylation-based tools, which were built on previous arrays/versions, on EPICv2, we looked in silico at the percentage of predictive CpGs which are absent in either of the EPIC versions and which are replicated on EPICv2. Across the seven epigenetic clocks investigated, we observed that ~77-96% predictive CpGs were retained on EPICv2, with a reintroduction to EPICv2 of several predictive CpGs which were absent in EPICv1 but present on the 450K (Horvath pan-tissue: 14 CpGs; Hannum: 4 CpGs; and epigenetic timer of cancer (epiTOC): 25 CpGs). In all the clocks we examined, ~1-7% of the predictive clock CpGs had replicate probes (replicate probes consist of 1% of total EPICv2 probes), except for epiTOC which did not contain replicate probes.
Similarly, across the four common biomarker predictors that we investigated here, namely IL-6, CRP, smoking and alcohol scores, we noted ~89-94% coverage of predictive CpGs on EPICv2, with 0.86-2.86% also including replicate probes. In the CRP predictor, we additionally observed that 5.84% (103 probes) predictive CpGs were absent in EPICv1, with 83 of these CpGs reintroduced in EPICv2, while there was no absent CpGs in EPICv1 in the other three predictors.
Next, on examining the 1200 pre-selected extended Identifying Optimal DNA methylation Libraries reference (IDOL) probes used in cell type deconvolution(Salas et al. 2022), we noted that 99.33% were represented on EPICv2, with 4% also having replicate probes on EPICv2 (Table 2). On estimating the correlation of β values of predictive CpGs employed by these DNA methylation-based tools between EPICv1 and EPICv2, weighted by corresponding CpG coefficients, we identified high Spearman correlation ranging from 0.9615-0.9952 (Supplementary Figure 30). We also noted that mean absolute differences in β values of these predictive CpGs between the EPIC versions range from 0.0157-0.0342 and pool SD from 0.0267-0.0658, with the Horvath pan-tissue clock having the lowest mean absolute difference between the two versions (Supplementary Table S13).

### Characterization of replicate probes in EPICv2 and comparing strategies for collapsing EPICv2 replicate probes

In total, there are 11,529 replicate probes on EPICv2 after excluding probes on chromosomes 0 and M, which consist of 1.23% of the total probes. These replicates are distributed across chromosomes 1-22 and X, and are predominantly located within CpG islands, consistent with the intended increased coverage of these genomic regions on EPICv2 (Supplementary Figure 31). Over 80% (4,174) of these probes have two replicates each, with the largest number of replicates being ten (1 in 5,190, 0.02%).

Given that predictive CpGs of several DNA methylation-based tools include EPICv2 replicate probes to varying extent, even some with large coefficient weights contributing substantially to the generated estimates, there is a need to collapse replicates to a single representative beta value analogous to corresponding probes on EPICv1. While significantly high correlation among replicate probes is expected, residual methylation differences between replicate probes of varying designs have been reported(Kaur et al. 2023). Choosing the replicate with lowest detection p-value as the representative probe has been suggested as a way to collapse replicates(Kaur et al. 2023), however this strategy has not been compared to other methods. To that end, we compared three strategies to collapse replicates into a single $\beta$ value per locus: (i) previously suggested method of choosing the replicate with lowest detection *p*-value(Kaur et al. 2023), (ii) estimating mean of all replicates mapping to a genomic locus, and (iii) estimating median of all replicates mapping to a genomic locus, and compared representative EPICv2 replicate beta values (3602 probes) thus obtained to respective EPICv1 probes. We also noted that all three methods showed significantly high correlation with EPICv1 in all four cohorts: VHAS (Spearman *rho* values by strategy-detection *p*-value-based: 0.9886, mean-based: 0.9893, median-based: 0.9899), CLHNS (Spearman *rho* values by strategy-detection P-value-based: 0.9850, mean-based: 0.9855, median-based: 0.9859), and CALERIE (Spearman *rho* values by strategy-detection *p*-value-based: 0.9869, mean-based: 0.9856, median-based: 0.9861) (Supplementary Figure 32). Comparing the three strategies to one another, we observed negligible absolute mean differences in average beta values of EPICv2 replicate probes across samples, ranging from 0.0002 - 0.0078 across the four cohorts.

**Epigenetic ages were significantly different between EPIC version using a newly developed epigenetic clock trained on the common probe sets of 450K, EPICv1 and EPICv2**

A newer epigenetic clock compatible with EPICv2 was recently developed to address the issue of absent probes on EPICv2, using the common probesets of 450K, EPICv1 and EPICv2 (Garma and Quintela-Fandino 2024). While the clock was tested on publicly available 450K and EPICv1 data, it was not directly validated on EPICv2 and its performance on EPICv1 and EPICv2 was not compared. On testing this clock on our four cohorts, we identified Pearson correlation ranging from -0.499 to 0.843 between chronological age and epigenetic age obtained from both EPICv1 and EPICv2, with mean absolute error (MAE) from 3 to 22.69 years (Supplementary Table S16) depending on cohort. EPICv2 epigenetic ages were highly correlated with EPICv1 epigenetic ages in three cohorts (0.974-0.996), but not significantly correlated in CALERIE ($r = -0.056$) (Supplementary Table S16). On comparing clock estimates between paired EPICv1 and EPICv2 samples, we noted significant differences in epigenetic ages between EPICv2 and EPICv1 in VHAS and CLHNS, with these differences ranging in effect size from 0.23-0.25 (Supplementary Table S16 and Supplementary Figure S26).
