## Supplementary Tables for "Accounting for differences between Infinium MethylationEPIC v2 and v1 in DNA methylation-based tools"

**Table of contents**

**Supplementary Table 1.** Summary of within and between EPICv1 and EPICv2 technical replicates Spearman correlation, RMSE, and ICC.
**Supplementary Table 2.** Summary of spearman correlations, RMSE between EPICv1 and EPICv2 on array and probe level, and pooled standard deviation on probe levels (721,378 shared probes)
**Supplementary Table 3.** Summary of spearman correlations between EPICv1 and EPICv2 and pooled standard deviation on probe level.
**Supplementary Table 4.** Summary of probes with spearman correlations ≤ 0.7 between EPICv1 and EPICv2 and pooled standard deviation over the first quartile on probe level using different levels of preprocessing
**Supplementary Table 5.** Probes shared among 450K, EPICv1 and EPICv2, and with Spearman correlation ≤ 0.7 between EPICv1 and EPICv2 in the four cohorts and overlap with previously published references comparing 450K and EPICv1.
**Supplementary Table 6.** Summary of probe counts using various Spearman correlations and pooled standard deviation thresholds.
**Supplementary Table 7.** Summary of spearman correlations between EPICv1 and EPICv2 and pooled standard deviation on probe levels of the 2,169 probes in cell type deconvolution panels.
**Supplementary Table 8.** Commonly identified low concordance probes with Spearman rho correlation ≤ 0.70(calculated by each cohort) and pool SD more than the lower quartile pooled SD (calculated by each cohort), commonly across VHAS, CLHNS, CALERIE, BeCOME
**Supplementary Table 9.** Overlap of probes based on Spearman correlation and pooled standard deviation thresholds and probes with low quality and previously published references.
**Supplementary Table 10.** Paired t-tests comparing DNA methylation-based immune cell type proportions estimated using EPICv1 and EPICv2 with IDOL probes independently for each cohort. Spearman correlation (rho), paired t-test p-values and Bonferroni adjusted p-values, and Cohen's d effect size between EPICv1 and EPICv2 estimates obtained from matched samples in each cohort are provided.
**Supplementary Table 11.** Paired t-tests comparing DNA methylation-based immune cell type proportions estimated using EPICv1 and EPICv2 with auto selected probes, independently for each cohort. Spearman correlation (rho), paired t-test p-values and Bonferroni adjusted p-values, and Cohen's d effect size between EPICv1 and EPICv2 estimates obtained from matched samples in each cohort are provided.
**Supplementary Table 12**. Pearson correlations of epigenetic clock estimates with chronological age in EPICv1 and EPICv2 samples of VHAS, CLHNS, CALERIE and BeCOME.
**Supplementary Table 13.** Comparison of epigenetic ages between EPICv1 and EPICv2 in first-, second-, rate-based and other epigenetic clocks.
**Supplementary Table 14.** Pearson correlation (r) and paired T tests results of EAAs of between EPICv1 and EPICv2 in Horvath pantissue, Hannum, Horvath SkinBlood, PhenoAge, and GrimAge.
**Supplementary Table 15.** Pearson correlation (r) and paired T tests results of EPICv1 and EPICv2 comparisons for DunedinPACE, DNAmTL and epiTOC.
**Supplementary Table 16.** Comparison of DNA methylation-based predictor estimations between EPv1 and EPICv2.

**Supplementary Table 17.** Pearson correlations of Garma and Quintela-Fandino's epigenetic clock estimates with chronological age in EPICv1 and EPICv2 samples of VHAS, CLHNS, CALERIE and BeCOME.

**Supplementary Table 18.** Illumina's 17 quality control metrics, detection p-value, beadcount, average methylated and unmethylated intensity values estimated based on EPICv1 and EPICv2 DNA methylation data. For all three cohorts, values for each metric are provided as mean (standard deviation) or percentages separated by array version.

**Supplementary Table 1. Summary of within and between EPICv1 and EPICv2 technical replicates Spearman correlation, RMSE, and ICC.**

| Within EPIC version | | | | | | |
|---|---|---|---|---|---|---|
| Cohort | VHAS | | CLHNS* | | CALERIE | |
| | EPICv1 | EPICv2 | EPICv1 | EPICv2 | EPICv1 | EPICv2 |
| Spearman rho (range) | 0.9909(0.9902-0.9916) | 0.9863(0.9858-0.9869) | NA | 0.9917 | 0.9908(0.9907-0.9909) | 0.9865(0.9860-0.9870) |
| RMSE (range) | 0.0226(0.0214-0.0238) | 0.0260(0.0256-0.0264) | NA | 0.0232 | 0.0216(0.0208-0.0223) | 0.0231(0.0224-0.0239) |
| ICC (range) | 0.9979(0.9977-0.9981) | 0.9973(0.9972-0.9974) | NA | 0.9980 | 0.9981(0.9979-0.9983) | 0.9978(0.9977-0.998) |
| Between EPICv1 and EPICv2 | | | | | | |
| Cohort | VHAS | | CLHNS* | | CALERIE | |
| Spearman rho (range) | 0.9756(0.9737-0.9774) | | NA | | 0.9757(0.9756-0.9758) | |
| RMSE (range) | 0.0401(0.0386-0.0416) | | | | 0.0371(0.0367-0.0375) | |
| ICC (range) | 0.9949(0.9947-0.9951) | | | | 0.9956(0.9955-0.9958) | |

RMSE: root mean square root error; ICC: intra-class correlation coefficients; *CLHNS did not have technical replicate in EPICv1

**Supplementary Table 2. Summary of spearman correlations, RMSE between EPICv1 and EPICv2 on array and probe level, and pooled standard deviation on probe levels (721,378 shared probes)**

| Array level | | | | |
|---|---|---|---|---|
| | **VHAS** | **CLHNS** | **CALERIE** | **BeCOME** |
| Mean array level Spearman rho (range;sd) | 0.9761(0.9688-0.9803; sd: 0.003) | 0.9759(0.9729-0.9792; sd: 0.002) | 0.976(0.9682-0.9807; sd: 0.0029) | 0.9737(0.9684-0.9766; sd: 0.0028) |
| Mean array level RMSE | 0.0384(0.0362-0.0421; sd: 0.0014) | 0.0401(0.0375-0.0434; sd: 0.0015) | 0.0362(0.0336-0.0436; sd: 0.0023) | 0.0463(0.0417-0.0532; sd: 0.0034) |
| **Probe level** | | | | |
| | **VHAS** | **CLHNS** | **CALERIE** | **BeCOME** |
| Mean probe level Spearman rho | 0.4109(-0.6484-0.9914; sd: 0.3483) | 0.3547(-0.9071-1; sd: 0.3878) | 0.3863(-0.7443-0.9957; sd: 0.3117) | 0.2464(-1-1; sd: 0.4311) |
| Mean probe level pooled standard deviation | 0.0305(0.001-0.3751; sd: 0.0297) | 0.0235(7e-04-0.3675; sd: 0.0199) | 0.0253(9e-04-0.3586; sd: 0.0246) | 0.0261(4e-04-0.4121; sd: 0.0271) |
| Mean probe level RMSE | 0.0297(0.0013-0.9303; sd: 0.0245) | 0.0314(0.001-0.9434; sd: 0.0251) | 0.0283(0.0011-0.9296; sd: 0.0227) | 0.033(4e-04-0.9128; sd: 0.0326) |
| First lower quartile pooled SD (threshold) | 0.0101 | 0.0099 | 0.0111 | 0.0101 |

RMSE: root mean square root error; sd: standard deviation

**Supplementary Table 3. Summary of spearman correlations between EPICv1 and EPICv2 and pooled standard deviation on probe level.**

| Cohort | VHAS | | CLHNS | |
|---|---|---|---|---|
| | Spearman correlation ≤ 0.7 | Spearman correlation >0.7 | Spearman correlation ≤ 0.7 | Spearman correlation >0.7 |
| Probe count (% of EPICv1 and EPICv2 shared probes) | 510533(70.7719 %) | 210845(29.2281 %) | 541285(75.0349 %) | 180093(24.9651 %) |
| Mean Spearman rho | 0.2343 (-0.6484-0.7, sd: 0.2499) | 0.8386 (0.7001-0.9914, sd: 0.0731) | 0.1951 (-0.9071-0.7, sd: 0.3108) | 0.8344 (0.7036-1, sd: 0.0741) |
| Mean pooled standard deviation | 0.0178 (0.001-0.3465, sd: 0.0159) | 0.0614 (0.003-0.3751, sd: 0.0325) | 0.0166 (7e-04-0.3318, sd: 0.0135) | 0.0441 (0.0011-0.3675, sd: 0.0217) |
| t-test results on comparing probe level pooled standard deviation | *t(253249)= -588.43, p < 2.2e-16* | | *t(228527) = -507.51, p < 2.2e-16* | |
| Cohort | **CALERIE** | | **BeCOME** | |
| | Spearman correlation ≤ 0.7 | Spearman correlation >0.7 | Spearman correlation ≤ 0.7 | Spearman correlation >0.7 |
| Probe count (% of EPICv1 and EPICv2 shared probes) | 582244(80.7127 %) | 139134(19.2873 %) | 592955(82.1975 %) | 128423(17.8025 %) |
| Mean Spearman rho | 0.283 (-0.7443-0.7, sd: 0.2525) | 0.8189 (0.7009-0.9957, sd: 0.0713) | 0.1196 (-1-0.6905, sd: 0.3668) | 0.8317 (0.7143-1, sd: 0.0794) |
| Mean pooled standard deviation | 0.0184 (9e-04-0.3349, sd: 0.0144) | 0.054 (0.0014-0.3586, sd: 0.0354) | 0.0206 (4e-04-0.3943, sd: 0.0193) | 0.0515 (4e-04-0.4121, sd: 0.0402) |
| t-test results on comparing probe level pooled standard deviation | *t(150243) = -367.40, p < 2.2e-16* | | *t(141500) = -268.79, p < 2.2e-16* | |
| | **Common across four cohorts** | | | |
| Probe count (% of EPICv1 and EPICv2 shared probes) | Spearman correlation ≤ 0.7 | Spearman correlation >0.7 | | |
| Mean Spearman rho | 432521(59.9576 %) | 56289(7.803%) | | |

**Supplementary Table 4. Summary of probes with spearman correlations ≤ 0.7 between EPICv1 and EPICv2 and pooled standard deviation over the first quartile on probe level using different levels of preprocessing**

| Spearman correlation ≤ 0.7 (percentage relative to the shared 721,378 EPICv1 and EPICv2 probes) | | | | | |
|---|---|---|---|---|---|
| Data processing method | Raw | Funnorm without background and without dye bias correction | Funnorm | Funnorm + Combat | Common to all four levels of preprocessing |
| VHAS | 546889(75.8117%) | 518771(71.9139%) | 510533(70.7719%) | 544111(75.4266%) | 502848(69.7066%) |
| CLHNS | 626724(86.8787%) | 545023(75.553%) | 541285(75.0349%) | 541285(75.0349%) | 522431(72.4213%) |
| CALERIE | 619432(85.8679%) | 595741(82.5837%) | 582244(80.7127%) | 637503(88.3729%) | 560359(77.679%) |
| BeCOME | 596614(82.7048%) | 598543(82.9722%) | 592955(82.1975%) | 719271(99.7079%) | 542858(75.2529%) |
| Common to all four cohorts | 474901(65.8325%) | 447945(62.0957%) | 432521(59.9576%) | 484827(67.2085%) | |

| Spearman correlation > 0.7 | | | | | |
|---|---|---|---|---|---|
| Data processing method | Raw | Funnorm without background and dye bias correction | Funnorm | Funnorm and Combat | Common to all four levels of preprocessing |
| VHAS | 174299(24.162%) | 202607(28.0861%) | 210845(29.2281%) | 177267(24.5734%) | 156253(21.6604%) |
| CLHNS | 94302(13.0725%) | 176355(24.447%) | 180093(24.9651%) | 180093(24.9651%) | 84206(11.6729%) |
| CALERIE | 101459(14.0646%) | 125637(17.4163%) | 139134(19.2873%) | 83875(11.6271%) | 62084(8.6063%) |
| BeCOME | 124571(17.2685%) | 122835(17.0278%) | 128423(17.8025%) | 2107(0.2921%) | 2034(0.282%) |
| Common to all four cohorts | 29717(4.1195%) | 53931(7.4761%) | 56289(7.803%) | 1496(0.2074%) | |

**Supplementary Table 5. Probes shared among 450K, EPICv1 and EPICv2, and with Spearman correlation ≤ 0.7 between EPICv1 and EPICv2 in the four cohorts and overlap with previously published references comparing 450K and EPICv1.**

| **Cohort** | **Our analyses** | **Sugden et al. 2020 (n=350)** | **Logue et al. 2017 (n=145)** | **Olstad et al. 2022 (n=17)** | **Shared among the three reference lists** |
|---|---|---|---|---|---|
| | Spearman rho ≤ 0.7 | Pearson correlation ≤ 0.7: 281193 probes | Pearson correlation ≤ 0.7: 337956 probes | ICC <0.4: 239848 probes | total probes: 187311 |
| VHAS | 266840 | 250054 (88.93%) | 263912 (78.09%) | 175388 (73.12%) | 165616 (88.42%) |
| CLHNS | 285917 | 256162 (91.1%) | 279259 (82.63%) | 186671 (77.83%) | 169245 (90.36%) |
| CALERIE | 303595 | 271574 (96.58%) | 299599 (88.65%) | 201596 (84.05%) | 180258 (96.23%) |
| BeCOME | 304693 | 261216 (92.9%) | 295894 (87.55%) | 201263 (83.91%) | 173422 (92.59%) |

ICC: intraclass correlation coefficient; percentage of probes identified as low concordant in both our analysis and previous studies (reference lists), relative to the total number of low concordant probes in the reference list.

**Supplementary Table 6. Summary of probe counts using various Spearman correlations and pooled standard deviation thresholds.**

| Thresholds | VHAS | CLHNS | CALERIE | BeCOME | Common across four cohorts** |
|---|---|---|---|---|---|
| Spearman correlation > 0.7 and pooled SD < threshold* (percent over probes with Spearman correlation >0.7; percent over 721378 probes) | 146 (0.069%; 0.02%) | 886 (0.492%; 0.123%) | 770 (0.553%; 0.107%) | 7162 (5.577%; 0.993%) | 0 |
| Spearman correlation > 0.7 and pooled SD ≥ threshold (percent over probes with Spearman correlation > 0.7; percent over 721378 probes) | 210699 (99.931%; 29.208%) | 179207 (99.508%; 24.842%) | 138364 (99.447%; 19.181%) | 121261 (94.423%; 16.81%) | 56227 (7.794%) |
| Spearman correlation ≤ 0.7 and pooled SD < threshold (percent over probes with Spearman correlation ≤0.7; percent over 721378 probes) | 180199 (35.296%; 24.98%) | 179459 (33.154%; 24.877%) | 179575 (30.842%; 24.893%) | 173183 (29.207%; 24.007%) | 103326 (14.323%) |
| Spearman correlation ≤ 0.7 and pooled SD ≥ threshold (poorly-concordant) (percent over probes with Spearman correlation ≤0.7; percent over 721378 probes) | 330334 (64.704%; 45.792%) | 361826 (66.846%; 50.158%) | 402669 (69.158%; 55.819%) | 419772 (70.793%; 58.19%) | 197208 (27.338%) |

| Cohort | VHAS | | CLHNS | |
|---|---|---|---|---|
| Thresholds | Correlation > threshold or pooled SD < threshold | Correlation ≤ threshold and pooled SD ≥ threshold (poorly-concordant ) | Correlation > threshold or pooled SD < threshold | Correlation ≤ threshold and pooled SD ≥ threshold (poorly-concordant ) |
| Spearman correlation: 0.7; pooled SD: lower quartile pooled SD for each cohort | 391044 (54.208%)** | 330334 (45.792%) | 359552 (49.842%) | 361826 (50.158%) |
| Spearman correlation: 0.7; pooled SD: 0.05 | 699675 (96.991%) | 21703 (3.009%) | 708661 (98.237%) | 12717 (1.763%) |
| Spearman correlation: 0.5; pooled SD: lower quartile pooled SD for each cohort | 488670 (67.741%) | 232708 (32.259%) | 462117 (64.06%) | 259261 (35.94%) |

| | | | | |
|---|---|---|---|---|
| Spearman correlation: 0.5; pooled SD: 0.05 | 713108 (98.854%) | 8270 (1.146%) | 715392 (99.17%) | 5986 (0.83%) |
| Cohort | CALERIE | | BeCOME | |
| Thresholds | Correlation > threshold or pooled SD < threshold | Correlation ≤ threshold and pooled SD ≥ threshold (poorly-concordant ) | Correlation > threshold or pooled SD < threshold | Correlation ≤ threshold and pooled SD ≥ threshold (poorly-concordant ) |
| Spearman correlation: 0.7; pooled SD: lower quartile pooled SD for each cohort | 318709 (44.181%) | 402669 (55.819%) | 301606 (41.81%) | 419772 (58.19%) |
| Spearman correlation: 0.7; pooled SD: 0.05 | 702485 (97.381%) | 18893 (2.619%) | 681275 (94.441%) | 40103 (5.559%) |
| Spearman correlation: 0.5; pooled SD: lower quartile pooled SD for each cohort | 445492 (61.756%) | 275886 (38.244%) | 395842 (54.873%) | 325536 (45.127%) |
| Spearman correlation: 0.5; pooled SD: 0.05 | 713172 (98.862%) | 8206 (1.138%) | 697530 (96.694%) | 23848 (3.306%) |
| Cohort | common | | | |
| Thresholds | Correlation > threshold or pooled SD < threshold | Correlation ≤ threshold and pooled SD ≥ threshold (poorly-concordant ) | | |
| Spearman correlation: 0.7; pooled SD: lower quartile pooled SD for each cohort | 167533 (23.224%) | 197208 (27.338%) | | |
| Spearman correlation: 0.7; pooled SD: 0.05 | 653063 (90.53%) | 1930 (0.268%) | | |
| Spearman correlation: 0.5; pooled SD: lower quartile pooled SD for each cohort | 250839 (34.772%) | 93530 (12.965%) | | |
| Spearman correlation: 0.5; pooled SD: 0.05 | 684694 (94.915%) | 704 (0.098%) | | |

* Lower quartile pooled SD threshold for each cohort: VHAS: 0.0101; CLHNS: 0.0100; CALERIE: 0.0111; BeCOME: 0.0101

** Percent calculated over 721378 probes

**Supplementary Table 7. Summary of spearman correlations between EPICv1 and EPICv2 and pooled standard deviation on probe levels of the 2,169 probes in cell type deconvolution panels.**

| | **VHAS** | **CLHNS** | **CALERIE** | **BeCOME** |
|---|---|---|---|---|
| **Probe level Spearman correlation** | | | | |
| I. Mean correlation across all shared probes | 0.4109 | 0.3547 | 0.3863 | 0.2464 |
| II. Mean correlation across cell type probes | 0.5815 | 0.5352 | 0.5338 | 0.3729 |
| t-statistics comparing I and II | t(2181.4)= 29.866 | t(2180)= 27.499 | t(2178.3)= 22.839 | t(2174.2)= 14.749 |
| **probe level pooled standard deviation (variability)** | | | | |
| I. Mean variability across all shared probes | 0.0305 | 0.0235 | 0.0253 | 0.0261 |
| II. Mean variability across cell type probes | 0.0374 | 0.0299 | 0.0307 | 0.0306 |
| t-statistics comparing I and II | t(2177.6)= 12.791 | t(2173.8)= 15.919 | t(2179.9)= 13.000 | t(2182.5)= 10.464 |

**Supplementary Table 8. Commonly identified low concordance probes with Spearman rho correlation ≤ 0.70(calculated by each cohort) and pool SD more than the lower quartile pooled SD (calculated by each cohort), commonly across VHAS, CLHNS, CALERIE, BeCOME**

Please see a separate table. This large table is also publically available on GitHub: https://github.com/kobor-lab/EPICv1v2_comparison_manuscript/blob/main/Supplementary_Table8.csv

**Supplementary Table 9. Overlap of probes based on Spearman correlation and pooled standard deviation thresholds and probes with low quality and previously published references.**

| | | | | | | |
|---|---|---|---|---|---|---|
| **Spearman correlation ≤0.7 and pooled SD ≥ lower quartile pooled SD (low concordance probes)** | | | | | | |
| Cohort | | VHAS | CLHNS | CALERIE | BeCOME | Common across four cohorts |
| Probe count* | | 330334 (100%) | 361826 (100%) | 402669 (100%) | 419772 (100%) | 197208 (100%) |
| High-confidence mapping probes in cell line (539313 probes) | | 217874 (65.96%) | 243546 (67.31%) | 279166 (69.33%) | 290844 (69.29%) | 123969 (62.86%) |
| Flagged detectionP and beadcount probes** | | 30987 (9.38%) | 14062 (3.89%) | 19328 (4.8%) | 12444 (2.96%) | 430 (0.22%) |
| Previously published references | EPICv2 design type switch probes (82 probes) | 27 (0.01%) | 34 (0.01%) | 40 (0.01%) | 40 (0.01%) | 16 (0.01%) |
| | Peters reference (11878 probes) | 319 (0.1%) | 342 (0.09%) | 336 (0.08%) | 313 (0.07%) | 220 (0.11%) |
| | Pidsley reference (56178 probes) | 2074 (0.63%) | 2325 (0.64%) | 2319 (0.58%) | 2251 (0.54%) | 1376 (0.7%) |
| | McCartney reference (56448 probes) | 2507 (0.76%) | 2737 (0.76%) | 2873 (0.71%) | 2898 (0.69%) | 1704 (0.86%) |
| | Price reference (73907 probes) | 7483 (2.27%) | 8093 (2.24%) | 8498 (2.11%) | 8403 (2%) | 4713 (2.39%) |
| Total overlapped flagged and previously published probes | | 38861 (11.76%) | 23123 (6.39%) | 28884 (7.17%) | 22214 (5.29%) | 6069 (3.08%) |
| **Spearman correlation >0.7 or pooled SD < lower quartile pooled SD (high concordance probes)** | | | | | | |
| Cohort | | VHAS | CLHNS | CALERIE | BeCOME | Common across four cohorts |
| Probe count* | | 391044 (100%) | 359552 (100%) | 318709 (100%) | 301606 (100%) | 167533 (100%) |
| High-confidence mapping probes in cell line (539313 probes) | | 321350 (82.18%) | 295678 (82.24%) | 260058 (81.6%) | 248380 (82.35%) | 146326 (87.34%) |
| Flagged detectionP and beadcount probes** | | 45262 (11.57%) | 14890 (4.14%) | 19544 (6.13%) | 6741 (2.24%) | 342 (0.2%) |
| | EPICv2 design type switch | 52 (0.01%) | 45 (0.01%) | 39 (0.01%) | 39 (0.01%) | 21 (0.01%) |

| | | | | | | |
|---|---|---|---|---|---|---|
| | probes (82 probes) | | | | | |
| Previously published references | Peters reference (11878 probes) | 187 (0.05%) | 164 (0.05%) | 170 (0.05%) | 193 (0.06%) | 89 (0.05%) |
| | Pidsley reference (56178 probes) | 1473 (0.38%) | 1222 (0.34%) | 1228 (0.39%) | 1296 (0.43%) | 533 (0.32%) |
| | McCartney reference (56448 probes) | 2088 (0.53%) | 1858 (0.52%) | 1722 (0.54%) | 1697 (0.56%) | 852 (0.51%) |
| | Price reference (73907 probes) | 6480 (1.66%) | 5870 (1.63%) | 5465 (1.71%) | 5560 (1.84%) | 2721 (1.62%) |
| Total overlapped flagged and previously published probes | | 52095 (13.32%) | 21628 (6.02%) | 25611 (8.04%) | 13057 (4.33%) | 3592 (2.14%) |

* Percentage was calculated over the probe count by the Spearman correlation and pooled SD thresholds. **flagged detectionP and beadcount probes were calculated by cohort: VHAS: 101104; CLHNS: 40373; CALERIE: 53171; BeCOME: 30222; Common: 3734

**Supplementary Table 10. Paired t-tests comparing DNA methylation-based immune cell type proportions estimated using EPICv1 and EPICv2 with IDOL probes independently for each cohort. Spearman correlation (rho), paired t-test p-values and Bonferroni adjusted p-values, and Cohen's d effect size between EPICv1 and EPICv2 estimates obtained from matched samples in each cohort are provided.**

| Cell type | VHAS | | | | | | | | |
|---|---|---|---|---|---|---|---|---|---|
| | EPICv1 mean proportion (sd) | EPICv2 mean proportion (sd) | rho | p value | Bonferroni p | Mean of the differences | Cohen's d | Within EPICv1 technical replicate absolute differences | Within EPICv2 technical replicate absolute differences |
| Bas | 0.011(0.021) | 0.018(0.024) | 0.787 | <0.0001 | <0.0001 | 0.007 | 0.264 | 0.001 | 0.005 |
| Bmem | 0.021(0.008) | 0.026(0.007) | 0.942 | <0.0001 | <0.0001 | 0.004 | 0.469 | 0.004 | 0.006 |
| Bnv | 0.021(0.013) | 0.02(0.014) | 0.97 | 0.1757 | 1 | -0.001 | -0.064 | 0.002 | 0.009 |
| CD4mem | 0.067(0.025) | 0.072(0.027) | 0.921 | 0.019 | 0.2092 | 0.005 | 0.181 | 0.006 | 0.007 |
| CD4nv | 0.008(0.018) | 0.01(0.018) | 0.773 | 0.0212 | 0.2327 | 0.002 | 0.104 | 0 | 0 |
| CD8mem | 0.136(0.061) | 0.117(0.06) | 0.973 | <0.0001 | <0.0001 | -0.02 | -0.32 | 0.005 | 0.013 |
| CD8nv* | 0 | 0 | NA | NA | NA | 0 | NA | 0 | 0 |
| Eos | 0.039(0.054) | 0.029(0.052) | 0.94 | <0.0001 | <0.0001 | -0.01 | -0.172 | 0.001 | 0 |
| Mono | 0.068(0.022) | 0.073(0.022) | 0.936 | 1.00E-04 | 0.0016 | 0.005 | 0.218 | 0.003 | 0.001 |
| Neu | 0.519(0.08) | 0.531(0.084) | 0.997 | <0.0001 | <0.0001 | 0.012 | 0.124 | 0.016 | 0.006 |
| NK | 0.063(0.03) | 0.072(0.029) | 0.985 | <0.0001 | <0.0001 | 0.009 | 0.291 | 0.002 | 0.006 |
| Treg | 0.008(0.006) | 0.012(0.009) | 0.666 | 0.0053 | 0.0584 | 0.004 | 0.507 | 0.008 | 0.003 |
| Cell type | CLHNS** | | | | | | | | |
| | EPICv1 mean proportion (sd) | EPICv2 mean proportion (sd) | rho | p value | Bonferroni p | Mean of the differences | Cohen's d | Within EPICv1 technical replicate absolute | Within EPICv2 technical replicate absolute |

| | | | | | | | | | |
|---|---|---|---|---|---|---|---|---|---|
| | | | | | | | | differences | differences |
| Bas | 0.006(0.015) | 0.015(0.016) | 0.746 | <0.0001 | 5.00E-04 | 0.009 | 0.566 | NA | 0.004 |
| Bmem | 0.027(0.011) | 0.027(0.011) | 0.914 | 0.4811 | 1 | 0.001 | 0.056 | NA | 0.003 |
| Bnv | 0.022(0.011) | 0.025(0.011) | 0.914 | 0.0031 | 0.0336 | 0.003 | 0.259 | NA | 0.001 |
| CD4mem | 0.092(0.041) | 0.11(0.043) | 0.936 | <0.0001 | <0.0001 | 0.018 | 0.408 | NA | 0.009 |
| CD4nv | 0.013(0.017) | 0.015(0.017) | 0.968 | 0.2348 | 1 | 0.001 | 0.069 | NA | 0 |
| CD8mem | 0.138(0.039) | 0.118(0.04) | 0.975 | <0.0001 | <0.0001 | -0.019 | -0.482 | NA | 0.003 |
| CD8nv | 0 | 0 | NA | NA | NA | 0 | NA | NA | 0 |
| Eos | 0.03(0.016) | 0.018(0.018) | 0.928 | <0.0001 | <0.0001 | -0.012 | -0.702 | NA | 0 |
| Mono | 0.069(0.015) | 0.079(0.016) | 0.971 | <0.0001 | <0.0001 | 0.01 | 0.6 | NA | 0.002 |
| Neu | 0.483(0.083) | 0.5(0.088) | 0.996 | <0.0001 | <0.0001 | 0.018 | 0.17 | NA | 0.001 |
| NK | 0.048(0.022) | 0.058(0.023) | 0.968 | <0.0001 | <0.0001 | 0.01 | 0.422 | NA | 0.002 |
| Treg | 0.008(0.004) | 0.005(0.005) | 0.624 | 0.0103 | 0.1132 | -0.003 | -0.66 | NA | 0.005 |
| Cell type | **CALERIE** | | | | | | | | |
| | EPICv1 mean proportion (sd) | EPICv2 mean proportion (sd) | rho | p value | Bonferroni p | Mean of the differences | Cohen's d | Within EPICv1 technical replicate absolute differences | Within EPICv2 technical replicate absolute differences |
| Bas | 0.006(0.007) | 0.012(0.01) | 0.869 | <0.0001 | 5.00E-04 | 0.006 | 0.507 | 0.001 | 0.002 |
| Bmem | 0.017(0.006) | 0.019(0.007) | 0.901 | 0.0022 | 0.0261 | 0.002 | 0.296 | 0.002 | 0.001 |
| Bnv | 0.036(0.018) | 0.036(0.017) | 0.94 | 0.9469 | 1 | 0 | 0.003 | 0 | 0.002 |
| CD4mem | 0.101(0.026) | 0.108(0.024) | 0.945 | 8.00E-04 | 0.0091 | 0.007 | 0.258 | 0.007 | 0.011 |
| CD4nv | 0.067(0.033) | 0.07(0.033) | 0.954 | 0.0926 | 1 | 0.002 | 0.068 | 0.001 | 0 |

| | | | | | | | | | |
|---|---|---|---|---|---|---|---|---|---|
| CD8mem | 0.064(0.04) | 0.046(0.036) | 0.969 | <0.0001 | <0.0001 | -0.018 | -0.414 | 0.004 | 0.002 |
| CD8nv | 0.013(0.019) | 0.011(0.017) | 0.835 | 0.0121 | 0.1449 | -0.002 | -0.1 | 0 | 0 |
| Eos | 0.011(0.017) | 0.005(0.012) | 0.902 | 6.00E-04 | 0.0066 | -0.006 | -0.256 | 0.001 | 0 |
| Mono | 0.075(0.019) | 0.08(0.02) | 0.923 | <0.0001 | 2.00E-04 | 0.005 | 0.25 | 0.003 | 0.005 |
| Neu | 0.51(0.084) | 0.512(0.083) | 0.984 | 0.1326 | 1 | 0.002 | 0.026 | 0.004 | 0.004 |
| NK | 0.041(0.017) | 0.051(0.018) | 0.944 | <0.0001 | <0.0001 | 0.01 | 0.564 | 0.004 | 0.004 |
| Treg | 0.008(0.008) | 0.012(0.007) | 0.59 | 0.005 | 0.0595 | 0.004 | 0.488 | 0.003 | 0.009 |
| | **BeCOME*** (external validation)** | | | | | | | | |
| Cell type | EPICv1 mean proportion (sd) | EPICv2 mean proportion (sd) | rho | p value | Bonferroni p | Mean of the differences | Cohen's d | Within EPICv1 technical replicate absolute differences | Within EPICv2 technical replicate absolute differences |
| Bas | 0.012(0.005) | 0.013(0.005) | 0.476 | 0.4487 | 1 | 0.001 | 0.258 | NA | NA |
| Bmem | 0.018(0.007) | 0.023(0.007) | 0.667 | 0.0306 | 0.3677 | 0.005 | 0.676 | NA | NA |
| Bnv | 0.026(0.02) | 0.026(0.017) | 0.898 | 0.638 | 1 | -0.001 | -0.027 | NA | NA |
| CD4mem | 0.084(0.026) | 0.084(0.028) | 0.976 | 0.9812 | 1 | 0 | 0.002 | NA | NA |
| CD4nv | 0.085(0.063) | 0.084(0.059) | 0.952 | 0.7289 | 1 | -0.001 | -0.02 | NA | NA |
| CD8mem | 0.057(0.042) | 0.038(0.04) | 0.905 | 1.00E-04 | 9.00E-04 | -0.019 | -0.438 | NA | NA |
| CD8nv | 0.024(0.023) | 0.027(0.02) | 0.786 | 0.4187 | 1 | 0.002 | 0.107 | NA | NA |
| Eos | 0.011(0.008) | 0.004(0.006) | 0.774 | 0.0077 | 0.0927 | -0.006 | -0.82 | NA | NA |
| Mono | 0.072(0.02) | 0.074(0.019) | 0.952 | 0.6056 | 1 | 0.001 | 0.057 | NA | NA |
| Neu | 0.517(0.064) | 0.518(0.06) | 1 | 0.4308 | 1 | 0.002 | 0.021 | NA | NA |

| | | | | | | | | | |
|---|---|---|---|---|---|---|---|---|---|
| NK | 0.057(0.024) | 0.067(0.021) | 0.976 | 9.00E-04 | 0.0106 | 0.01 | 0.354 | NA | NA |
| Treg | 0.008(0.009) | 0.017(0.009) | 0.671 | 0.0045 | 0.0544 | 0.009 | 1.002 | NA | NA |

* CD8nv proportions were estimated to be 0 for all samples in VHAS and CLHNS for both EPIC versions. **CLHNS did not have technical replicate in EPICv1. ***BeCOME did not have technical replicate either EPIC versions.

**Supplementary Table 11. Paired t-tests comparing DNA methylation-based immune cell type proportions estimated using EPICv1 and EPICv2 with auto selected probes, independently for each cohort. Spearman correlation (rho), paired t-test p-values and Bonferroni adjusted p-values, and Cohen's d effect size between EPICv1 and EPICv2 estimates obtained from matched samples in each cohort are provided.**

| Cell type | VHAS | | | | | | | | |
|---|---|---|---|---|---|---|---|---|---|
| | EPICv1 mean proportion (sd) | EPICv2 mean proportion (sd) | rho | p value | Bonferroni p | Mean of the differences | Cohen's d | Within EPICv1 technical replicate absolute differences | Within EPICv2 technical replicate absolute differences |
| Bas | 0.023(0.019) | 0.029(0.02) | 0.897 | <0.0001 | <0.0001 | 0.006 | 0.308 | <0.001 | 0.001 |
| Bmem | 0.019(0.009) | 0.028(0.008) | 0.907 | <0.0001 | <0.0001 | 0.009 | 1.129 | 0.004 | 0.001 |
| Bnv | 0.024(0.016) | 0.025(0.015) | 0.971 | 0.0214 | 0.2571 | 0.002 | 0.109 | 0.004 | 0.003 |
| CD4mem | 0.113(0.019) | 0.07(0.022) | 0.743 | <0.0001 | <0.0001 | -0.043 | -2.103 | 0.008 | 0.009 |
| CD4nv | 0.049(0.023) | 0.015(0.013) | 0.733 | <0.0001 | <0.0001 | -0.034 | -1.429 | 0.002 | <0.001 |
| CD8mem | 0.086(0.04) | 0.099(0.046) | 0.946 | <0.0001 | <0.0001 | 0.014 | 0.243 | 0.001 | 0.004 |
| CD8nv* | 0 | 0.013(0.013) | NA | 1.00E-04 | 0.0014 | 0.013 | 1.423 | <0.001 | 0.003 |
| Eos | 0.03(0.046) | 0.032(0.046) | 0.923 | 0.1273 | 1 | 0.002 | 0.04 | 0.002 | <0.001 |
| Mono | 0.074(0.023) | 0.079(0.023) | 0.948 | 1.00E-04 | 7.00E-04 | 0.006 | 0.255 | 0.003 | <0.001 |
| Neu | 0.491(0.082) | 0.488(0.085) | 0.971 | 0.2984 | 1 | -0.002 | -0.026 | 0.01 | 0.002 |
| NK | 0.062(0.029) | 0.068(0.027) | 0.966 | <0.0001 | <0.0001 | 0.006 | 0.206 | 0.004 | <0.001 |
| Treg* | 0 | 0.033(0.008) | NA | <0.0001 | <0.0001 | 0.033 | 5.756 | 0 | 0.009 |
| Cell type | CLHNS** | | | | | | | | |
| | EPICv1 mean proportion (sd) | EPICv2 mean proportion (sd) | rho | p value | Bonferroni p | Mean of the differences | Cohen's d | Within EPICv1 technical replicate absolute | Within EPICv2 technical replicate absolute |

| | | | | | | | | differences | differences |
|---|---|---|---|---|---|---|---|---|---|
| Bas | 0.012(0.012) | 0.02(0.012) | 0.575 | <0.0001 | <0.0001 | 0.008 | 0.642 | NA | 0.002 |
| Bmem | 0.016(0.011) | 0.028(0.011) | 0.932 | <0.0001 | <0.0001 | 0.012 | 1.079 | NA | 0.005 |
| Bnv | 0.028(0.013) | 0.029(0.013) | 0.979 | 0.0014 | 0.017 | 0.002 | 0.127 | NA | 0.005 |
| CD4mem | 0.13(0.034) | 0.09(0.032) | 0.954 | <0.0001 | <0.0001 | -0.04 | -1.198 | NA | 0.005 |
| CD4nv | 0.054(0.022) | 0.012(0.013) | 0.88 | <0.0001 | <0.0001 | -0.042 | -1.862 | NA | <0.001 |
| CD8mem | 0.09(0.031) | 0.11(0.036) | 0.986 | <0.0001 | <0.0001 | 0.02 | 0.522 | NA | 0.006 |
| CD8nv* | 0 | 0.025(0.012) | NA | <0.0001 | <0.0001 | 0.025 | 2.847 | NA | 0.001 |
| Eos | 0.023(0.015) | 0.024(0.016) | 0.935 | 0.3864 | 1 | 0.001 | 0.082 | NA | 0.002 |
| Mono | 0.075(0.013) | 0.082(0.013) | 0.954 | <0.0001 | <0.0001 | 0.007 | 0.559 | NA | 0.002 |
| Neu | 0.455(0.085) | 0.468(0.094) | 0.982 | 2.00E-04 | 0.0027 | 0.013 | 0.075 | NA | <0.001 |
| NK | 0.052(0.022) | 0.056(0.021) | 0.946 | 0.0037 | 0.0441 | 0.004 | 0.192 | NA | 0.001 |
| Treg* | 0 | 0.027(0.006) | NA | <0.0001 | <0.0001 | 0.027 | 6.012 | NA | 0.004 |
| Cell type | **CALERIE** | | | | | | | | |
| | EPICv1 mean proportion (sd) | EPICv2 mean proportion (sd) | rho | p value | Bonferroni p | Mean of the differences | Cohen's d | Within EPICv1 technical replicate absolute differences | Within EPICv2 technical replicate absolute differences |
| Bas | 0.011(0.008) | 0.014(0.008) | 0.937 | <0.0001 | <0.0001 | 0.003 | 0.423 | <0.001 | 0.003 |
| Bmem | 0.01(0.007) | 0.016(0.007) | 0.934 | <0.0001 | <0.0001 | 0.006 | 0.785 | <0.001 | <0.001 |
| Bnv | 0.038(0.02) | 0.043(0.018) | 0.964 | <0.0001 | <0.0001 | 0.005 | 0.216 | 0.002 | 0.001 |
| CD4mem | 0.118(0.024) | 0.095(0.02) | 0.92 | <0.0001 | <0.0001 | -0.023 | -0.971 | 0.007 | 0.003 |

| | | | | | | | | | |
|---|---|---|---|---|---|---|---|---|---|
| CD4nv | 0.112(0.036) | 0.046(0.03) | 0.908 | <0.0001 | <0.0001 | -0.066 | -1.818 | 0.004 | 0.007 |
| CD8mem | 0.049(0.029) | 0.056(0.03) | 0.958 | 5.00E-04 | 0.006 | 0.007 | 0.222 | 0.004 | <0.001 |
| CD8nv* | 0.001(0.004) | 0.05(0.019) | 0.573 | <0.0001 | <0.0001 | 0.049 | 2.238 | <0.001 | 0.007 |
| Eos | 0.01(0.013) | 0.009(0.013) | 0.87 | 0.3018 | 1 | -0.001 | -0.06 | <0.001 | <0.001 |
| Mono | 0.082(0.019) | 0.086(0.018) | 0.92 | 0.0015 | 0.0179 | 0.004 | 0.211 | <0.001 | 0.002 |
| Neu | 0.482(0.086) | 0.478(0.086) | 0.995 | 8.00E-04 | 0.0091 | -0.004 | -0.05 | 0.006 | <0.001 |
| NK | 0.041(0.014) | 0.046(0.015) | 0.957 | <0.0001 | <0.0001 | 0.005 | 0.346 | 0.001 | <0.001 |
| Treg* | 0(0) | 0.022(0.01) | NA | <0.0001 | <0.0001 | 0.022 | 3.022 | 0 | <0.001 |
| Cell type | **BeCOME*** (external validation)** | | | | | | | | |
| | EPICv1 mean proportion (sd) | EPICv2 mean proportion (sd) | rho | p value | Bonferroni p | Mean of the differences | Cohen's d | Within EPICv1 technical replicate absolute differences | Within EPICv2 technical replicate absolute differences |
| Bas | 0.024(0.004) | 0.028(0.005) | 0.683 | 0.0104 | 0.1247 | 0.004 | 0.912 | NA | NA |
| Bmem | 0.02(0.006) | 0.026(0.007) | 0.857 | 0.0012 | 0.0138 | 0.006 | 0.779 | NA | NA |
| Bnv | 0.03(0.023) | 0.033(0.022) | 0.952 | 0.0099 | 0.1185 | 0.003 | 0.132 | NA | NA |
| CD4mem | 0.11(0.021) | 0.066(0.022) | 0.81 | <0.0001 | 1.00E-04 | -0.044 | -2.018 | NA | NA |
| CD4nv | 0.128(0.054) | 0.057(0.045) | 0.976 | <0.0001 | <0.0001 | -0.07 | -0.993 | NA | NA |
| CD8mem | 0.051(0.027) | 0.052(0.027) | 0.976 | 0.4374 | 1 | 0.002 | 0.06 | NA | NA |
| CD8nv | 0.006(0.012) | 0.064(0.018) | 0.733 | <0.0001 | <0.0001 | 0.057 | 2.933 | NA | NA |
| Eos | 0.032(0.011) | 0.015(0.012) | 0.881 | <0.0001 | 4.00E-04 | -0.017 | -1.37 | NA | NA |
| Mono | 0.075(0.017) | 0.076(0.014) | 0.952 | 0.5139 | 1 | 0.001 | 0.07 | NA | NA |

| | | | | | | | | | |
|---|---|---|---|---|---|---|---|---|---|
| Neu | 0.449(0.061) | 0.442(0.051) | 0.976 | 0.1557 | 1 | -0.007 | -0.078 | NA | NA |
| NK | 0.05(0.017) | 0.053(0.018) | 0.881 | 0.1129 | 1 | 0.003 | 0.167 | NA | NA |
| Treg | 0.001(0.002) | 0.054(0.011) | 0.514 | <0.0001 | <0.0001 | 0.053 | 4.985 | NA | NA |

* CD8nv proportions were estimated to be 0 for all samples in VHAS and CLHNS, and 21 out of 24 samples of CALERIE for EPICv1; Treg proportions were estimated to be 0 for all VHAS, CLHNS and CALERIE for EPICv1. The Spearman correlations were therefore not calculated. **CLHNS did not have technical replicate in EPICv1. ***BeCOME did not have technical replicate either EPIC versions.

**Supplementary Table 12. Pearson correlations of epigenetic clock estimates with chronological age in EPICv1 and EPICv2 samples of VHAS, CLHNS, CALERIE and BeCOME.**

| Cohort | Clock | EPICv1 | | | EPICv2 | | |
|---|---|---|---|---|---|---|---|
| | | r | MAE | MaxAE | r | MAE | MaxAE |
| VHAS | Horvath_pan-tissue | 0.725 | 4.57 | 11.6 | 0.765 | 5.04 | 11.73 |
| | Hannum | 0.743 | 12.57 | 21.14 | 0.656 | 34.42 | 43.22 |
| | Horvath_SkinBlood | 0.817 | 5.64 | 11.92 | 0.812 | 14.04 | 20.15 |
| | PhenoAge | 0.677 | 11.83 | 23.03 | 0.583 | 7.67 | 20.46 |
| | GrimAge | 0.794 | 4.15 | 12.51 | 0.769 | 6.54 | 12.2 |
| CLHNS | Horvath_pan-tissue | 0.883 | 7.23 | 13.92 | 0.807 | 9.08 | 12.89 |
| | Hannum | 0.912 | 6.15 | 14.18 | 0.87 | 25.96 | 32.35 |
| | Horvath_SkinBlood | 0.905 | 3.08 | 6.52 | 0.918 | 8 | 14.22 |
| | PhenoAge | 0.853 | 4.54 | 8.99 | 0.891 | 2.26 | 7.48 |
| | GrimAge | 0.819 | 6.57 | 15.34 | 0.845 | 2.71 | 7.97 |
| CALERIE | Horvath_pan-tissue | 0.862 | 5.63 | 13.54 | 0.888 | 4.66 | 13.02 |
| | Hannum | 0.872 | 7.45 | 13.06 | 0.898 | 29.17 | 36.31 |
| | Horvath_SkinBlood | 0.947 | 3.06 | 7.03 | 0.956 | 11.28 | 16.14 |
| | PhenoAge | 0.857 | 11.85 | 20.02 | 0.822 | 8.02 | 20.51 |
| | GrimAge | 0.954 | 14.42 | 20.53 | 0.96 | 14.75 | 20.7 |
| BeCOME (external validation) | Horvath_pan-tissue | 0.984 | 4.97 | 8.89 | 0.98 | 4.74 | 8.69 |
| | Hannum | 0.955 | 9.91 | 19.89 | 0.957 | 29.09 | 39.65 |
| | Horvath_SkinBlood | 0.995 | 4.52 | 7.16 | 0.996 | 10.32 | 13.73 |
| | PhenoAge | 0.967 | 11.86 | 20.21 | 0.976 | 6.58 | 12.79 |
| | GrimAge | 0.995 | 13.33 | 21.87 | 0.992 | 4.26 | 9.74 |

r: Pearson correlation between epigenetic age and chronological age. MAE: mean absolute error in years. MaxAE: maximum absolute error in years

**Supplementary Table 13. Comparison of epigenetic ages between EPICv1 and EPICv2 in first-, second-, rate-based and other epigenetic clocks.**

| Cohort | Clock | r | p-value | Bonferroni p | Mean of the differences | Cohen's d | Within EPICv1 technical replicate absolute differences | Within EPICv2 technical replicate absolute differences | Mean absolute CpG beta value differences (sd) | Mean CpG beta value pooled sd (sd) |
|---|---|---|---|---|---|---|---|---|---|---|
| VHAS | Horvath_pan-tissue | 0.952 | 0.4018 | 1 | 0.586 | 0.057 | 1.936 | 2.654 | 0.0157(0.0161) | 0.0375(0.0222) |
| | Hannum | 0.934 | <0.0001 | <0.0001 | -21.851 | -3.342 | 1.713 | 1.906 | 0.0222(0.0475) | 0.045(0.0187) |
| | Horvath_SkinBlood | 0.963 | <0.0001 | <0.0001 | -8.874 | -1.373 | 2.497 | 0.546 | 0.02(0.0315) | 0.0449(0.0223) |
| | PhenoAge | 0.925 | <0.0001 | <0.0001 | 6.051 | 0.604 | 4.206 | 2.994 | 0.016(0.0186) | 0.039(0.0248) |
| | GrimAge** | 0.978 | <0.0001 | <0.0001 | -5.608 | -0.648 | 0.523 | 0.955 | NA | NA |
| | DunedinPACE | 0.94 | <0.0001 | <0.0001 | 0.058 | 0.458 | 0.028 | 0.074 | 0.023(0.0258) | 0.0531(0.0274) |
| | DNAmTL | 0.955 | <0.0001 | <0.0001 | 0.21 | 0.968 | 0.052 | 0.057 | 0.0185(0.0182) | 0.0574(0.0229) |
| | epiTOC | 0.983 | 0.0938 | 0.7501 | -0.002 | -0.069 | 0.003 | 0.003 | 0.0162(0.0184) | 0.032(0.0177) |
| CLHNS* | Horvath_pan-tissue | 0.879 | 0.0141 | 0.113 | 1.791 | 0.356 | NA | 3.408 | 0.0179(0.0183) | 0.0276(0.0169) |
| | Hannum | 0.913 | <0.0001 | <0.0001 | -19.811 | -4.137 | NA | 1.303 | 0.0279(0.0378) | 0.0314(0.0102) |
| | Horvath_SkinBlood | 0.935 | <0.0001 | <0.0001 | -10.454 | -1.723 | NA | 0.151 | 0.023(0.0281) | 0.0324(0.0177) |
| | PhenoAge | 0.897 | <0.0001 | <0.0001 | 5.583 | 0.928 | NA | 2.683 | 0.018(0.0201) | 0.0281(0.0188) |
| | GrimAge** | 0.992 | <0.0001 | <0.0001 | -7.182 | -1.034 | NA | 1.306 | NA | NA |
| | DunedinPACE | 0.912 | 0.044 | 0.352 | 0.021 | 0.24 | NA | 0.01 | 0.0256(0.0258) | 0.0367(0.0198) |
| | DNAmTL | 0.838 | <0.0001 | <0.0001 | 0.187 | 1.496 | NA | 0.045 | 0.0245(0.0197) | 0.0415(0.0193) |
| | epiTOC | 0.967 | <0.0001 | 1.00E-04 | -0.007 | -0.426 | NA | 0.004 | 0.0191(0.0201) | 0.0231(0.0126) |

| | | | | | | | | | | |
|---|---|---|---|---|---|---|---|---|---|---|
| CALERIE | Horvath_pan-tissue | 0.918 | 0.097 | 0.7761 | -1.419 | -0.15 | 0.13 | 2.477 | 0.0182(0.015) | 0.0299(0.0164) |
| | Hannum | 0.971 | <0.0001 | <0.0001 | -21.723 | -2.74 | 1.738 | 1.441 | 0.0282(0.0388) | 0.0377(0.0173) |
| | Horvath_SkinBlood | 0.988 | <0.0001 | <0.0001 | -9.062 | -0.836 | 0.72 | 2.095 | 0.023(0.0274) | 0.0348(0.0177) |
| | PhenoAge | 0.942 | <0.0001 | 1.00E-04 | 4.33 | 0.425 | 2.25 | 4.142 | 0.0203(0.0175) | 0.0302(0.0176) |
| | GrimAge** | 0.996 | 0.014 | 0.112 | 0.33 | 0.048 | 0.658 | 0.245 | NA | NA |
| | DunedinPACE | 0.912 | <0.0001 | <0.0001 | 0.069 | 0.576 | 0.024 | 0.039 | 0.0238(0.0223) | 0.0387(0.0191) |
| | DNAmTL | 0.94 | <0.0001 | <0.0001 | 0.156 | 0.632 | 0.02 | 0.028 | 0.0232(0.0174) | 0.0463(0.0176) |
| | epiTOC | 0.813 | 4.00E-04 | 0.0036 | 0.004 | 0.542 | 0.004 | 0.001 | 0.0131(0.0121) | 0.0158(0.0104) |
| BeCOME*** | Horvath_pan-tissue | 0.965 | 0.9172 | 1 | 0.167 | 0.01 | NA | NA | 0.0229(0.0221) | 0.0329(0.0228) |
| | Hannum | 0.995 | <0.0001 | <0.0001 | -19.175 | -1.247 | NA | NA | 0.0353(0.0412) | 0.0496(0.031) |
| | Horvath_SkinBlood | 0.996 | 1.00E-04 | 7.00E-04 | -5.806 | -0.267 | NA | NA | 0.0268(0.0285) | 0.0409(0.0284) |
| | PhenoAge | 0.99 | 9.00E-04 | 6.90E-03 | 5.495 | 0.275 | NA | NA | 0.0217(0.0234) | 0.0331(0.025) |
| | GrimAge** | 0.988 | <0.0001 | <0.0001 | 16.51 | 1.007 | NA | NA | NA | NA |
| | DunedinPACE | 0.869 | 9.00E-04 | 0.007 | 0.087 | 1.003 | NA | NA | 0.022(0.0263) | 0.042(0.0268) |
| | DNAmTL | 0.975 | 9.00E-04 | 0.0075 | 0.144 | 0.432 | NA | NA | 0.0268(0.0246) | 0.0566(0.0291) |
| | epiTOC | 0.699 | 2.00E-04 | 0.0016 | 0.008 | 1.931 | NA | NA | 0.0123(0.0155) | 0.0117(0.0101) |

r: Pearson correlation between EPICv1 epigenetic age and EPICv2 epigenetic age. p value: p value from paired T test between EPICv1 epigenetic age and EPICv2 epigenetic age. sd: standard deviation. *CLHNS did not have technical replicate in EPICv1; **GrimAge does not have a publically available list of clock CpGs; ***BeCOME did not have technical replicate either EPIC versions.

**Supplementary Table 14. Pearson correlation (r) and paired T tests results of EAAs of between EPICv1 and EPICv2 in Horvath pantissue, Hannum, Horvath SkinBlood, PhenoAge, and GrimAge.**

| VHAS | | | | | |
|---|---|---|---|---|---|
| EAA Category | Clock | r | p value | Bonferroni adjusted p | Cohen's d |
| EPIC version separate | Horvath_pan-tissue | 0.895 | 1 | 1 | 0 |
| | Hannum | 0.885 | 1 | 1 | 0 |
| | Horvath_SkinBlood | 0.89 | 1 | 1 | 0 |
| | PhenoAge | 0.887 | 1 | 1 | 0 |
| | GrimAge | 0.946 | 1 | 1 | 0 |
| EPIC versions combined | Horvath_pan-tissue | 0.851 | 0.4018 | 1 | 0.099 |
| | Hannum | 0.867 | <0.0001 | <0.0001 | -4.75 |
| | Horvath_SkinBlood | 0.889 | <0.0001 | <0.0001 | -2.371 |
| | PhenoAge | 0.884 | <0.0001 | <0.0001 | 0.751 |
| | GrimAge | 0.946 | <0.0001 | <0.0001 | -1.021 |
| EPIC versions combined and adjusted | Horvath_pan-tissue | 0.851 | 1 | 1 | 0 |
| | Hannum | 0.867 | 1 | 1 | 0 |
| | Horvath_SkinBlood | 0.889 | 1 | 1 | 0 |
| | PhenoAge | 0.884 | 1 | 1 | 0 |
| | GrimAge | 0.946 | 1 | 1 | 0 |
| **CLHNS** | | | | | |
| EAA Category | Clock | r | p value | Bonferroni adjusted p | Cohen's d |
| EPIC version separate | Horvath_pan-tissue | 0.6 | 1 | 1 | 0 |
| | Hannum | 0.592 | 1 | 1 | 0 |
| | Horvath_SkinBlood | 0.616 | 1 | 1 | 0 |
| | PhenoAge | 0.577 | 1 | 1 | 0 |
| | GrimAge | 0.976 | 1 | 1 | 0 |
| EPIC versions combined | Horvath_pan-tissue | 0.597 | 0.0141 | 0.1412 | 0.65 |
| | Hannum | 0.59 | <0.0001 | <0.0001 | -8.98 |
| | Horvath_SkinBlood | 0.609 | <0.0001 | <0.0001 | -4.214 |
| | PhenoAge | 0.574 | <0.0001 | <0.0001 | 1.887 |
| | GrimAge | 0.958 | <0.0001 | <0.0001 | -2.311 |
| EPIC versions combined and adjusted | Horvath_pan-tissue | 0.597 | 0.9329 | 1 | 0.02 |
| | Hannum | 0.59 | 0.896 | 1 | -0.031 |

| | | | | | |
|---|---|---|---|---|---|
| | Horvath_SkinBlood | 0.609 | 0.7817 | 1 | 0.064 |
| | PhenoAge | 0.575 | 0.8568 | 1 | 0.044 |
| | GrimAge | 0.958 | 0.4918 | 1 | 0.053 |
| **CALERIE** | | | | | |
| EAA Category | Clock | r | p value | Bonferroni adjusted p | Cohen's d |
| EPIC version separate | Horvath_pan-tissue | 0.653 | 1 | 1 | 0 |
| | Hannum | 0.872 | 1 | 1 | 0 |
| | Horvath_SkinBlood | 0.873 | 1 | 1 | 0 |
| | PhenoAge | 0.808 | 1 | 1 | 0 |
| | GrimAge | 0.96 | 1 | 1 | 0 |
| EPIC versions combined | Horvath_pan-tissue | 0.619 | 0.097 | 0.9701 | -0.323 |
| | Hannum | 0.863 | <0.0001 | <0.0001 | -5.936 |
| | Horvath_SkinBlood | 0.83 | <0.0001 | <0.0001 | -3.1 |
| | PhenoAge | 0.796 | <0.0001 | 1.00E-04 | 0.795 |
| | GrimAge | 0.96 | 0.014 | 0.14 | 0.161 |
| EPIC versions combined and adjusted | Horvath_pan-tissue | 0.619 | 1 | 1 | 0 |
| | Hannum | 0.863 | 1 | 1 | 0 |
| | Horvath_SkinBlood | 0.83 | 1 | 1 | 0 |
| | PhenoAge | 0.796 | 1 | 1 | 0 |
| | GrimAge | 0.96 | 1 | 1 | 0 |
| **BeCOME** | | | | | |
| EAA Category | Clock | r | p value | Bonferroni adjusted p | Cohen's d |
| EPIC version separate | Horvath_pan-tissue | -0.014 | 1 | 1 | 0 |
| | Hannum | 0.947 | 1 | 1 | 0 |
| | Horvath_SkinBlood | 0.485 | 1 | 1 | 0 |
| | PhenoAge | 0.837 | 1 | 1 | 0 |
| | GrimAge | 0.087 | 1 | 1 | 0 |
| EPIC versions combined | Horvath_pan-tissue | -0.051 | 0.9172 | 1 | 0.055 |
| | Hannum | 0.944 | <0.0001 | <0.0001 | -4.333 |
| | Horvath_SkinBlood | 0.25 | 1.00E-04 | 9.00E-04 | -3.477 |
| | PhenoAge | 0.812 | 9.00E-04 | 0.0087 | 1.203 |

| | | | | | |
|---|---|---|---|---|---|
| | GrimAge | -0.199 | <0.0001 | <0.0001 | 10.056 |
| EPIC versions combined and adjusted | Horvath_pan-tissue | -0.051 | 1 | 1 | 0 |
| | Hannum | 0.944 | 1 | 1 | 0 |
| | Horvath_SkinBlood | 0.25 | 1 | 1 | 0 |
| | PhenoAge | 0.812 | 1 | 1 | 0 |
| | GrimAge | -0.199 | 1 | 1 | 0 |

r: Pearson correlation EPICv1 EAA (epigenetic age acceleration) and EPICv2 EAA. p value: p value from paired T test between EPICv1EAA and EPICv2 EAA.

**Supplementary Table 15. Pearson correlation (r) and paired T tests results of EPICv1 and EPICv2 comparisons for DunedinPACE, DNAmTL and epiTOC.**

| **VHAS** | | | | | | | | |
|---|---|---|---|---|---|---|---|---|
| Category | Clock | r | p value | Bonferroni p | Mean of the differences (rate/clock unit) | Cohen's d | Within EPICv1 technical replicate absolute differences | Within EPICv2 technical replicate absolute differences |
| EPIC version separate | DunedinPACE | 0.94 | <0.0001 | <0.0001 | 0.058 | 0.458 | 0.028 | 0.074 |
| | DNAmTL | 0.955 | <0.0001 | <0.0001 | 0.21 | 0.968 | 0.052 | 0.057 |
| | epiTOC | 0.983 | 0.0938 | 0.2813 | -0.002 | -0.069 | 0.003 | 0.003 |
| EPIC versions combined and adjusted | DunedinPACE | 0.94 | 0.9399 | 1 | -0.001 | -0.006 | 0.028 | 0.074 |
| | DNAmTL | 0.955 | 0.9763 | 1 | 0 | 0.002 | 0.052 | 0.057 |
| | epiTOC | 0.983 | 0.9425 | 1 | 0 | 0.003 | 0.003 | 0.003 |
| **CLHNS*** | | | | | | | | |
| Category | Clock | r | p value | Bonferroni p | Mean of the differences (rate/clock unit) | Cohen's d | Within EPICv1 technical replicate absolute differences | Within EPICv2 technical replicate absolute differences |
| EPIC version separate | DunedinPACE | 0.912 | 0.044 | 0.132 | 0.021 | 0.24 | NA | 0.01 |
| | DNAmTL | 0.838 | <0.0001 | <0.0001 | 0.187 | 1.496 | NA | 0.045 |
| | epiTOC | 0.967 | <0.0001 | <0.0001 | -0.007 | -0.426 | NA | 0.004 |
| EPIC versions combined and adjusted | DunedinPACE | 0.912 | 0.6902 | 1 | 0.004 | 0.044 | NA | 0.01 |
| | DNAmTL | 0.838 | 0.7781 | 1 | 0.005 | 0.042 | NA | 0.045 |
| | epiTOC | 0.967 | 0.2163 | 0.649 | -0.001 | -0.086 | NA | 0.004 |
| **CALERIE** | | | | | | | | |
| Category | Clock | r | p value | Bonferroni p | Mean of the differences (rate/clock unit) | Cohen's d | Within EPICv1 technical replicate absolute differences | Within EPICv2 technical replicate absolute differences |
| | DunedinPACE | 0.912 | <0.0001 | <0.0001 | 0.069 | 0.576 | 0.024 | 0.039 |

| | | | | | | | | |
|---|---|---|---|---|---|---|---|---|
| EPIC version separate | DNAmTL | 0.94 | <0.0001 | <0.0001 | 0.156 | 0.632 | 0.02 | 0.028 |
| | epiTOC | 0.813 | 4.00E-04 | 0.0013 | 0.004 | 0.542 | 0.004 | 0.001 |
| EPIC versions combined and adjusted | DunedinPACE | 0.912 | 0.7477 | 1 | -0.003 | -0.029 | 0.024 | 0.039 |
| | DNAmTL | 0.94 | 0.9929 | 1 | 0 | 0.001 | 0.02 | 0.028 |
| | epiTOC | 0.813 | 0.8109 | 1 | 0 | -0.032 | 0.004 | 0.001 |
| **BeCOME**** | | | | | | | | |
| Category | Clock | r | p value | Bonferroni p | Mean of the differences (rate(rate/clock unit)) | Cohen's d | Within EPICv1 technical replicate absolute differences | Within EPICv2 technical replicate absolute differences |
| EPIC version separate | DunedinPACE | 0.869 | 9.00E-04 | 0.0026 | 0.087 | 1.003 | NA | NA |
| | DNAmTL | 0.975 | 9.00E-04 | 0.0028 | 0.144 | 0.432 | NA | NA |
| | epiTOC | 0.699 | 2.00E-04 | 6.00E-04 | 0.008 | 1.931 | NA | NA |
| EPIC versions combined and adjusted | DunedinPACE | 0.869 | 1 | 1 | 0 | 0 | NA | NA |
| | DNAmTL | 0.975 | 1 | 1 | 0 | 0 | NA | NA |
| | epiTOC | 0.699 | 1 | 1 | 0 | 0 | NA | NA |

r: Pearson correlation EPICv1 estimates and EPICv2 estimates. p value: p value from paired T test between EPICv1 estimates and EPICv2 estimates; *CLHNS did not have technical replicate in EPICv1. **BeCOME did not have technical replicate either EPIC versions.

**Supplementary Table 16. Comparison of DNA methylation-based predictor estimations between EPv1 and EPICv2.**

| VHAS | | | | | | | | | | |
|---|---|---|---|---|---|---|---|---|---|---|
| Category | Predictor | r | *p-value* | Bonferroni p | Mean of the differences | Cohen's d | Within EPICv1 technical replicate absolute differences | Within EPICv2 technical replicate absolute differences | Mean absolute CpG beta value differences (sd) | Mean CpG beta value pooled sd (sd) |
| EPIC version separate | IL-6 Score | 0.894 | 0.0929 | 0.3716 | 0.012 | 0.173 | 0.029 | 0.018 | 0.0203(0.0409) | 0.0658(0.0318) |
| | CRP Score | 0.993 | <0.0001 | <0.0001 | 9.423 | 0.258 | 4.373 | 3.666 | 0.022(0.024) | 0.049(0.0203) |
| | Smoking Score | 0.982 | 0.1862 | 0.7448 | -0.048 | -0.055 | 0.155 | 0.19 | 0.0271(0.0339) | 0.0533(0.0422) |
| | Alcohol Score | 0.788 | 0.0552 | 0.2206 | 0.096 | 0.282 | 0.128 | 0.075 | 0.0258(0.0386) | 0.0498(0.0366) |
| EPIC versions combined and adjusted | IL-6 Score | 0.894 | 0.9285 | 1 | -0.001 | -0.009 | 0.029 | 0.018 | NA | NA |
| | CRP Score | 0.993 | 0.8989 | 1 | 0.118 | 0.003 | 4.373 | 3.666 | NA | NA |
| | Smoking Score | 0.982 | 0.7222 | 1 | 0.013 | 0.014 | 0.155 | 0.19 | NA | NA |
| | Alcohol Score | 0.788 | 0.973 | 1 | -0.002 | -0.005 | 0.128 | 0.075 | NA | NA |
| **CLHNS*** | | | | | | | | | | |
| Category | Predictor | r | *p-value* | Bonferroni p | Mean of the differences | Cohen's d | Within EPICv1 technical replicate absolute differences | Within EPICv2 technical replicate absolute differences | Mean absolute CpG beta value differences (sd) | Mean CpG beta value pooled sd (sd) |
| EPIC version separate | IL-6 Score | 0.912 | 0.0239 | 0.0958 | 0.02 | 0.275 | NA | 0.001 | 0.032(0.0416) | 0.0445(0.0237) |
| | CRP Score | 0.992 | <0.0001 | <0.0001 | 8.734 | 0.271 | NA | 1.255 | 0.0261(0.0269) | 0.0372(0.0165) |
| | Smoking Score | 0.99 | <0.0001 | <0.0001 | -0.274 | -0.301 | NA | 0.028 | 0.0342(0.0407) | 0.0371(0.0289) |

| | | | | | | | | | | |
|---|---|---|---|---|---|---|---|---|---|---|
| | Alcohol Score | 0.76 | 0.2589 | 1 | -0.055 | -0.211 | NA | 0.148 | 0.0285(0.04) | 0.0361(0.0283) |
| EPIC versions combined and adjusted | IL-6 Score | 0.912 | 0.2462 | 0.9847 | 0.009 | 0.131 | NA | 0.001 | NA | NA |
| | CRP Score | 0.992 | 0.0405 | 0.1619 | -2.43 | -0.075 | NA | 1.255 | NA | NA |
| | Smoking Score | 0.99 | 0.6759 | 1 | 0.014 | 0.016 | NA | 0.028 | NA | NA |
| | Alcohol Score | 0.76 | 0.6096 | 1 | -0.025 | -0.094 | NA | 0.148 | NA | NA |
| **CALERIE** | | | | | | | | | | |
| Category | Predictor | r | *p-value* | Bonferroni p | Mean of the differences | Cohen's d | Within EPICv1 technical replicate absolute differences | Within EPICv2 technical replicate absolute differences | Mean absolute CpG beta value differences (sd) | Mean CpG beta value pooled sd (sd) |
| EPIC version separate | IL-6 Score | 0.89 | 7.00E-04 | 0.0027 | 0.03 | 0.399 | 0.027 | 0.027 | 0.0213(0.0359) | 0.0416(0.0193) |
| | CRP Score | 0.945 | 0.3791 | 1 | -1.992 | -0.063 | 12.097 | 4.614 | 0.0245(0.0244) | 0.0381(0.0144) |
| | Smoking Score | 0.736 | 0.2754 | 1 | -0.049 | -0.173 | 0.064 | 0.2 | 0.0294(0.0352) | 0.0377(0.0311) |
| | Alcohol Score | 0.769 | <0.0001 | <0.0001 | 0.326 | 1.224 | 0.2 | 0.137 | 0.0255(0.0393) | 0.0393(0.031) |
| EPIC versions combined and adjusted | IL-6 Score | 0.89 | 0.4664 | 1 | -0.006 | -0.074 | 0.027 | 0.027 | NA | NA |
| | CRP Score | 0.945 | 0.8003 | 1 | 0.568 | 0.018 | 12.097 | 4.614 | NA | NA |
| | Smoking Score | 0.736 | 0.6544 | 1 | -0.02 | -0.07 | 0.064 | 0.2 | NA | NA |
| | Alcohol Score | 0.769 | 0.9802 | 1 | -0.001 | -0.004 | 0.2 | 0.137 | NA | NA |
| **BeCOME**** | | | | | | | | | | |
| Category | Predictor | r | *p-value* | Bonferroni p | Mean of the differences | Cohen's d | Within EPICv1 technical replicate absolute differences | Within EPICv2 technical replicate absolute | Mean absolute CpG beta value differences (sd) | Mean CpG beta value pooled sd (sd) |

| | | | | | | | | differences | | |
|---|---|---|---|---|---|---|---|---|---|---|
| EPIC version separate | IL-6 Score | 0.971 | 0.0042 | 0.0169 | 0.051 | 0.353 | NA | NA | 0.0242(0.0337) | 0.0455(0.0262) |
| | CRP Score | 0.964 | 0.8343 | 1 | -0.717 | -0.021 | NA | NA | 0.0249(0.0279) | 0.0408(0.0173) |
| | Smoking Score | 0.897 | 0.9412 | 1 | 0.004 | 0.012 | NA | NA | 0.0281(0.0369) | 0.0365(0.0355) |
| | Alcohol Score | 0.873 | 0.0249 | 0.0994 | 0.163 | 0.506 | NA | NA | 0.0282(0.046) | 0.0397(0.0351) |
| EPIC versions combined and adjusted | IL-6 Score | 0.971 | 1 | 1 | 0 | 0 | NA | NA | NA | NA |
| | CRP Score | 0.964 | 1 | 1 | 0 | 0 | NA | NA | NA | NA |
| | Smoking Score | 0.897 | 1 | 1 | 0 | 0 | NA | NA | NA | NA |
| | Alcohol Score | 0.873 | 1 | 1 | 0 | 0 | NA | NA | NA | NA |

r: Pearson correlation between EPICv1 predictor estimates and EPICv2 predictor estimates . p -value: p value from paired T test between EPICv1predictor estimates and EPICv2 predictor estimates. sd: standard deviation. *CLHNS did not have technical replicate in EPICv1.**BeCOME did not have technical replicate either EPIC versions.

**Supplementary Table 17. Pearson correlations of Garma and Quintela-Fandino's epigenetic clock estimates with chronological age in EPICv1 and EPICv2 samples of VHAS, CLHNS, CALERIE and BeCOME.**

| | EPICv1 and Age | | | EPICv2 and Age | | | EPICv1 and EPICv2 | | |
|---|---|---|---|---|---|---|---|---|---|
| Cohort | r | MAE | MaxAE | r | MAE | MaxAE | r | MAE | MaxAE |
| VHAS | 0.802 | 3.63 | 8.91 | 0.843 | 3 | 7.28 | 0.974 | 1.5 | 3.41 |
| CLHNS | 0.748 | 2.81 | 11.77 | 0.788 | 3.42 | 13.18 | 0.986 | 1.63 | 3.73 |
| CALERIE | 0.084 | 8.39 | 28.51 | 0.007 | 8.27 | 26.09 | -0.056 | 8.68 | 19.31 |

**Supplementary Table 18. Illumina's 17 quality control metrics, detection p-value, beadcount, average methylated and unmethylated intensity values estimated based on EPICv1 and EPICv2 DNA methylation data. For all three cohorts, values for each metric are provided as mean (standard deviation) or percentages separated by array version.**

| Quality control metrics | Recommended cutoff | VHAS (n=48) | | CLHNS (n=15) | | CALERIE (n=24) | | BeCOME (n=8) | |
|---|---|---|---|---|---|---|---|---|---|
| | | EPICv1 mean (sd) | EPICv2 mean (sd) | EPICv1 mean (sd) | EPICv2 mean (sd) | EPICv1 mean (sd) | EPICv2 mean (sd) | EPICv1 mean (sd) | EPICv2 mean (sd) |
| Restoration | >0 | 0.048(0.014) | 0.083(0.019) | 0.085(0.029) | 0.068(0.021) | 0.063(0.013) | 0.12(0.021) | 0.04(0.007) | 0.091(0.016) |
| Staining Green | >5 | 172.255(117.868) | 116.111 (16.278) | 159.744 (123.311) | 124.404 (18.032) | 163.991(68.689) | 78.66(17.939) | 173.641 (53.698) | 108.362 (9.153) |
| Staining Red | >5 | 38.79(12.157) | 62.18(7.861) | 51.26(15.632) | 89.799(11.848) | 51.423(7.972) | 62.969(11.31) | 78.829(21.417) | 72.545(3.749) |
| Extension Green | >5 | 46.501(3.916) | 28.602(3.204) | 33.173(5.29) | 34.466(3.886) | 45.396(4.612) | 26.34(3.574) | 77.723(21.253) | 38.598(3.561) |
| Extension Red | >5 | 16.225(1.122) | 16.1(0.821) | 21.388(1.902) | 26.002(1.611) | 19.855(1.081) | 22.494(1.313) | 16.932(3.504) | 16.579(3.901) |
| Hybridization High/Medium | >1 | 1.563(0.029) | 1.621(0.041) | 1.565(0.031) | 1.726(0.045) | 1.696(0.029) | 1.621(0.033) | 1.645(0.058) | 1.638(0.103) |
| Hybridization Medium/Low | >1 | 1.839(0.039) | 1.831(0.077) | 1.856(0.037) | 1.983(0.067) | 1.894(0.062) | 1.904(0.07) | 1.896(0.047) | 1.867(0.068) |
| Target Removal 1 | >1 | 18.251(3.384) | 9.356(1.487) | 12.283(3.954) | 8.944(1.567) | 12.735(1.521) | 5.726(0.856) | 20.554(4.221) | 10.183(1.517) |
| Target Removal 2 | >1 | 14.577(2.334) | 6.522(1.023) | 9.698(2.674) | 6.635(0.987) | 10.401(1.334) | 4.629(0.696) | 17.484(2.097) | 7.752(1.015) |
| Bisulfite Conversion I Green | >1 | 12.669(1.34) | 12.946(2.004) | 12.613(2.677) | 9.633(1.729) | 12.018(1.252) | 11.494(1.491) | 7.617(6.364) | 3.931(2.178) |
| Bisulfite Conversion I Red | >1 | 5.761(0.65) | 6.112(0.79) | 5.541(0.584) | 4.343(0.317) | 5.119(0.568) | 4.197(0.302) | 7.674(4.879) | 3.767(0.442) |
| Bisulfite Conversion II | >1 | 4.835(0.469) | 6.198(0.7) | 6.749(0.695) | 2.769(0.604) | 4.561(0.613) | 3.586(0.832) | 3.096(2.321) | 1.258(0.233) |
| Specificity I Green | >1 | 15.549(2.122) | 10.132(1.888) | 13.475(2.956) | 10.645(2.132) | 12.733(2.079) | 8.693(1.404) | 9.224(2.115) | 6.803(1.296) |

| | | | | | | | | | |
|---|---|---|---|---|---|---|---|---|---|
| Specificity I Red | >1 | 4.934(0.491) | 5.562(0.463) | 6.237(0.789) | 4.753(0.429) | 4.971(0.603) | 4.573(0.517) | 3.966(0.792) | 2.924(0.599) |
| Specificity II | >1 | 21.965(2.374) | 25.497(3.644) | 26.033(5.584) | 20.571(2.864) | 21.471(1.961) | 13.928(3.142) | 23.623(5.108) | 16.682(1.949) |
| Non-polymorphic Green | >5 | 10.724(0.839) | 8.402(0.679) | 8.722(0.788) | 10.33(0.82) | 10.425(0.877) | 8.369(0.829) | 15.067(2.85) | 9.379(0.917) |
| Non-polymorphic Red | >5 | 17.633(1.527) | 15.871(1.109) | 14.635(1.156) | 19.132(2.132) | 16.884(2.321) | 14.759(2.01) | 19.094(2.408) | 17.003(1.251) |
| (Methylated median intensity + Unmethylated median intensity)/2 | >10.5 | 11.99(0.143) | 12.18(0.184) | 12.659(0.191) | 11.852(0.137) | 12.093(0.144) | 12.213(0.238) | 10.901(0.307) | 11.165(0.191) |
| Percentage probes with detection p-value < 0.01 | >99% of total probes | 99.929(0.02) | 99.893(0.03) | 99.922(0.005) | 99.916(0.011) | 99.937(0.017) | 99.948(0.012) | 99.802(0.052) | 99.525(0.242) |
| Percentage probes with beadcount > 3 | >99% of total probes | 99.788(0.04) | 99.898(0.031) | 99.789(0.058) | 99.888(0.023) | 99.821(0.034) | 99.894(0.014) | 99.863(0.025) | 99.909(0.013) |
